# Charting the biosynthetic landscape of hybrid polyketide-nonribosomal peptide-specialized lipids

**DOI:** 10.1101/2025.10.24.682833

**Authors:** Fatima El Arnouki Belhaji, Dries De Ruysscher, Giel Vanreppelen, Laura-Lynn Huybrechts, Mohammad M. Alanjary, Mitja M. Zdouc, Emmanuel L.C. de los Santos, Odessa Van Goethem, Hans Gerstmans, Patrick Van Dijck, Marnix H. Medema, Angus Weir, Eveline Lescrinier, Joleen Masschelein

## Abstract

Polyunsaturated fatty acid (PUFA) synthase-like enzymes are best known for their role in membrane lipid biosynthesis in marine bacteria, but have also been repurposed for the assembly of specialized lipid metabolites with unique biological functions. Here, we illuminate their broader biosynthetic potential by charting the unexplored landscape of hybrid peptide-polyketide-specialized lipid biosynthesis in bacteria. Using a targeted genome mining strategy, we identified more than 60 biosynthetic gene clusters that combine PUFA synthase-like, polyketide synthase (PKS), and nonribosomal peptide synthetase (NRPS) enzymes across diverse bacterial lineages. Comparative analysis revealed extensive diversification of these triple hybrid pathways through gene fusion, domain reshuffling and recruitment of accessory enzymes. We further expand the known repertoire of peptide-polyketide-specialized lipid hybrids by identifying the chitinimines, a new family of amphiphilic metabolites produced by *Chitinimonas koreensis* featuring a C22 polyunsaturated lipid chain conjugated to a cyclic peptide-polyketide and a pyruvate-derived cyclic acetal moiety. The chitinimines exhibit surfactant properties, as well as moderate antibacterial activity against Gram-positive bacteria and contribute to a growth-promoting interaction between *C. koreensis* and *Salmonella* spp. Together, these findings demonstrate that PUFA synthase-like systems are far more versatile than previously appreciated, playing a key role in combinatorial biosynthetic innovation and serving as a rich, untapped source of chemically and functionally diverse specialized lipids.

## INTRODUCTION

Polyunsaturated fatty acids (PUFAs), such as eicosapentaenoic acid (EPA), docosahexaenoic acid (DHA) and arachidonic acid (AA), are essential components of cell membranes and serve as precursors for diverse signalling molecules.^[1-3]^ They are well known for their beneficial effects on human health, influencing various physiological and biochemical processes.^[2,3]^ PUFAs are synthesized via two distinct metabolic pathways. The first pathway is commonly found in eukaryotic cells and involves the elongation and oxygen-dependent desaturation of existing fatty acids.^[4-6]^ Prokaryotic microorganisms, on the other hand, such as marine psychrophilic γ-proteobacteria, are capable of synthesizing PUFAs *de novo* from simple carboxylic acid building blocks. In these bacteria, PUFAs play a key role in maintaining cell membrane fluidity and offer protection against reactive oxygen species.^[1]^ Their biosynthesis is directed by the *pfaABCD* gene cluster, which encodes an iterative type I fatty acid synthase (FAS)/polyketide synthase (PKS) multienzyme complex (**Fig. 1A**).^[7]^ Each Pfa enzyme is composed of a set of discrete catalytic domains responsible for the sequential incorporation, elongation and modification of malonyl-CoA building blocks. Canonical *pfa*A genes encode a ketosynthase (KS), an acyltransferase (AT), four to six acyl carrier proteins (ACPs), a ketoreductase (KR) and a dehydratase (DH) domain. PfaB is monofunctional, harboring a single AT domain, while *pfaC* encodes a KS, a chain-length factor (CLF) and two dehydratase/isomerase (DH/I) domains. The final gene, *pfaD*, encodes an enoyl reductase (ER) domain. Throughout the assembly process, all acyl intermediates remain covalently bound to the ACP domains as thioesters. Although the *pfaA-D* operon is generally conserved among PUFA-producing bacteria, its genetic organization and domain architecture can vary depending on the producing species and the type of PUFA that is assembled.^[1,8]^ Despite recent progress, the exact mechanisms underlying the biosynthesis of PUFAs in bacteria are still not fully understood.^[9-11]^

**Figure 1.**
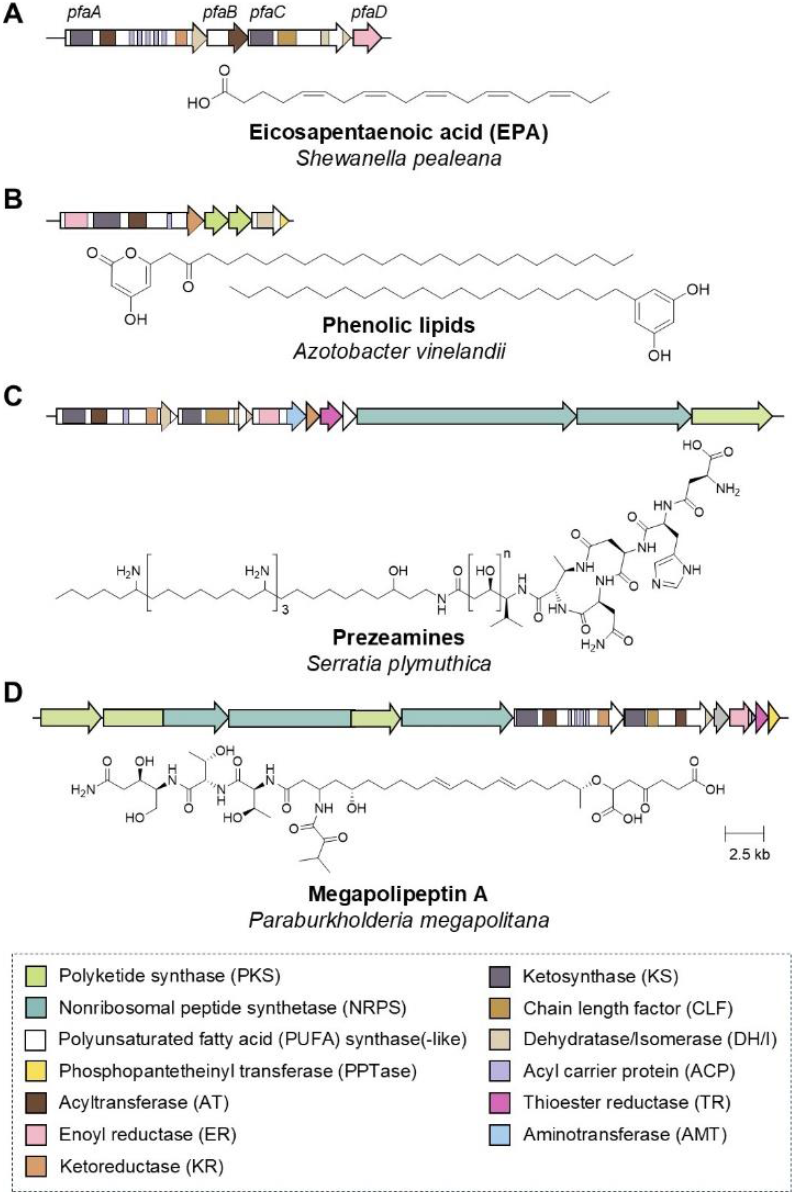
Genetic and structural diversity of polyunsaturated fatty acid (PUFA)-derived natural products. Biosynthetic gene cluster and chemical structures of A) the PUFA eicosapentaenoic acid, B) phenolic lipid-type secondary lipids and the C) prezeamine and D) megapolipeptin hybrid polyketide-nonribosomal peptide-secondary lipid metabolites.

Interestingly, a large-scale bioinformatic analysis of sequenced bacterial genomes has revealed that PUFA synthase-like gene clusters are more widespread than initially anticipated.^[12]^ They occur in bacteria from diverse ecological niches and are predicted to assemble a wide range of long-chain (>C20) lipid products with specialized functions. These FAS/PKS biosynthetic pathways coexist with primary fatty acid metabolism and are referred to as secondary lipid synthases. So far, only a handful of secondary, or specialized, lipids have been identified.^[12]^ Known examples include the C26-C32 alkyl chains of heterocyst glycolipids in nitrogen-fixing cyanobacteria and the C22-C26 phenolic lipid alkyl chains in cysts of *Azotobacter vinelandii* (**Fig. 1B**).^[13-15]^ Notably, in *A. vinelandii*, the PUFA synthase-like genes collaborate with two flanking type III PKS genes that direct the biosynthesis of the phenolic head group.

The integration of PUFA synthase-like pathways with other types of biosynthetic machinery is a powerful strategy to expand the structural and functional diversity of specialized lipids. The most striking example of such combinatorial biosynthesis is found in the zeamine pathway, which combines nonribosomal peptide, type I modular polyketide and PUFA synthase-like biosynthetic machinery.^[16,17]^ The zeamines are a family of broad-spectrum antibiotics produced by phytopathogenic *Serratia* and *Dickeya* spp. that contain variable peptide-polyketide moieties linked to a common 40-carbon pentaamino-hydroxyalkyl chain (**Fig. 1C**).^[16,18,19]^ This unusual polyamino alcohol chain is assembled by a unique secondary lipid synthase that has recruited several additional catalytic domains, including an aminotransferase (AMT), KR and thioester reductase (TR) domain.^[16,17]^ In a parallel pathway, hexapeptide-mono- and diketide thioesters are generated by a hybrid PKS-NRPS and subsequently condensed to the 40-carbon alkyl chain.^[17]^ The resulting zeamines thus represent a novel class of ‘triple hybrid’ bioactive specialized lipid metabolites. Rather than serving as membrane constituents, they are secreted as antibiotics to fight and eliminate a broad range of organisms, including bacteria, fungi, oomycetes, nematodes and plants.^[19-21]^ This broad-spectrum antagonistic activity stems from the cationic amphiphilic nature of the long polyamino alcohol chain, which interacts directly with cellular membranes and disrupts their integrity.^[22]^

Several structurally related metabolites, named fabclavines, with minor differences in the length of the polyamino and polyketide moieties, and in the amino acid composition of the hexapeptide, have been discovered in entomopathogenic *Xenorhabdus* and *Photorhabdus* bacteria that live in mutualistic symbiosis with insect-infecting nematodes. Like the zeamines, fabclavines exhibit potent antibacterial, antifungal and antiprotozoal activity. They are believed to be produced to help kill the insect host and protect its cadaver against potential food competitors.^[23]^ Very recently, a third example of a nonribosomal peptide-polyketide-specialized lipid hybrid was uncovered through genome mining of a *Burkholderiales* strain collection.^[24]^ Heterologous expression of a silent biosynthetic gene cluster (BGC) from *Paraburkholderia megapolitana* led to the discovery of megapolipeptins A and B. These metabolites are composed of a C16 or C18 partially unsaturated fatty acid coupled to a 4-oxoheptanedioic moiety and the product of a heptamodular PKS-NPRS (**Fig. 1D**). Overall, the zeamine, fabclavine and megapolipeptin pathways are a testament to Nature’s remarkable ability to mix and match different types of biosynthetic machinery to access novel chemical scaffolds with useful biological properties. Their discovery raises the intriguing question of how widespread such hybrid specialized lipid metabolites are and whether additional examples remain to be discovered.

In this work, we chart the biosynthetic landscape of hybrid peptide-polyketide-specialized lipid metabolites by conducting a computational search for gene clusters encoding combinations of PKS, NRPS and PUFA synthase-like enzymes across a large, dereplicated collection of high-quality prokaryotic genomes. We uncover various additional examples of such clusters in diverse bacterial lineages, providing, for the first time, a comprehensive view of the distribution, diversity and evolution of these triple hybrid BGCs, as well as new insights into the recruitment of unconventional catalytic domains within the PUFA synthase-like multienzyme complexes. Furthermore, we expand the limited catalogue of known secondary lipid-containing natural products by demonstrating that one of the newly identified clusters in *Chitinimonas koreensis* directs the biosynthesis of a novel family of hybrid metabolites, which we name chitinimines. The chitinimines were isolated, structurally characterized and found to be essential for the growth-promoting activity of *C. koreensis* towards *Salmonella* spp., while also exhibiting surfactant properties and moderate antibacterial activity against Gram-positive bacteria. Detailed sequence analysis of the proteins encoded within the chitinimine cluster enabled us to propose a plausible biosynthetic pathway, raising intriguing questions about the catalytic activity and biosynthetic programming of the PUFA synthase-like enzymes involved.

## RESULTS & DISCUSSION

### Diversity and distribution of hybrid PKS-NRPS-PUFA synthase-like biosynthetic pathways

To explore the distribution of hybrid polyketide-nonribosomal peptide-specialized lipid biosynthesis in bacteria, we designed a custom set of profile Hidden Markov Models (pHMMs) to search the antiSMASH database (v4) for gene clusters containing genes encoding all three types of biosynthetic machinery within 20 kb of each other (**Figure 2A**). This search yielded 87 candidate (proto)clusters in bacteria from diverse taxonomic lineages (**Table S1**). Following manual curation, we discarded 24 clusters either because the predicted PKS gene was a misannotated *pfaBC* homolog, the co-localization and synteny of the three types of biosynthetic genes were not conserved among closely-related strains, the *pfa*-like and the PKS-NRPS genes were separated by many intervening genes with no clear operon-like organization, and/or because a subset of the biosynthetic genes were highly similar in sequence to a known natural product BGC listed in the MIBiG database.^[25]^

**Figure 2.**
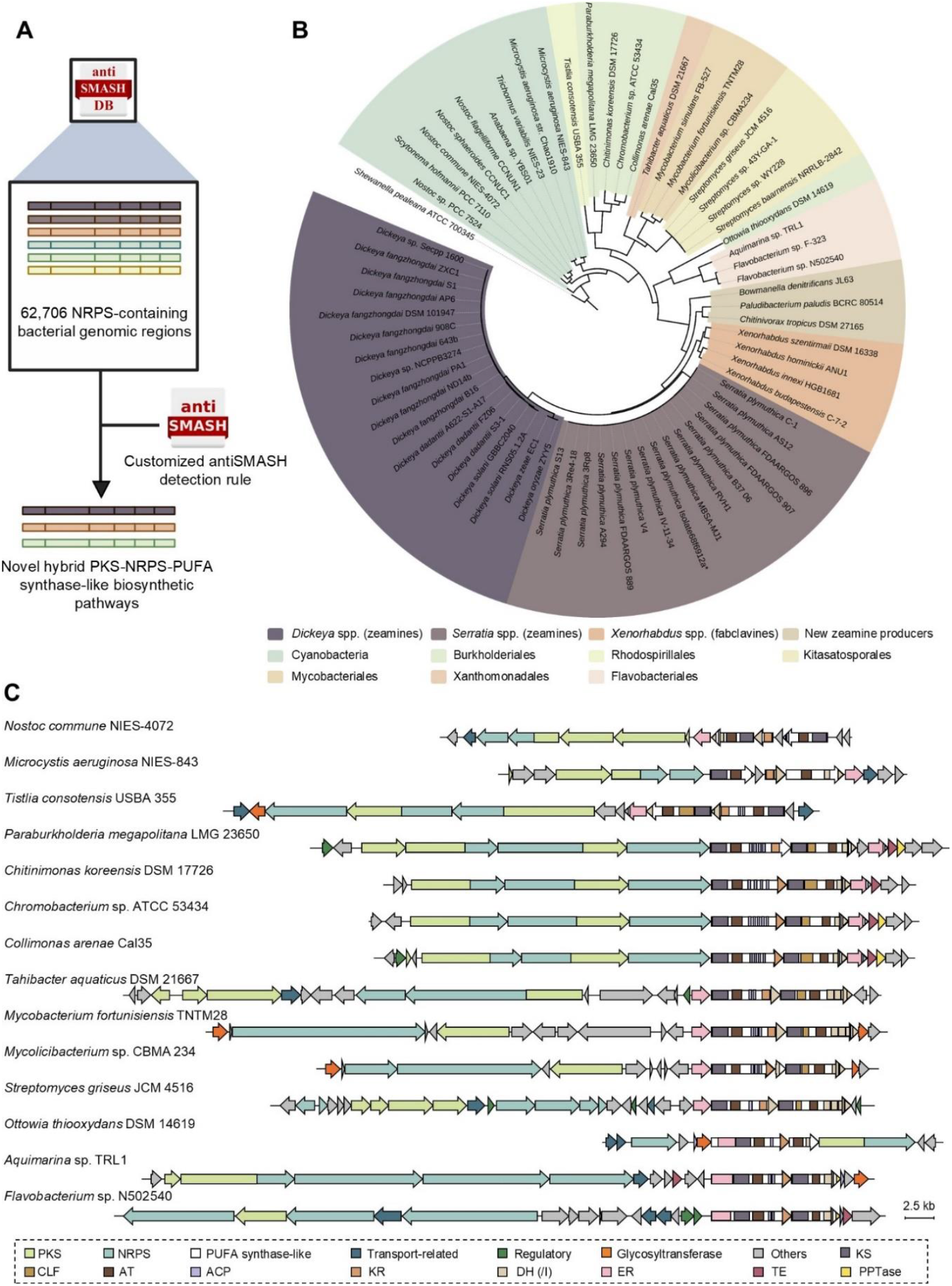
Computational search for gene clusters harboring PKS and NRPS genes, as well as genes encoding PUFA synthase-like biosynthetic machinery. A) Genome mining strategy used in this study. NRPS-encoding bacterial genomic regions from the antiSMASH database were screened with a customized search rule to detect candidate hybrid PKS-NRPS-PUFA synthase-like pathways. B) Phylogenetic analysis of KS domains from PfaA homologs encoded in the hybrid pathways. A neighbor-joining tree was constructed from the KS domain sequences, with the PfaA KS domain from the *S. pealeana* PUFA synthase as the outgroup. Colored clades correspond to related pathways and producing organisms. C) Representative examples of triple hybrid BGCs identified. The three biosynthetic classes are highlighted in different colors, and the predicted domain organization of the Pfa homologs is shown.

The remaining 63 clusters are broadly distributed among both Gram-positive and Gram-negative bacteria from diverse ecological niches, consistent with previous observations that *pfa*-like genes are widespread across multiple bacterial lineages.^[12]^ To investigate their diversity and evolutionary relationships, we constructed a phylogenetic tree based on the protein sequences of the KS domains encoded within the *pfaA* homologs in each cluster (**Figure 2B**). Given that the phylogeny of CLF domains has been shown to correlate closely with the carbon chain length and overall chemical scaffold of their polyketide products, a phylogeny of KS-CLF pairs (where present) was also constructed, which revealed a similar clade pattern (**Figure S1**).^[26]^

The largest clade, comprising 40 clusters, corresponds primarily to *Serratia* and *Dickeya* species known to produce the zeamine antibiotics, as well as *Xenorhabdus* species that synthesize the structurally related fabclavines (**Figure 2B**). Interestingly, this clade also includes highly similar BGCs from three strains that have not previously been reported to produce zeamines or fabclavines: *Chitinivorax tropicus* DSM 27165, *Paludibacterium paludis* BCRC 80514 and *Bowmanella denitrificans* JL63. Based on the architecture of the BGCs and the predicted substrate specificity of the adenylation (A) domains encoded by the NRPS genes, we hypothesize that these strains produce zeamine-like, rather than fabclavine-like metabolites (**Figure S2, Table S2**). Comparative gene cluster analysis further revealed some noticeable species-specific differences in the genomic regions flanking the zeamine BGC in *Dickeya* and *Serratia* strains. In *Dickeya*, the clusters are typically adjacent to Resistance-Nodulation-Division (RND) family transporter genes, whereas in *Serratia*, they are flanked by a pyrrolnitrin biosynthetic operon (**Figure S2**). The genomic insertion site of the zeamine genes can be clearly identified by comparing the genome of *S. plymuthica* RVH1, which harbors the zeamine BGC, with that of the closely related strain UBCF_13, which does not. Extensive synteny up- and downstream of the insertion site indicates that the zeamine genomic island is integrated between the pyrrolnitrin operon and the *hyc* and *hyp* loci involved in the assembly and maturation of the formate hydrogenlyase complex (**Figure S3**).

The second largest clade comprises a group of uncharacterized triple hybrid pathways found in cyanobacteria belonging to the *Microcystis, Trichormus, Anabaena, Scytonema* and *Nostoc* genera (**Figure 2B**). Filamentous nitrogen-fixing cyanobacteria are known to use PUFA synthase-like machinery to assemble heterocyst glycolipids: specialized lipids in the cell envelope of heterocysts that limit oxygen diffusion and thereby protect nitrogenase enzymes from inactivation.^[13,14]^ The strains within this clade, however, do not have the genetic capacity to produce these glycolipids, but harbor a distinct genomic region in which *pfa*-like genes are directly flanked by genes encoding a hybrid PKS-NRPS (**Figure 2C**). The *pfa*-like genes exhibit several atypical features. Most notably, the *pfaA* homolog is split into two separate genes that are separated by a gene encoding a protein of unknown function with a predicted NAD(P)-binding Rossmann fold (**Figure S4**). The *pfaB* and *pfaC* homologs are fused into single genes encoding an AT and two DH/I domains. Interestingly, all *pfaBC* genes in this clade lack a CLF domain and in the *Microcystis aeruginosa* strains, the PfaB KS domain is also absent. The hybrid PKS-NRPS assembly lines are mostly composed of two PKS modules and one NRPS module. In *Nostoc commune* NIES-4072, *Scytonema hofmannii* PCC7110, *Nostoc flagelliforme* CCNUN1 and *Nostoc sphaeroides* CCNUC1, an additional PKS module is present. In the *M. aeruginosa* NIES-843 and Chao 1910 assembly lines, this third module is absent and replaced by one or two genes encoding a putative tryptophan halogenase.

Another well-defined clade of triple hybrid biosynthetic pathways is found in several Burkholderiales bacteria, such as *Collimonas arenae* Cal35, *Chromobacterium* sp. ATCC 53434, *Chitinimonas koreensis* DSM 17726 and *Paraburkholderia megapolitana* LMG 23650, which produces the megapolipeptins (**Figure 1D, 2B and 2C)**.^[24]^ The gene cluster from *C. arenae* was recently reported to be involved in the biosynthesis of the carenaemins, a new family of structurally undefined metabolites with potent antifungal and plant-protective bioactivities.^[27]^ The *pfa*-like genes in this clade are highly conserved and share some remarkable features (**Figure S5**). Most notably, the *P. megapolitana* PUFA synthase-like genetic machinery encodes an unusual stand-alone cupin-family enzyme, which is predicted to act as a hydroxylase, and the PfaD homologs in this clade harbor an additional ACP domain.^[24]^ A downstream TE domain-encoding gene is followed by a *pfaE*-like phosphopantetheinyl transferase (PPTase) gene in all members of this clade except *C. koreensis*, where the PPTase is presumably encoded elsewhere in the genome, as also observed in the PUFA producer *Moritella marina*.^[28]^ Several genes encoding putative tailoring enzymes have also been recruited, including a conserved polysaccharide pyruvyl transferase. In *P. megapolitana*, this enzyme is thought to act in concert with a 2-succinyl-5-enolpyruvyl-6-hydroxy-3-cyclohexene-1-carboxylic acid (SEPHCHC) synthase to assemble the 4-oxoheptanedioic moiety that is appended to the long-chain unsaturated fatty acid (**Figure 1D**). The hybrid PKS-NRPS assembly lines in this clade contain at least five NRPS and two PKS modules, with *P. megapolitana* encoding an additional PKS and NRPS module. Interestingly, a highly similar pathway is also present in *Tistlia consotensis* USBA 355, an α-proteobacterium from the Rhodospirillales order. Its *pfa* genes have the same overall architecture as those in the Burkholderiales members, but the *pfaE* PPTase and pyruvyl transferase are missing, while the SEPHCHC synthase is retained. The module and domain organization of the PKS-NRPS closely resembles that of *P. megapolitana*, the only difference being the presence of a C-MT domain in the first PKS module (**Figure S5**).

Another member of the Burkholderiales order, *Ottowia thiooxydans* DSM 14619, harbors a markedly distinct hybrid PKS-NRPS-specialized lipid pathway (**Figure 2C**). In this cluster, the *pfaD* and *pfaA* homologs are fused into a single gene, and a fused *pfaB*C homolog lacks the canonical KS-CLF heterodimer (**Figure S6**). These pfa-like genes are directly followed by a trimodular hybrid PKS-NRPS gene. Upstream, the cluster contains a second NRPS gene of the non-α-polyamino acid (NAPAA) type, encoding a single NRPS module with an A and PCP domain, as well as a highly atypical C-terminal Pls/PosA-like domain previously reported in δ-poly-L-ornithine and ε-poly-L-lysine synth(et)ases.^[29,30]^ Aside from this core biosynthetic machinery, the cluster encodes a glycosyltransferase and a protein that is not similar in sequence to any proteins of known function.

Our genome mining search further revealed that hybrid PKS-NRPS-PUFA synthase-like clusters are also distributed among Gram-positive bacteria, including *Mycobacterium fortunisiensis* TNTM28 and *Mycolicibacterium* sp. CBMA 234 (**Figure 2B and 2C**). These clusters share similar *pfa*-like genes, a monomodular PKS and a coenzyme A ligase (CAL) gene, as well as a gene encoding an NRPS module with a C-terminal amino acid adenylation domain-containing protein (TIGR01720). While little is known about this domain, the Conserved Domain Database (CDD) suggests a possible role in post-condensation events.^[31]^ Remarkably, the fused *pfaBC* homolog within the *M. fortunisiensis* cluster encodes no less than four DH/I domains (**Figure S7**). Compared to *M. fortunisiensis*, the *Mycolicibacterium* sp. cluster contains four additional NRPS modules, one of which also harbors a second TIGR01720 domain. In both organisms, the PKS and NRPS genes are separated from each other by a gene encoding a putative isoprenylcysteine carboxylmethyltransferase (ICTM) family protein. The region between the PKS-NRPS genes and the *pfa*-like genes shows limited conservation, with predicted protein functions ranging from HNH endonucleases and Ig-like domain-containing proteins to PE (Pro– Glu) family proteins. Although technically not a triple hybrid cluster, a related BGC is present in *M. simulans* FB-527 (**Figure S7**). This cluster encodes a highly similar PUFA synthase-like pathway together with a bimodular NRPS terminating in a TIGR01720 domain, but lacks the PKS genes. Intriguingly, phylogenetic analysis also uncovered a highly similar cluster in the Gram-negative strain *Tahibacter aquaticus* DSM 21667 (**Figure 2B and S7**). Here, the *pfa*-like and PKS genes display a very similar architecture to those of the Mycobacteriales, but the CAL domain is replaced by a KS-ACP didomain. This cluster further encodes an additional PKS module followed by four NRPS modules, none of which contain a TIGR01720 domain or are flanked by an ICTM homolog.

Other Gram-positive bacteria that harbor triple hybrid pathways include several *Streptomyces* species in which PUFA synthase-like and PKS machinery appear to collaborate with NRPS enzymes to direct the biosynthesis of a metallophore-type metabolite (**Figure 2C**). Nearly identical clusters were detected in *S. griseus* JCM 4516, *Streptomyces* sp. 43Y-GA-1, *S. baarnensis* NRRL B-2842 and *Streptomyces* sp. WY228. In each case, the *pfa*-like genes comprise a *pfaD*, a *pfaA*, and a fused *pfaB-pfaC*-like homolog that stands out by encoding four DH/I domains (**Figure S8**). The PKS and NPRS assembly lines each comprise three modules. Notably, the clusters also harbor several tailoring enzymes, including a salicylate synthase, a SAM-dependent methyltransferase, a metallopeptidase, and a Rossmann fold-containing oxidoreductase, consistent with the predicted metal-binding function of the metabolic product of these BGCs.

A final well-defined clade is formed by hybrid PKS-NRPS-PUFA synthase-like pathways in two Flavobacteriales members: *Aquimarina* sp. TRL1 and *Flavobacterium* sp. N502540 (**Figure 2C**). Several members of the Flavobacteriales are known PUFA producers, including species of the *Flavobacterium, Flexibacter* and *Psychroflexus* genera. These are often psychrophilic, halophilic and/or piezophilic bacteria that rely on PUFA production to maintain membrane integrity and fluidity under harsh environmental conditions.^[32-34]^ The *pfa*-like genes in the triple hybrid clusters from *Aquimarina* sp. TRL1 and *Flavobacterium* sp. N502540 have a highly similar organization, containing fused *pfaDA* and *pfaBC* homologs, the latter also harboring a PPTase domain (**Figure S9**). This unusual arrangement closely resembles that of a gene cluster encoding an uncharacterized secondary lipid synthase in *Renibacterium salmoninarum* ATCC 33209.^[12]^ The PKS-NRPS assembly lines of the two strains differ in complexity. In *Aquimarina* sp. TRL1, the cluster encodes two PKS and nine NRPS modules, two of which terminate in a TIGR07120 domain. In *Flavobacterium* sp. N502540, the cluster instead harbors a single PKS module, embedded between eleven NRPS modules, one of which harbors a TIGR01720 domain, while another has an unusual A domain with integrated oxidase activity (A–Ox). In both pathways, the *pfa*-like operon is separated from the PKS-NRPS genes by a variable set of intervening genes predicted to encode regulatory proteins, transporters, oxidoreductases, ATPases and endopeptidases. Interestingly, a related *pfa*-like operon located adjacent to a large NRPS gene cluster, but lacking PKS genes, is also present in *Flavobacterium* sp. F-323 (**Figure S9**).

Beyond its taxonomic and architectural diversity, this biosynthetic landscape also offers a unique view on how *pfa*-like enzymes within such triple hybrid pathways have evolved through domain reshuffling, gene fusion and the recruitment of diverse accessory enzymes and catalytic domains to expand the structural diversity of their secondary lipid products (**Figure S10**). Prominent examples of the latter include the pyridoxal phosphate (PLP)-dependent AMT domains and stand-alone KR and TR enzymes in the zeamine and fabclavine pathways, which contribute to the formation of the long polyamino alcohol chain in these metabolites. Similarly, a cupin-like domain is incorporated in the megapolipeptin secondary lipid synthase, whereas the cyanobacterial Pfa-like pathway from *T. variabilis* NIES-23 features a protein with unknown function harbouring an NAD(P)-binding Rossmann fold. The most striking example of such catalytic diversification is found in the hybrid NRPS-PUFA synthase-like pathway in *Nostoc* sp. PCC 7524, which was detected in our initial genome mining search but excluded during manual curation because the *pfaB* homolog had been misannotated as a PKS gene (**Figure 2B**). In this cluster, the *pfa*-like operon has acquired genes encoding two FAD-dependent oxidoreductases, a putative hydrolase, and even an entire additional NRPS module. Elucidating how these enzymes act in concert with the PUFA synthase-like machinery may uncover new routes to structural diversification in bacterial lipids.

### Purification and structure elucidation of the chitinimines

To validate our genome mining strategy, we selected the hybrid PKS-NRPS-PUFA synthase-like gene cluster from *Chitinimonas koreensis* DSM 17726 for in-depth characterization (**Figure 2C**). The cluster comprises seven core biosynthetic genes (*chtnA-G*), which are flanked by several genes encoding putative tailoring enzymes and regulation-related proteins (**Table S3**). To identify the metabolic product of this cryptic gene cluster, we inactivated the *chtnA* gene by insertional mutagenesis. UHPLC-ESI-Q-TOF-MS analysis of ethyl acetate extracts from cultures of *C. koreensis*, grown for four days in minimal medium containing glucose as a sole carbon source, identified two metabolites whose production was abolished in the Δ*chtnA* mutant: a metabolite with the molecular formula C_49_H_80_N_6_O_11_ (calculated for C_49_H_81_N_6_O_11+_: 929.5957, found: 929.5968), and one with the molecular formula C_48_H_78_N_6_O_11_ (calculated for C_48_H_79_N_6_O_11+_: 915.5801, found: 915.5827), which we named chitinimine I and II, respectively (**Figure 3A**). To identify these metabolites, large-scale cultures of *C. koreensis* DSM 17726 were grown on minimal medium and ethyl acetate extracts of the agar plates were fractionated by preparative HPLC (**Figure S11**).

**Figure 3.**
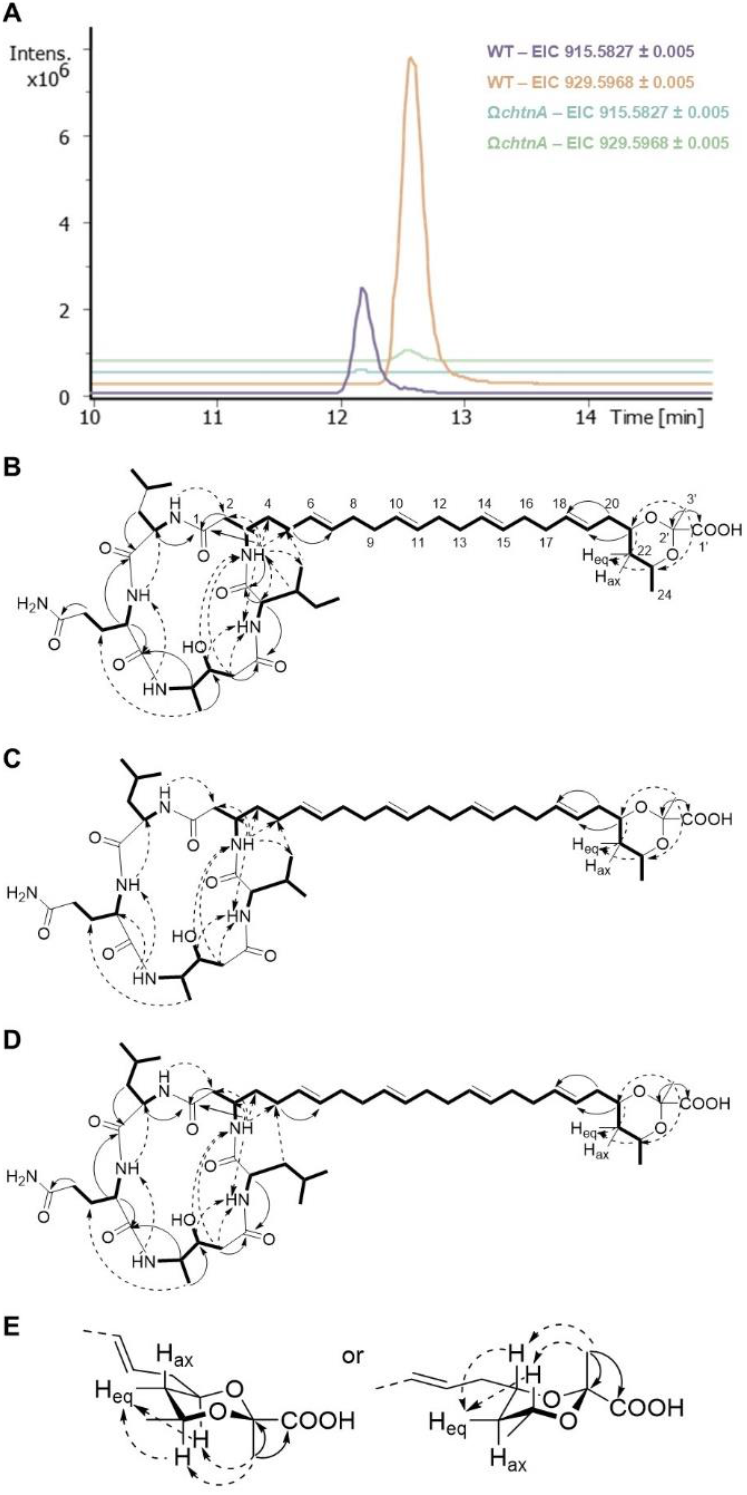
Identification and structure elucidation of the chitinimines. A) Insertional mutagenesis abolishes chitinimine production by *C. koreensis* DSM 17726. Extracted ion chromatograms (EICs) at *m/z* 915.5827 ± 0.005 and 929.5968 ± 0.005 (corresponding to the [M + H]^+^ ion of chitinimine II and I/III, respectively) from LC-MS analyses of crude extracts from agar-grown cultures of wildtype *C. koreensis* (purple and orange) and the insertional mutant *C. koreensis* Ω*chtnA* (blue and green). B-D) TOCSY (bold lines) and key ^1^H-^13^C HMBC (full arrows) and NOESY (dashed arrows) correlations observed for chitinimine I (B), chitinimine II (C) and chitinimine III (D). E) Two distinct conformations of the cyclic acetal are compatible with the observed HMBC and NOESY correlations.

The planar structure of the purified metabolites was elucidated using a combination of [^1^H, ^1^H] COSY, [^1^H, ^1^H]-TOCSY, [^1^H, ^1^H]-NOESY, [^1^H, ^13^C]-HSQC and [^1^H, ^13^C]-HMBC experiments (**Figures 3B** and **S12**-**S21, Tables S4**-**S6**). In the COSY spectrum of chitinimine I, four resonances with chemical shifts characteristic of amino acid Cα protons (δH 4.01, 4.03, 3.41, and 4.01) correlated with signals attributable to exchangeable N-H protons (δ_H_ 8.89, 8.58, 7.75, and 8.11, respectively). Further analysis of COSY and TOCSY spectra identified four amino acid spin systems, which, supported by HSQC and HMBC data, were assigned to leucine, glutamine, 4-amino-3-hydroxypentanoic acid (4A3HPA) and (iso)leucine in the sequence Leu-Gln-4A3HPA-Ile/Leu. The identity of these tetrapeptide fragments was further confirmed by LC-ESI-MS/MS analysis (**Figure S22**). Thus, the chitinimine I sample was found to be a mixture of two isomers, present in a 2:1 ratio, differing only by the incorporation of either Ile or Leu at a single position in the peptide moiety. For clarity, we designated these as chitinimine I and III, respectively. An unbroken network of COSY correlations further established the structure of the C1 to C24 fragment of chitinimine I and III, which corresponds to a long alkyl chain bearing E-configured double bonds between C6-C7, C10-C11, C14-15 and C18-19 (based on ^3^*J*_HH_ coupling constants of 15-16 Hz). Signals at 172.9, 45.7, 68.2 and 64.4 ppm in the ^13^C NMR spectrum led us to propose that the alkyl chain also has a carbonyl group at C1, an amine group at C3 and oxygen substitutions at C21 and C23, respectively. A ^2^*J*_CH_ correlation between the N-H proton (δ_H_ 8.17) of this amine group and the carbonyl carbon of the Ile/Leu residue (δ_C_ 170.5), as well as an HMBC correlation between the Cα-H proton (δ_H_ 4.01) of Leu and the C1 carbonyl carbon (δ_C_ 172.9) indicate that the peptide is appended to C1 of the alkyl chain and forms a macrolactam ring via condensation with the C3 amine group. Finally, signals with chemical shift values corresponding to a hydroxylated pyruvate moiety were observed in the ^1^H, ^13^C and HMBC spectra.^[35]^ NOESY correlations between the pyruvate methyl group and both H-21 and H-23, and between H-21 and H-23 and the equatorial proton at C22 indicate that the pyruvate is condensed to the C21 and C23 oxygen substituents, forming a six-membered cyclic acetal that likely adopts one of two possible chair conformations (**Figure 3B-E**). The bridging position of this pyruvate moiety was further confirmed by LC-ESI-MS/MS analysis (**Figure S22**) and acid hydrolysis (**Figure S23**).

Comparison with the NMR spectroscopic data of chitinimine I and III showed that chitinimine II has an identical structure, except for the presence of Val in place of the Ile/Leu residue, consistent with the difference of a CH_2_ group in the molecular formula. The absolute stereochemistry of the amino acid residues in chitinimine I-III was determined by acid hydrolysis and derivatization using Marfey’s reagent. UHPLC-ESI-Q-TOF-MS comparisons with Marfey’s derivatives of the appropriate L- and D-amino acid standards revealed that the Gln and 4A3HPA residues are L-configured, while Leu, Ile and Val were detected in both the L- and D-configuration (**Figure S24**). To distinguish between these stereoisomers, we renamed chitinimines I, II and III as Ia/b, IIa/b and IIIa/b, depending on whether they contained L- or D-Ile/Val/Leu, respectively. Notably, Gln was found to be unstable under the hydrolysis conditions and was rapidly converted into Glu during the treatment with concentrated HCl and heat. Its stereochemical assignment was therefore based on comparison with Marfey’s derivatives of L- and D-Glu standards.

### Analysis of the chitinimine biosynthetic gene cluster

Detailed bioinformatic analysis of the chitinimine BGC enabled us to propose a plausible pathway for their biosynthesis. Like the zeamine, fabclavine and megapolipeptin pathways, the chitinimine BGC encodes two parallel assembly lines: a hybrid PKS-NRPS (ChtnA-C) and a PUFA synthase-like multienzyme complex (ChtnD-G) (**Figure 4**). ChtnD-G share a high degree of sequence similarity with the canonical PUFA biosynthetic enzymes PfaA, PfaBC and PfaD, respectively, differing only in the number of ACP domains and in the presence of an additional ACP domain in ChtnF and a stand-alone thioesterase (TE) (ChtnG). Based on the biosynthetic logic of related secondary lipid synthases, we propose that ChtnD-G act iteratively to assemble the C22 alkyl chain that constitutes the lipid scaffold of the chitinimines. The assembly process is likely initiated by the loading of a malonyl unit, followed by 10 cycles of chain elongation with varying degrees of β-carbon processing. This hypothesis is supported by analysis of conserved sequence motifs, which indicate that the acyl transferase (AT) domains within ChtnD and ChtnE have specificity for malonyl-CoA (**Figure S25**). The use of malonyl-CoA as both starter and extender unit aligns well with observations from *in vitro* PUFA biosynthetic studies.^[36]^

**Figure 4.**
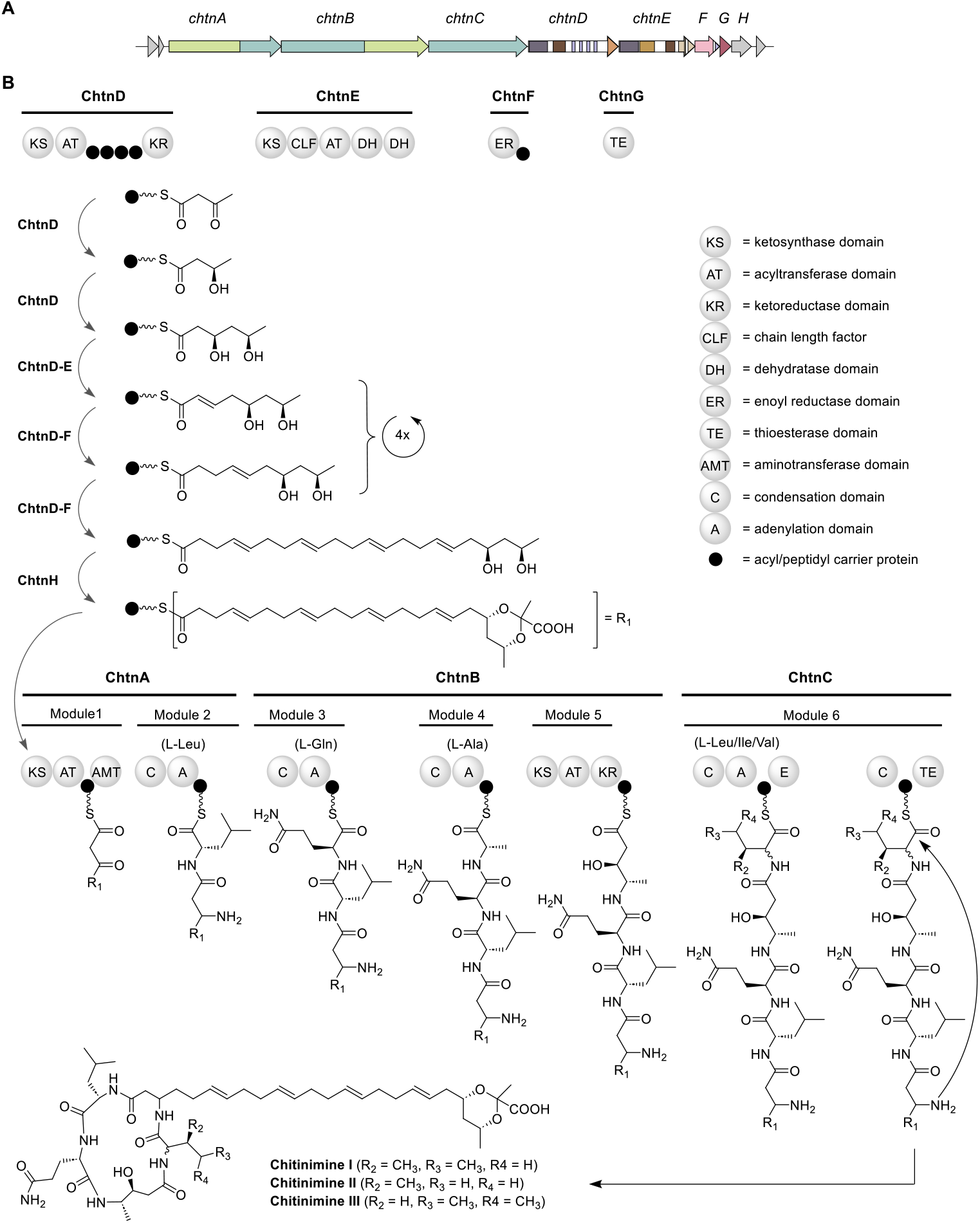
Organization of the chitinimine BGC and proposed pathway for chitinimine biosynthesis. A) ChtnA-H direct the biosynthesis of chitinimines in *C. koreensis* DSM 17726. The NRPS and hybrid PKS/NPRS genes are highlighted in dark and light green, respectively. The PUFA synthase-like genes are marked in white with their individual domains colored according to the legend in Figure 2. The gene encoding the pyruvyl transferase (*chtnH*) is shown in grey. B) The proposed roles played by ChtnA-H in chitinimine biosynthesis. The proposed structures of the ACP- and PCP-bound thioester intermediates following β-carbon processing are presented. For clarity, only one chair conformations of the cyclic acetal is shown.

In the first two chain extension cycles, the β-keto groups in the elongated intermediates are reduced to generate β-hydroxythioesters, but no further dehydration takes place. Sequence comparisons with KR domains of known stereospecificity from other PKSs suggest that the KR domain in ChtnD is a B-type KR domain (**Figure S26**), allowing the configuration of the C21 and C23 stereocenters, which could not be elucidated experimentally, to be tentatively assigned as R. During the following eight elongation cycles, the β-ketothioester intermediates are alternately converted into either α,β-unsaturated or -fully saturated thioesters by the ChtnE DH and ChtnF ER domains, respectively. Notably, the second DH domain within ChtnE lacks the active site Asp residue and has a truncated HxxxGxxxxP motif, suggesting it may be catalytically inactive (**Figure S27**). The successive condensation reactions are presumed to be catalysed by the KS domains within ChtnD and ChtnE, each of which contains the conserved CHH catalytic triad, except for the second KS domain of ChtnE, which is predicted to function as a CLF. Whether the ChtnD KS domain catalyzes chain elongation during the early stages of assembly and the KS-CLF dimer acts in the later cycles, as seen in PUFA synthases, remains to be investigated (**Figure S28**).^[11]^ Throughout the assembly of the alkyl chain, the biosynthetic intermediates are tethered to the ACP domains within ChtnD. Sequence analysis of the ACP domains confirmed that they all harbor the conserved Ser residue required for post-translational attachment of the phosphopantetheinyl prosthetic arm (**Figure S29**). The presence of variable numbers of tandem ACP domains in PfaA homologs is well-documented and has been linked to increased biosynthetic productivity by enhancing intermediate flux and enabling parallel processing.^[37,38]^ Interestingly, the PfaD homolog ChtnF harbors an unusual ACP domain whose function is unclear. It is tempting to speculate that this domain may play a role in binding α,β-unsaturated thioester intermediates in selected cycles to facilitate enoyl reduction by the ER domain. At some point during or after assembly of the long alkyl chain, the two hydroxyl groups are condensed with a pyruvate moiety, forming a ring structure. This transformation is likely catalyzed by the putative pyruvyl transferase ChtnH. In the megapolipeptin pathway, a related pyruvyl transferase that shares 72.5% sequence similarity with ChtnH catalyzes a distinct esterification reaction in which a 4-oxoheptanedioic moiety is appended to the terminal ω-1 hydroxyl group.^[24]^

Rather than being released from the PUFA synthase-like assembly line, the fully assembled C22 acyl thioester is presumably transferred directly onto the active site Cys residue of the N-terminal KS domain of ChtnA for further PKS/NRPS-mediated chain extension. *ChtnG*, which encodes a stand-alone type II TE (TE_II_), likely has a proofreading function, hydrolysing aberrant acyl chains from the carrier proteins to maintain biosynthetic efficiency. While TE_II_s are also known to be capable of catalysing aminoacyl chain transfer between PCP domains on separate subunits, it remains to be investigated whether ChtnG also participates in transferring the fully assembled alkyl chain to ChtnA.^[39,40]^

The peptide moieties of the chitinimines are proposed to originate from the ChtnA-C hybrid assembly line, which comprises two PKS and four NRPS modules (**Figure 4**). The ChtnA PKS module is predicted to extend the C22 acyl thioester intermediate with another malonyl unit, consistent with the predicted substrate specificity of its AT domain (**Figure S25**). Based on the role of related aminotransferase (AMT) domains in the zeamine, mycosubtilin and microcystin, the AMT domain within ChtnA is then proposed to convert the resulting β-ketothioester intermediate to the corresponding β-aminothioester. The next three NRPS modules incorporate L-Leu, L-Gln, and L-Ala, respectively (**Table S7**). This is consistent with the predicted specificity of the adenylation (A) domains and also aligns well with phylogenetic analysis of the condensation (C) domains, which place the ChtnA C domain in the hybrid group, typically found directly downstream of PKS modules, while the ChtnB C domains cluster with ^L^C_L_-type domains known to catalyse peptide bond formation between L-configured amino acyl thioesters attached to up- and downstream PCP domains (**Figure S30-31, Table S8**). The tripeptidyl acyl thioester assembled by the first four modules in ChtnA-B is then proposed to undergo another round of PKS-mediated two-carbon chain elongation with concomitant reduction of the β-keto group by the KR domain in ChtnB. Comparative sequence analysis predicts the S configuration for the resulting hydroxyl group, matching the results from the Marfey’s analysis (**Figure S24D**). The final NRPS module activates and incorporates either Ile (chitinimine I), Val (chitinimine II) or Leu (chitinimine III), indicating that the ChtnC A domain exhibits relaxed substrate specificity (**Table S7**). The presence of an epimerization (E) domain in this module is consistent with the observed D-configuration of Ile, Leu and Val in some chitinimine variants (**Figure S24 and S30**) and aligns with our phylogenetic analysis, which places the downstream C domain in a clade along with other C domains that accept D-configured amino acids (^D^C_L_ domains) (**Figure S31**). Although this C domain possesses the conserved active site motif HHxxxDG, it is not predicted to catalyse peptide bond formation. In the final step, the mature chitinimines I-III are proposed to be released from the assembly line via macrolactamization catalyzed by the C-terminal TE domain.

Beyond the core biosynthetic genes, putative functions were also assigned to the genes flanking *chtnA-G* based on sequence analyses (**Table S3**). In addition to the predicted pyruvyl transferase ChtnH, several other genes encode proteins with sequence similarity to monooxygenases (*F559_RS0116890*), transcriptional regulators (*F559_RS0116880*), and transport-related proteins (*F559_RS0116835, F559_RS0116840, F559_RS0116855, F559_RS0116860, F559_RS0116865, F559_RS0116875*), which may play a role in tailoring, export or regulation of the chitinimines.

### Biological activity of the chitinimines

To investigate the antimicrobial properties of the chitinimines, we first compared the activity of *C. koreensis* DSM 17726 wildtype and the Δ*chtnA* mutant against a panel of Gram-positive and Gram-negative bacteria, including representative members of the ESKAPE group of pathogens. Weak to moderate antibacterial activity was observed in disk diffusion assays against Gram-positive bacteria, including *Enterococcus faecium, Staphylococcus aureus, Bacillus cereus, Bacillus subtilis* and *Mycobacterium smegmatis* (**Figure S32**), while Gram-negative bacteria were resistant. To quantify this anti-Gram-positive activity, minimum inhibitory concentration (MIC) values for the purified chitinimines were determined (**Table S9**). Interestingly, *C. koreensis* DSM 17726 exhibited growth-promoting rather than inhibitory effects on several *Salmonella* species (**Figure 5A and S33**). This activity was abolished in the Δ*chtnA* mutant, indicating a link to chitinimine biosynthesis. However, this phenotype did not occur when purified chitinimines were tested in a spot-on-lawn assay or when they were added to *Salmonella* cultures in liquid medium, suggesting that these metabolites contribute to, but are not solely responsible for, the growth-promoting effect (**Figure S34**). Given their amphiphilic nature, we next examined whether the chitinimines possess surfactant properties. Both the drop-collapse and microplate assay showed clear indications of surface activity, as evidenced by droplet spreading and distortion patterns characteristic of biosurfactants (**Figure 5B**). The chitinimines showed no antifungal activity against a panel of *Candida albicans, C. auris* and *C. glabrata* strains (**Figure S35**). Moreover, no cytotoxicity was observed against HeLa and CaCo-2 cells in a lactate dehydrogenase (LDH) release assay (**Figure S36**).

**Figure 5.**
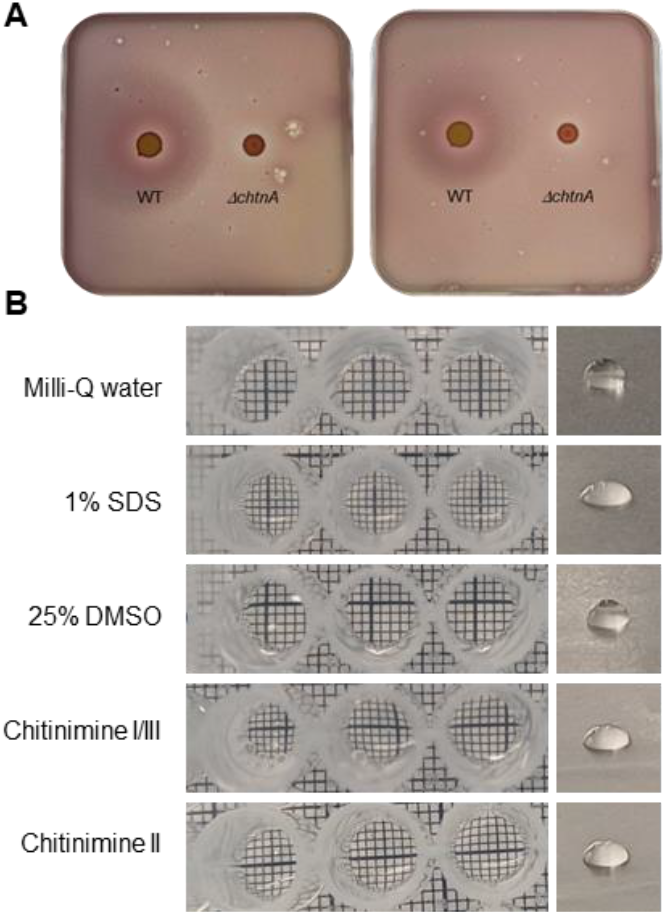
Growth-promoting properties of *C. koreensis* DSM 17726 and surfactant activity of the chitinimines. A) Growth-promoting effects of *C. koreensis* DSM 17726 and the Δ*chtnA* mutant were monitored with a colony overlay assay using *Salmonella enterica* 14029 (left) or *Salmonella enteritidis* ATCC 13046 (right). B) Surfactant activity of the chitinimines was observed in a drop-collapse (right) and microplate (left) assay.

## Conclusion

Bacteria have evolved a remarkable ability to mix and match different types of biosynthetic machinery to generate structurally and functionally diverse natural products. Hybrid assembly lines that combine FAS, PKS and NRPS enzymes are well known, and increasing reports of pathways that also recruit terpene or ribosomally synthesized and post-translationally modified peptide (RiPP) biosynthetic enzymes highlight the exceptional evolutionary plasticity of microbial secondary metabolism.^[41-43]^ Among these, the zeamine, fabclavine and megapolipeptin pathways represent the first examples of triple hybrid systems that merge PUFA synthase-like, PKS and NRPS biosynthetic machinery to assemble amphiphilic metabolites featuring specialized lipid moieties.

Here, we systematically charted the biosynthetic landscape of such hybrid peptide-polyketide-specialized lipid pathways across bacteria, revealing their diversity, distribution and evolutionary relationships. Using a targeted genome mining approach, we identified more than 60 previously unrecognized BGCs that combine PUFA synthase-like, PKS and NRPS modules in Gram-positive and Gram-negative bacteria from diverse taxonomic lineages. Comparative analysis showed that these triple hybrid systems have diversified significantly through domain reshuffling, gene fusion and recruitment of auxiliary enzymes to expand their catalytic abilities.

We experimentally validated our genome mining approach by isolating and elucidating the structures of the chitinimines, novel triple hybrid metabolites produced by a newly identified hybrid PKS-NRPS-PUFA synthase-like pathway in *Chitinimonas koreensis* DSM 17726. The chitinimines have very unusual structures, featuring a C22 polyunsaturated lipid chain linked to a macrocyclic peptide-polyketide and decorated with a pyruvate-derived cyclic acetal. Detailed bioinformatic analysis of their BGC enabled us to propose a plausible pathway for their biosynthesis, involving a PUFA synthase-like complex that exerts precise control over chain length and β-carbon processing during lipid chain assembly. Further studies will be needed to elucidate how this remarkable level of enzymatic programming and specificity is achieved.

Despite originating from the same phylogenetic clade as the megapolipeptin biosynthetic pathway, the chitinimines exhibit clear structural differences. They contain a highly unsaturated lipid chain integrated into a macrocyclic peptide framework, rather than a long-chain ω-oxo-fatty acid conjugated to a linear peptide as in the megapolipeptins, indicating functional divergence even between closely related pathways.^[24]^ Interestingly, the chitinimines also bear a striking structural resemblance to the bolagladins, bolaamphiphilic lipodepsipeptides from *Burkholderia gladioli*, which contain a citrate-derived fatty acid decorated with several polar functionalities and linked to a depsitetrapeptide.^[44-45]^ Despite their similarity in overall design, the bolagladins are assembled via a fundamentally different biosynthetic logic, relying on type III PKS-like enzymes that act in concert with primary fatty acid metabolism and an NRPS. These pathways elegantly illustrate Nature’s enzymatic ingenuity in generating related chemical scaffolds through distinct biosynthetic strategies.

The chitinimines were found to exhibit moderate antibacterial activity against Gram-positive bacteria, along with clear surfactant properties, consistent with their amphiphilic architecture. Intriguingly, they also contributed to a growth-promoting effect of the producing strain *C. koreensis* towards *Salmonella* species. While the underlying mechanisms remain unclear, these findings suggest that chitinimines may serve as multifunctional metabolites, mediating both antagonistic and cooperative interactions. Altogether, this work broadens the chemical and ecological scope of hybrid peptide-polyketide-specialized lipid metabolites and lays the foundation for future studies into their ecological roles.

## Data availability statement

The chitinimine BGC will be deposited in the MIBiG database under the accession number BGC0003185. The ChtnH putative pyruvyl transferase enzyme will be deposited at the Minimum Information about a Tailoring Enzyme (MITE) database under the accession number MITE0000215. The antiSMASH files corresponding to the genome mining section will be archived in Zenodo record (link to be announced). Data supporting this preprint is included in the Supporting information. Other data supporting the findings of this study are available from the corresponding author upon reasonable request.

