## Supplementary information for "Charting the biosynthetic landscape of hybrid polyketide-nonribosomal peptide-specialized lipids"

### Experimental Procedures

#### Bacterial strains, plasmids, media and growth conditions

*Chitinimonas koreensis* DSM 17726, first described by Kim *et al*. (2006), was obtained from the Deutsche Sammlung von Mikroorganismen und Zellkulturen (DSMZ, Leibniz-Institut, Germany, DSM no. 17726) and used for chitinimine production.[1] The strain was cultured at 28°C in Reasoner's 2A (R2A) medium for liquid cultivation and on R2A agar (R2A supplemented with 15 g/L agar) for solid cultivation. Standard cloning procedures were performed using *Escherichia coli* SY327.[2] The auxotrophic strain *E. coli* RHO3 was used for conjugative transfer of DNA into wildtype (WT) *C. koreensis*.[3] *E. coli* RHO3 was grown in Lysogeny Broth (LB) supplemented with 200 µg/mL 2,6-diaminopimelic acid (DAP) at 37°C. The suicide vector pSF100 was used for insertional mutagenesis in *C. koreensis*.[4] For plasmid selection, kanamycin at 50 µg/mL was used.

#### Genome mining for novel hybrid polyketide-nonribosomal peptide-specialized lipid biosynthetic pathways

Genome mining for novel hybrid PKS-NRPS-PUFA synthase-like clusters was carried out using antiSMASH (version 7.0), which was installed locally as a conda package.[5] A dataset comprising 62,706 NRPS-containing bacterial genomic regions in Genbank format, obtained from the antiSMASH database (version 4), was analyzed via a custom-designed search rule, which we termed 'zeamine-like'.[6] This rule integrates pre-existing profile Hidden Markov Models (pHMMs) targeting PKS, NRPS and PUFA synthase-like biosynthetic machinery (cds(Condensation and (AMP-binding or A-OX)) and cds(PKS_AT and (PKS_KS or ene_KS or mod_KS or hyb_KS or itr_KS or tra_KS)) and (hglE or hglD or PUFA_KS)). A 'relaxed' level of strictness was applied, allowing for the detection of incomplete clusters lacking one or more functional components. The maximum allowed distance between core genes was set at 20 kbp, and an additional 20 kbp was included beyond the core genes to define the protocluster boundaries.

All positive hits were manually curated and further analysed via antiSMASH with all optional parameters enabled. Particular attention was given to the KnownClusterBlast tool, which compares query clusters against the MIBiG database to identify similarities with experimentally characterized biosynthetic gene clusters (BGCs).[7] Initial hits were excluded if (i) the predicted PKS gene corresponded to a misannotated *pfaBC* homolog, (ii) the co-localization and synteny of PKS-, NRPS-, and PUFA synthase-like genes were not conserved among closely related strains, as determined using the *ClusterBlast* tool in antiSMASH, (iii) the *pfa*-like and PKS–NRPS genes were separated by numerous intervening genes lacking operon-like organization, and/or (iv) more than 71% of the biosynthetic genes matched a known MIBiG reference cluster, indicating the likely detection of a known BGC.

Sequences of either ketosynthase (KS) domains encoded in *pfaA* homologs (**Table S10**) or KS-CLF heterodimers encoded in *pfaC*-like genes (**Table S11**) were manually extracted and aligned using Clustal Omega via Geneious Prime (version 2024.0.5). Maximum likelihood phylogenetic trees were constructed using the IQ-TREE web server with default parameters and 10,000 bootstrap alignments.[8] The resulting trees were visualized with interactive Tree Of Life (iTOL) (version 7.1).[9]

To explore the diversity of zeamine and fabclavine BGCs from both known producers (e.g., *Dickeya* and *Serratia* spp.) and newly identified ones (e.g., *Chitinivorax tropicus*, *Paludibacterium paludis* and *Bowmanella denitrificans*), the Biosynthetic Gene Similarity Clustering And Prospecting Engine (BiG-SCAPE) (version 2.0.0-beta.5) tool was used.[10] The Affinity Propagation's internal preference parameter was set at a negative value and the cutoff was maximized (value of 0.99) to minimize the formation of multiple Gene Cluster Families (GCFs).

Comparative analyses of BGCs and/or genomic regions were visualized with the clinker tool from the CompArative GEne Cluster Analysis Toolbox (CAGECAT).[11]

#### Insertional mutagenesis of chitinimine biosynthetic gene *chtnA*

Insertional mutagenesis was performed using the *pir* replication-dependent pSF100 suicide plasmid. Primers amplifying a ~1000 bp region of the gene *chtnA* (Fw: 5'-AGGTCTCATATCGCCATCATCGGCGCCGCCTGCC-3'; Rev: 5'-TGGTCTCAGCTCCCGCTTCGAGGTGGCCGATATTGGTCTTGATCGAGCC-3') were designed with *Bsa*I restriction sites at the 5'-end to enable directional cloning of the PCR products into pSF100. Analogous primers were designed to amplify the pSF100 plasmid (Fw: 5'-TGGTCTCAGATATCGCATGCGGTACCTCTAGAAG-3'; Rev: 5'-AGGTCTCAGAGCTCTCCCGGGAATTCGATC-3'). The PCR products were digested with *Bsa*I and ligated using T4 DNA ligase. All enzymes and kits were purchased from Thermo Fisher Scientific and used according to the manufacturer's instructions unless otherwise stated. Next, chemically competent *E. coli* SY327 cells were transformed with the ligation mixtures and transformants were selected on LB agar plates supplemented with 50 µg/mL kanamycin. Plasmids were isolated from kanamycin-resistant colonies using the GeneJET plasmid miniprep kit and their sequence was confirmed by Sanger sequencing (Eurofins genomics) using vector primers (Fw: 5'-GCGATTCAGGCCTGGTATG-3'; Rev: 5'-CGCACTGAGAAGCCCTTAG-3').The validated construct was introduced into chemically competent *E. coli* RHO3 cells and transformants were selected on LB agar plates supplemented with 200 µg/mL DAP and 50 µg/mL kanamycin. For conjugative transfer of the construct into WT *C. koreensis,* a biparental mating protocol from Garcia (2018) was adapted.[12] Specifically, liquid cultures of *C. koreensis* and *E. coli* RHO3 were cultured overnight and subsequently diluted to a final optical density at 600 nm ($OD_{600}$) of 0.5 – 0.7. *E. coli* cells were subsequently harvested via centrifugation (10 min, 4600 rpm) and resuspended in 5 ml of liquid R2A medium. *C. koreensis* acceptor cells were spread a sterile swap on half of an R2A agar plate supplemented with 200 µg/mL DAP and 10 mM $MgCl_2$. *E. coli* RHO3 donor cells were spread on top of the acceptor cells. One fourth of the plate was streaked with *C. koreensis* control cells and the other fourth with *E. coli* RHO3 control cells. The bacteria were incubated at 28°C for 48 hours. Bacteria from each section were collected with a swap and restreaked onto selective R2A agar plates supplemented with 50 µg/mL kanamycin, 10 mM $MgCl_2$ and no DAP. Plates were incubated at 28°C for five days to observe *C. koreensis* cells in the conjugation section, with no growth in the control areas. The correct integration of pSF100 into the genomic DNA of *C. koreensis* was verified via junction PCR and Sanger sequencing. The primers used for junction PCR were the pair 5'-CCGAATTGCCGTATTCACCGTTC-3' (complementary to a region upstream of the *chtnA* gene) and 5'-CGCACTGAGAAGCCCTTAG-3' (complementary to a region downstream of the MCS site in pSF100); and the pair 5'-GATTGGGCCGGCTGAAGTTC-3' (complementary to a region downstream of the homology region from the *chtnA* gene) and 5'-GCGATTCAGGCCTGGTATG-3' (complementary to a region upstream of the MCS site in pSF100).

#### Comparative metabolic profiling and UHPL-ESI-Q-TOF-MS(/MS) analyses

To investigate the effect of the insertional mutagenesis on the metabolite profile of *C. koreensis*, WT and mutant *C. koreensis* cultures were streaked on Basal Salts Medium (BSM) ($K_2HPO_4.3H_2O$ 4.25 g/L, $NaH_2PO4.H_2O$ 1 g/L, $NH_4Cl$ 2 g/L, $MgSO_4.7H_2O$ 0.2 g/L, $FeSO_4.7H_2O$ 0.012 g/L, $MnSO_4.H_2O$ 0.003 g/L, $ZnSO_4.7H_2O$ 0.003 g/L, $CoSO_4.7H_2O$ 0.001 g/L, nitrilotriacetic acid 0.1 g/L, casamino acids 0.5 g/L, yeast extract 0.5 g/L) agar plates supplemented with 4 g/L glucose.[13] Following incubation for four days at 28°C, the agar-grown cultures were extracted with ethyl acetate for 1 hour under minimal light exposure. The resulting extracts were dried by rotary evaporation *in vacuo*. The dried extracts were then resuspended in methanol, centrifuged for 1 min at 13,200 rpm and analyzed via UHPLC-ESI-Q-TOF-MS.

UHPLC-ESI-Q-TOF-MS analyses were performed using a Dionex UltiMate 3000 UHPLC coupled to a Zorbax RRHP Eclipse Plus C18 column (2.1x100 mm, 1.8 µL) connected to a Bruker Impact II mass spectrometer. Mobile phases consisted of water (A) and acetonitrile (B), each supplemented with 0.1% formic acid. The following gradient was used at a flow rate of 0.200 mL/min: 0–2.5 min 5% B, 2.5–14 min 5-100% B, 14-19 min 100% B, 19-20.4 min 100-5% B, 20.4-25 min 5% B. The mass spectrometer was operated in positive ion mode with a scan range of 50-3000 m/z. Source conditions were: end plate offset at −500 V; capillary at −4500 V; nebulizer gas ($N_2$) at 1.6 bar; dry gas ($N_2$) at 8 L min−1; dry temperature at 180 °C. Ion transfer conditions were: ion funnel RF at 200 Vpp; multiple RF at 200 Vpp; quadrupole low mass at 55 m/z; collision energy at 5.0 eV; collision RF at 600 Vpp; ion cooler RF at 50–350 Vpp; transfer time at 121 µs; pre-pulse storage time at 1 µs. Calibration was performed with 1 mM sodium formate through a loop injection of 20 µL at the start of each run. Mass spectra were analysed using the Compass Data Analysis software (Bruker).

High-resolution LC-ESI-MS/MS analyses were performed using a Dionex UltiMate 3000 UHPLC coupled to a Zorbax RRHP Eclipse Plus C18 column (2.1x100 mm, 1.8 µL) connected to a Bruker Impact II mass spectrometer. Mobile phases consisted of water (A) and acetonitrile (B), each supplemented with 0.1% formic acid. The following gradient was used at a flow rate of 0.200 mL/min: 0–2.5 min 5% B, 2.5–14 min 5-100% B, 14-19 min 100% B, 19-20.4 min 100-5% B, 20.4-25 min 5% B. The mass spectrometer was operated in positive ion mode using the following parameters: scan range: 50-1500 m/z, nanospray voltage: 3.5 kV, source temperature: 200°C, normalized collision energy: 10 eV-20-30-40, isolation window: ± 8 Da. The lock mass 150.0000 was used as an internal calibrant. The source conditions were: end plate offset at −500 V; capillary at −3500 V; nebulizer gas ($N_2$) at 40 psi; dry gas ($N_2$) at 8 L min−1; dry temperature at 200 °C. Ion transfer conditions were: ion funnel RF at 350 Vpp; multiple RF at 350 Vpp; quadrupole low mass at 150 m/z; collision energy at 5.0 eV; collision RF at 1500 Vpp; ion cooler RF at 50–350 Vpp; transfer time at 80 µs; pre-pulse storage time at 10 µs. Calibration was performed with 1 mM sodium formate through a loop injection of 20 µL at the start of each run. Mass spectra were analysed using the Compass HyStar software (Bruker).

#### Isolation and structure elucidation of the chitinimines

For chitinimine production, *C. koreensis* DSM 17726 was grown on BSM agar plates supplemented with 4 g/L glucose. Following incubation for 4 days at 28°C, the cells and the agar were extracted with ethyl acetate under minimal light exposure. Extracts were dried by rotary evaporation *in vacuo*, and the resulting solids were resuspended in 50%

acetonitrile in water. The chitinimine-containing extract was then fractionated by preparative HPLC on a Shimadzu Nexera Prep instrument equipped with a Shimadzu Shim pack GIS column (5 µm, C18, 100 Å, 250 × 10 mm), monitoring absorbance at 190 nm. Mobile phases consisted of water (A) and acetonitrile (B), each supplemented with 0.1% formic acid. The following gradient was used at a flow rate of 5 mL/min: 0–2.5 min 5% B, 2.5–30 min 5-100% B, 30-35 min 100% B, 35-36 min 100-5% B, 36-38 min 5% B. Chitinimine-containing fractions were identified via UHPLC-ESI-Q-TOF analysis. These fractions were concentrated and lyophilized prior to further characterization. The structure of the chitinimines was elucidated using a combination of UHPLC-ESI-Q-TOF-MS and 1- and 2-D NMR experiments.

For NMR spectroscopic analyses, each sample was separately dissolved in 550 µL of DMSO. All spectra were measured on a Bruker Neo spectrometer operating at 600Mhz with quadruple cryoprobe ($^{1}H$,$^{31}P$,$^{15}N$,$^{13}C$) and processed with Topspin software. The 2D DQF-COSY[14],TOCSY[15], and NOESY[16] spectra were recorded with a sweep width of 6600 Hz in both dimensions. The total TOCSY mixing time was set to 62 ms. Zero-quantum interference in TOCSY spectra was eliminated by gradients.[14] Homonuclear spectra were acquired with 8 to 72 scans depending on the sample concentration, 4096 data points in ($t_2$) and 512 to 1024 FIDs in ($t_1$). The data were apodized with a shifted sine-bell square function in both dimensions and processed to a 4K × 1K matrix. The NOESY experiments were acquired with mixing time 200 ms. Natural abundance [$^{1}H$, $^{13}C$]-HSQC[17] were recorded with sensitivity enhancement and gradient coherence selection optimized for multiplicity editing with negative signals for $CH_2$ moieties and positive signals for CH and $CH_3$ groups ($^{1}J_{CH}$ = 145 Hz) using 8 to 32 scans (depending on sample concentration) and 1K/4K complex data points and 150/11 ppm spectral widths in $t_1$ and $t_2$, respectively. For the most concentrated sample (chitinimine I/III) a 2D HSQC-TOCSY was also obtained using an HSQC building block followed by a clean MLEV17 TOCSY transfer step with 80 ms mixing time.[18] The spectrum was recorded using 40 scans and 512/4K complex data points and 150/11 ppm spectral widths in $t_1$ and $t_2$, respectively. Natural abundance [$^{1}H$, $^{13}C$]-HMBC were measured with 64 scans and 4096/512 complex data points and 11/230 ppm spectral widths in $t_2$ and $t_1$, respectively ($^{1}J_{CH}$ = 145 Hz and $^{3}J_{CH}$ = 4 Hz).

#### Determination of absolute stereochemical configurations using Marfey's method

The absolute stereochemistry of the amino acid constituents of the chitinimines was determined using Marfey's method. The chitinimines were subjected to hydrolysis with 1 M HCl for 24 hours at 100°C, and subsequently derivatized with 1-fluoro-2-4-dinitrophenyl-5-L-alanine amide (FDAA) as described by Tanino *et al.* (2010). The resulting diastereomers were compared to Marfey's derivatives of the appropriate D- and L-amino acid standards by UHPLC-ESI-Q-TOF-MS.[19] All amino acid standards were commercially available, except for (3S,4S)-4-amino-3-hydroxypentanoic acid, which was acquired with an FMOC protecting group. For deprotection, 50 mg of Fmoc-(3S,4S)-4-amino-3-hydroxy-pentanoic acid was dissolved in a water-DMF mixture to a final concentration of 0.1 M, followed by reaction with excess piperidine (5.0 equivalents) in the presence of DCM. This reaction was allowed to proceed overnight at room temperature. The mixture was then dried *in vacuo*, washed three times with water and dissolved in methanol for UHPLC-ESI-Q-TOF-MS analysis to confirm successful deprotection.

The authentic standards were prepared by mixing 1.0 equivalent of each amino acid, dissolved in 1 M $NaHCO_3$, with 2.0 equivalents of FDAA dissolved in acetone. The mixtures were incubated for 2 hours at 40°C, followed by quenching with 1 M HCl. The solvents were removed under reduced pressure, and the resulting residues were dissolved in DMSO and diluted 10 times with methanol. The UHPLC-ESI-Q-TOF-MS analyses were performed as described above using the following gradient at a flow rate of 0.200 mL/min: 0–2.5 min 5% B, 2.5–7 min 5-40% B, 7-17 min 40% B, 17-19 min 40-100% B, 19-24 min 100% B, 24-25.4 min 100-5% B, 25.4–30 min 5% B.

#### Acid hydrolysis of the chitinimines

For acid hydrolysis, 4 mg of sample containing a mixture of chitinimine I/III and chitinimine II at a ratio of 8:1 was dissolved in 400 µL of DCM and 100 µL of water. 100 µL of TFA was added dropwise, and the resulting mixture stirred at room temperature for 24 hours. Solvents were removed under reduced pressure and the resulting product was washed twice with 100 µL of DCM. The hydrolysed chitinimines were dissolved in 500 µL of methanol and analyzed with UHPLC-ESI-Q-TOF-MS.

#### Phylogenetic and bioinformatic analysis of the chitinimine biosynthetic pathway

The hybrid PKS-NRPS-PUFA synthase-like biosynthetic gene cluster from *C. koreensis* was analyzed in-depth using antiSMASH version 7.0, with all optional parameters for extended analysis enabled. The domain organization of the FAS, PKS and NRPS subunits was determined through InterPro protein classification and NCBI's Conserved Domain Database (CDD) searches. Putative functions were assigned to the proteins encoded by each gene using a combination of NCBI BlastP and UniProt BLAST. Proteins belonging to the same enzyme class were subjected to Clustal Omega (EMBL-EBI) and the resulting multiple sequence alignment were examined for conserved residues indicative of catalytic activity and stereochemical or substrate specificity. The amino acid substrate selectivity of the adenylation domains was predicted using antiSMASH and PARAS.[20] Condensation domain sequences were extracted and analyzed via the Natural Product Domain Seeker (version 2) (NaPDoS2).[21] Specifically, these sequences were aligned against 172 reference condensation domains from the NaPDoS2 database (**Table S8**), and the resulting neighbor-joining phylogenetic tree was visualized with iTOL.

### Bioactivity assays

The antimicrobial activity of the chitinimines was tested against the bacterial and fungal strains listed in **Table S12**.

Soft agar halo assays were carried out to compare the antibacterial activity of WT and mutant *C. koreensis* strains. 10 µL of stationary-phase cultures of *C. koreensis* WT and mutant strains were spotted onto BSM agar plates supplemented with glucose (4 g/L). Following incubation for four days at 28°C, stationary-phase cultures of the target strains were diluted 1:1000 in 15 ml of soft agar prepared in their preferred growth medium and poured over the surface of the BSM plates. After incubation for 24 hours at the optimal growth temperature of the target strains (**Table S12**), the presence and size of the inhibition zones was evaluated. If growth-promoting effects were observed instead of inhibition, the assay was repeated with the addition of iodonitrotetrazolium chloride (0.2 mg/mL) to the soft agar for improved visualization of bacterial growth.

The bioactivity of the purified chitinimines was also determined via plate lawn assays. A mixed sample containing both chitinimine I/III and II at a ratio of 10:1, respectively, and at a final concentration of 1 mg/mL (dissolved in DMSO) was tested. Agar plates were overlaid with lawns of stationary-phase cultures of target bacterial strains, diluted to an $OD_{600}$ of 0.03. 10 µL of the chitinimine mixture was spotted onto sterile filter paper placed on top of the lawns. DMSO was used as a negative control. Plates were incubated for 24 hours under the preferred growth conditions for each target strain. Presence and size of inhibition halos were evaluated.

To assess the growth-promoting effect of purified chitinimines, *Salmonella enterica* 14029, *S. enteritidis* ATCC 13046 and *S. newport* C487 were cultured in the presence of the compounds and growth was monitored at $OD_{600}$. In a sterile 96-well microtiter plate, 2.5 µL of chitinimine I/III or chitinimine II (dissolved in DMSO) was added at a final concentration of 125 µg/mL to 97.5 µL of bacterial suspension ($10^6$ CFU/ml in LB broth). In control experiments, DMSO was added instead of the chitinimines, and non-inoculated LB medium was used to subtract background signal. The $OD_{600}$ was monitored every 15 minutes for 13 hours at 37°C in a CLARIOstar Plus (BMG LABTECH), and in between reads, cells were shaken at 300 rpm. The resulting growth curves were visualized with QurvE.[22]

Minimal inhibitory concentration (MIC) values for the chitinimines were determined by the broth microdilution method, following the guidelines from the Clinical Laboratory Standards Institute (CLSI) (document M07, 12th edition).[23] In a 96-well microtiter plate, 50 µL of serial twofold dilutions of the metabolites in Mueller Hinton (M-H) broth were mixed with 50 µL of bacterial suspension ($10^6$ CFU/mL in M-H broth). After incubation 18 hours at the preferred growth temperature of the target bacteria, MIC values were defined as the lowest concentrations that visibly inhibited bacterial growth.

For antifungal assays, pre-warmed RPMI-MOPS medium, supplemented with 0.2% glucose, was inoculated with fungal pathogen cultures (see **Table S12**) that had been washed twice with Phosphate Buffered Saline (PBS). The final inoculum was adjusted to an optical density at 600 nm ($OD_{600}$) of 0.001. In a sterile 96-well microtiter plate, wells were filled with 196 µL of the inoculated medium and 4 µL of either chitinimine I/III or chitinimine II at a concentration of 5 mg/mL (dissolved in DMSO). Plates were incubated for 24 hours at 37 °C for *Candidozyma auris* (formerly *Candida auris*) and *Nakaseomyces glabratus* (formerly *Candida glabrata*) strains, or at 30 °C for *Candida albicans* strains. Following incubation, the $OD_{600}$ was measured using a Synergy H1 plate reader (BioTek). To establish the background signal, a blank consisting of non-inoculated RPMI-MOPS was included. Wells supplemented with DMSO instead of compound served as growth controls for each pathogen. The percentage of growth inhibition relative to the control was calculated using the following formula:

$$\text{Inhibition (\%)} = \left(1 - \frac{ODsample - ODblank}{ODcontrol - ODblank}\right) * 100$$

The cytotoxicity of the chitinimines on mammalian cells was assessed using the CyQUANT™ LDH Cytotoxicity Assay Kit (Invitrogen, Thermo Fisher Scientific; Cat. No. C20300) according to the manufacturer's instructions. HeLa (ATCC CCL-2) and CaCo-2 (ATCC HTB-37) cells were used for this assay. Cultures were maintained and incubated during cytotoxicity determination at 37 °C in a humidified 5% $CO_2$ atmosphere. One day prior to compound exposure, cells were seeded in Nunclon Delta Surface 96-well plates (Thermo Fisher Scientific) at a density of 1 × $10^5$ cells/mL, with 100 µL of cell suspension per well. Cytotoxicity measurements followed the CyQUANT™ LDH protocol. 2 µL of chitinimine I/III or chitinimine II at an initial concentration of 5 mg/mL (dissolved in DMSO) and 8 µL of MilliQ-water were added to the wells. The compounds were substituted with DMSO for cell growth control. Wells containing untreated cells served as background controls, while maximum LDH release controls were obtained by lysing cells with the supplied lysis buffer. Exposure of the cells to the compound lasted 45 min until addition of reaction substrate. Absorbance of the resulting formazan product was measured at 490 nm and 680 nm spectrophotometrically, and percent cytotoxicity was calculated following the manufacturer's formula:

$$\text{Cytotoxicity (\%)} = \frac{\text{(Experimental LDH release - Spontaneous LDH release)}}{\text{(Maximum LDH release - Spontaneous LDH release)}} *100$$

For surfactant activity testing, a microtiter assay adapted from the method by Vaux[24], and described by Walter *et al.*[25], was employed. In a 96-well microtiter plate, 25 µL of either chitinimine I/III or chitinimine II (dissolved in DMSO) were added to 75 µL of Milli-Q water to obtain a final concentration of 1.25 mg/mL. 100 µL of either Milli-Q water, 1% sodium dodecyl sulphate (SDS) solution or 25% DMSO were used as negative, positive and solvent effect controls, respectively. The plate was placed on top of millimeter graph paper sheet, and distortion of the grid was interpreted as an indication of surfactant activity. In addition, the drop-collapse assay from Dose *et al.*[26], and originally described by Jain *et al.*[27], was applied for further confirmation. 10 µL droplets of 25% chitinimine I/III or chitinimine II (dissolved in DMSO) at a final concentration of 1.25 mg/mL and 75% Milli-Q water were spotted on top of Parafilm 'M'. 10 µL of either Milli-Q water, 1% SDS solution or 25% DMSO were spotted as negative, positive and solvent effect controls, respectively. Collapse of the droplets was interpreted as a positive indication of surfactant activity.

### Results and Discussion

#### Supplementary Figures

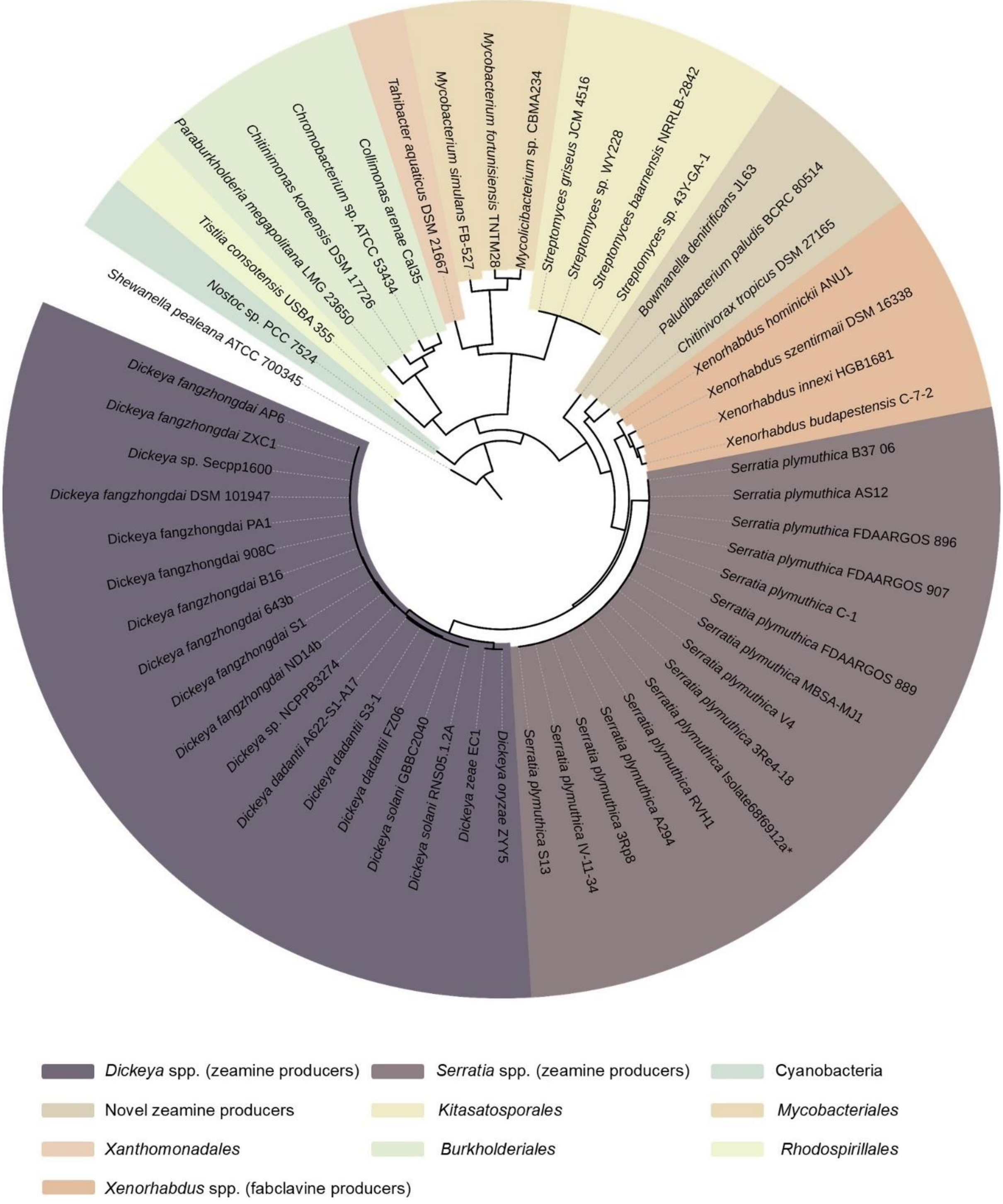

**Figure S1. Phylogenetic distribution of KS-CLF heterodimers from hybrid PKS-NRPS-PUFA synthase-like clusters.** A neighbor-joining tree was constructed from the KS-CLF domain sequences, with the PfaC KS-CLF didomain from the *S. pealeana* PUFA synthase used as the outgroup. Colored clades correspond to related pathways and producing organisms.

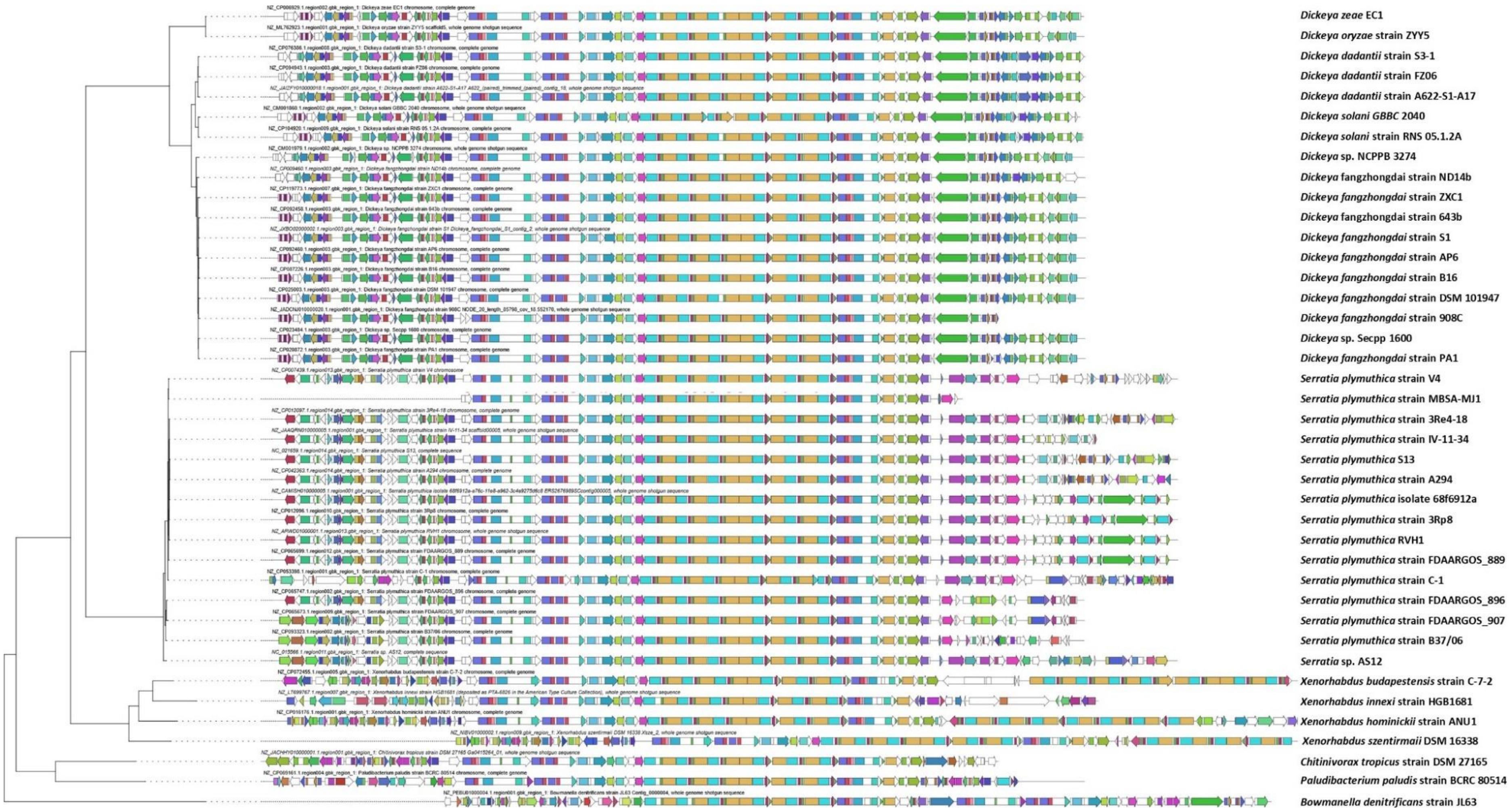

**Figure S2. Phylogenetic distribution and comparative analysis of the zeamine and fabclavine BGCs detected using the customized antiSMASH search.** The evolutionary relationships, synteny, and architectural diversity of the BGCs across multiple *Dickeya*, *Serratia* and *Xenorhabdus* species are illustrated. The newly identified zeamine-producing strains *Chitinivorax tropicus* strain DSM 27165, *Paludibacterium paludis* strain BCRC 80514 and *Bowmanella denitrificans* strain JL63 are also included in the analysis. Figure adapted from BiG-SCAPE.[10]

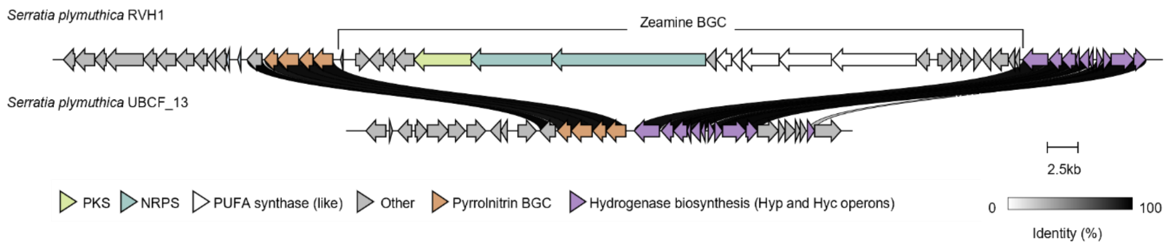

**Figure S3. Genomic comparison of the region surrounding the zeamine BGC in two closely-related *Serratia plymuthica* strains, revealing the genomic integration site of the cluster.** In strain RVH1, the zeamine BGC is located between the pyrrolnitrin BGC (colored in orange) and the hyp and hyc operons involved in hydrogenase biosynthesis (colored in purple). In contrast, strain UBCF_13 contains only these flanking regions, with the zeamine cluster absent, suggesting that this locus serves as the insertion site for the zeamine BGC. Figure adapted from clinker [11].

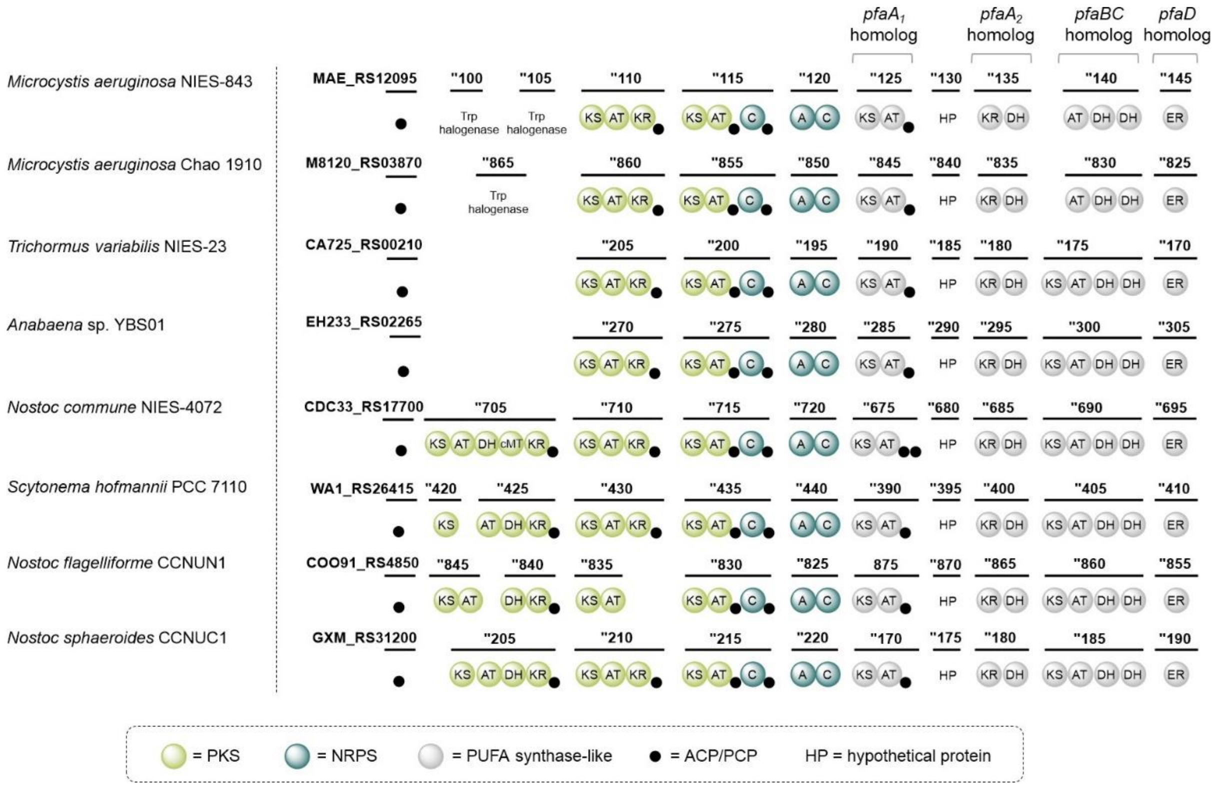

**Figure S4. Comparison of the domain architecture of the hybrid PKS-NRPS-PUFA synthase-like pathways in *M. aeruginosa, T. variabilis, Anabaena* sp., *N commune, S. hofmannii, N. flagelliforme* and *N. sphaeroides.*** PKS domains are colored in light green, NRPS domain in dark green, PUFA synthase-like biosynthetic machinery in grey, acyl and peptidyl carrier proteins in black. Putative tryptophan halogenases are indicated and conserved hypothetical proteins with a predicted Rossmann-fold NAD(P)-binding domain are labelled as HP.

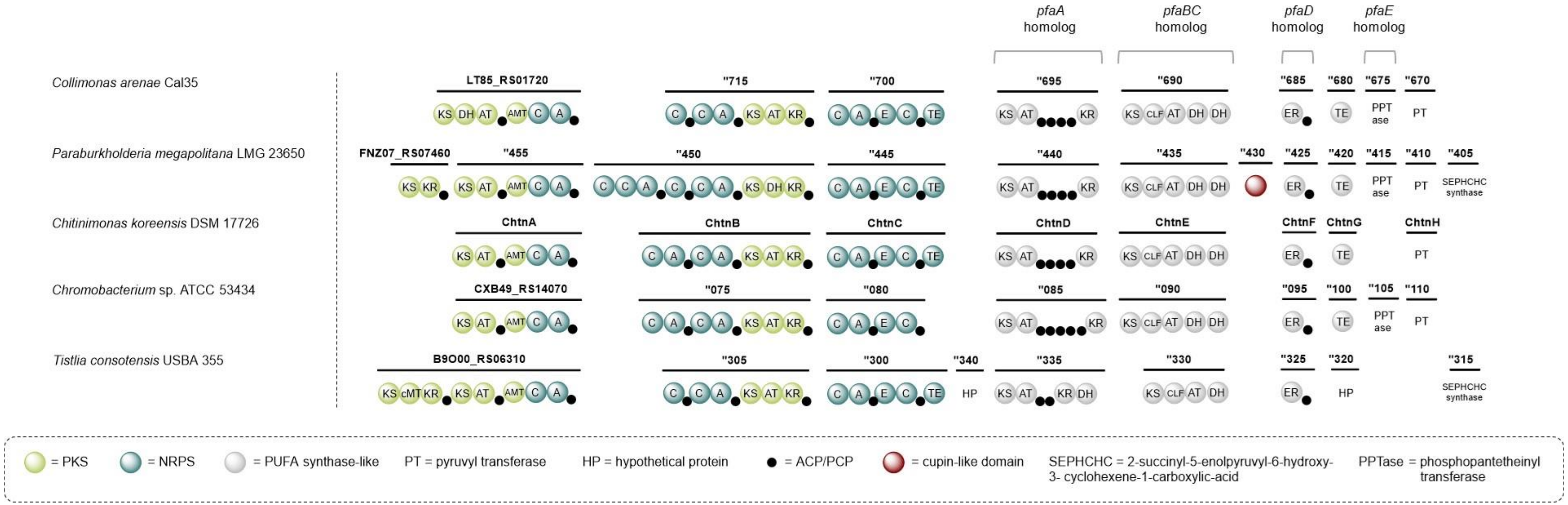

**Figure S5. Comparison of the domain architecture of the hybrid PKS-NRPS-PUFA synthase-like pathways in *C. arenae, P. megapolitana, C. koreensis, Chromobacterium* sp. and *T. consotensis.*** PKS domains are colored in light green, NRPS domains in dark green, PUFA synthase-like biosynthetic machinery in grey, acyl and peptidyl carrier proteins in black and cupin-like domains in brown. Putative tailoring enzymes and other proteins that are conserved across (a subset of) these pathways are shown with abbreviated labels.

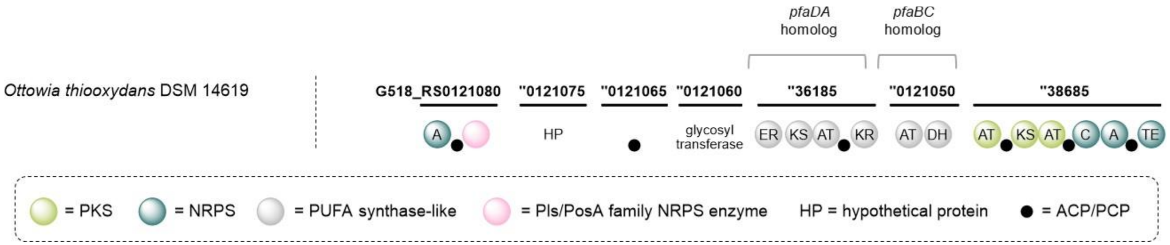

**Figure S6. Domain architecture of the hybrid PKS-NRPS-PUFA synthase-like pathway in *O. thiooxydans.*** PKS domains are colored in light green, NRPS domains in dark green, PUFA synthase-like biosynthetic machinery in grey, acyl and peptidyl carrier proteins in black, putative Pls/PosA family NRPS enzymes in pink, glycosyltransferases in orange and hypothetical proteins in blue-grey.

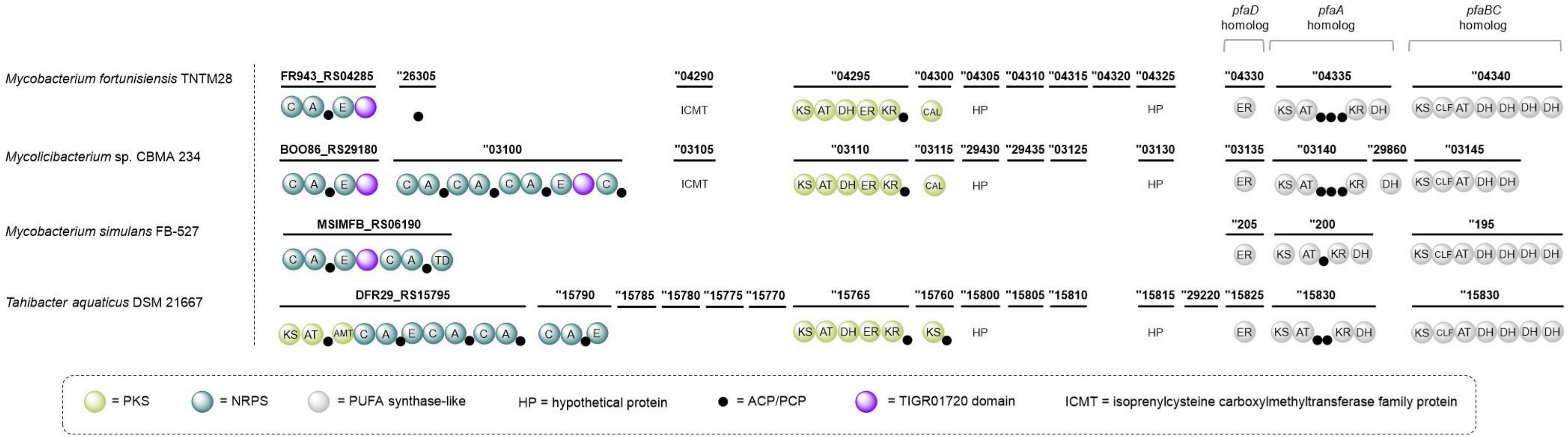

**Figure S7. Comparison of the domain architecture of the hybrid PKS-NRPS-PUFA synthase-like pathways in *M. fortunisiensis*, *Mycolicibacterium* sp., *M. simulans* and *T. aquaticus*.** PKS domains are colored in light green, NRPS domains in dark green, PUFA synthase-like biosynthetic machinery in grey, acyl and peptidyl carrier proteins in black and TIGR01720 domains in bright purple. Putative tailoring enzymes and other proteins that are conserved across (a subset of) these pathways are shown with abbreviated labels.

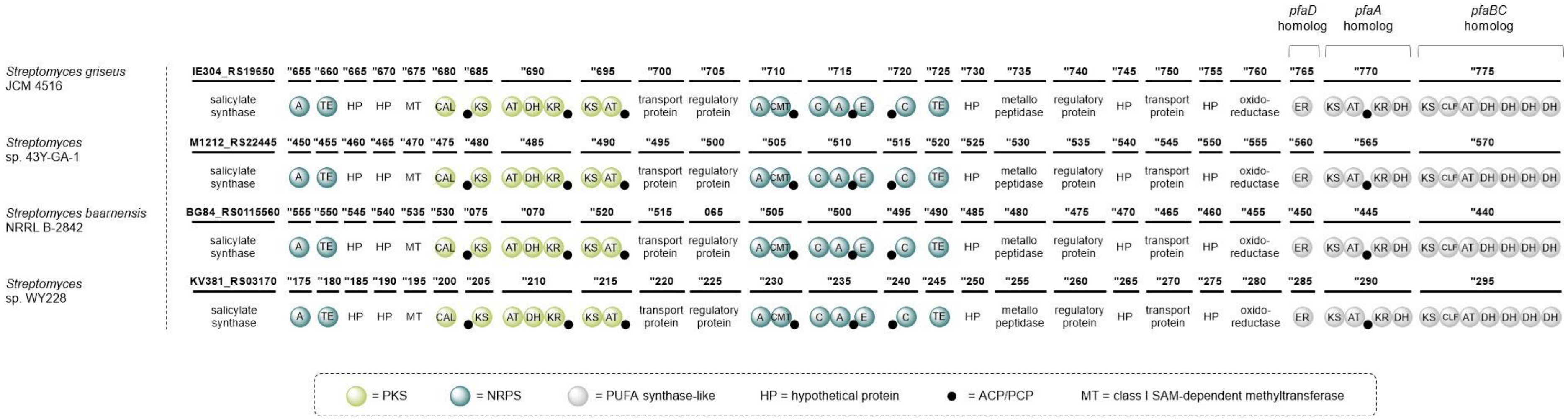

**Figure S8. Comparison of the domain architecture of the hybrid PKS-NRPS-PUFA synthase-like pathway in *S. griseus*, *Streptomyces*. sp. 43Y-GA-1, *S. baarnensis* and *Streptomyces*. sp. WY228.** PKS domains are colored in light green, NRPS domains in dark green, PUFA synthase-like biosynthetic machinery in grey, and acyl and peptidyl carrier proteins in black. Putative tailoring enzymes and other proteins that are conserved across (a subset of) these pathways are shown with abbreviated labels.

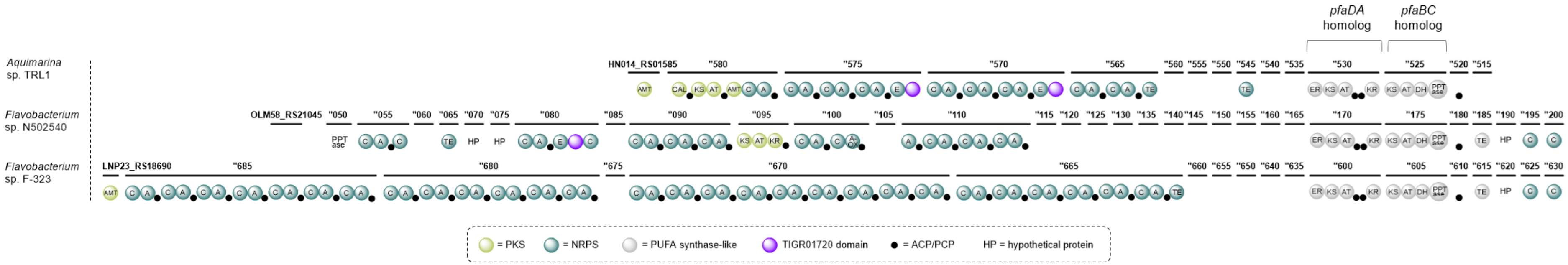

**Figure S9. Comparison of the domain architecture of the hybrid PKS-NRPS-PUFA synthase-like pathways in *Aquimarina* sp., *Flavobacterium* sp. N502540 and *Flavobacterium* sp. F-323.** PKS domains are colored in light green, NRPS domains in dark green, PUFA synthase-like biosynthetic machinery in grey, acyl and peptidyl carrier proteins in black and TIGR01720 domains in bright purple. Putative tailoring enzymes and other proteins that are conserved across (a subset of) these pathways are shown with abbreviated labels.

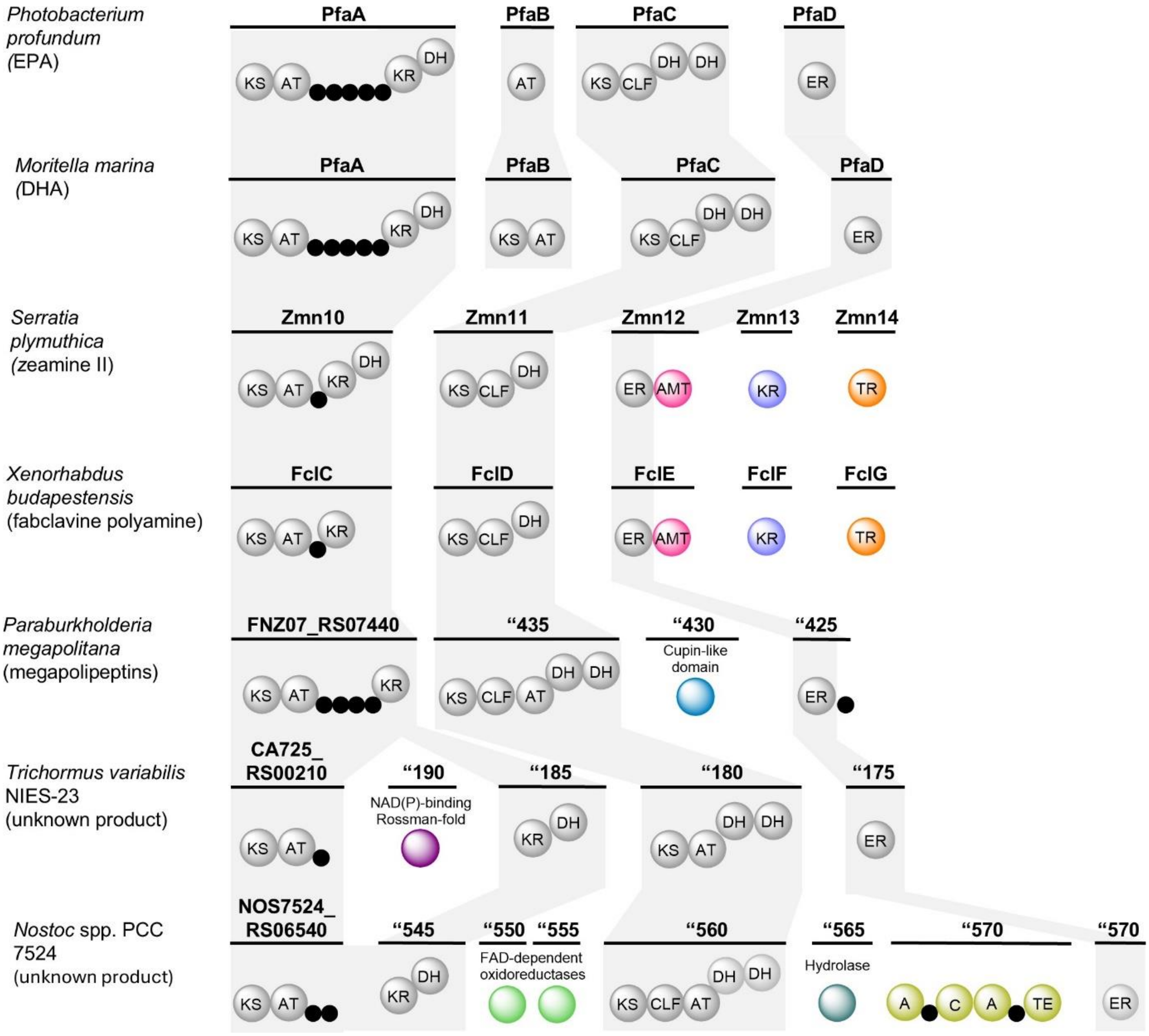

**Figure S10. Unusual catalytic domains in secondary lipid synthases.** Overview of unconventional catalytic domains that have been recruited by PUFA synthases (and homologs) over the course of evolution to diversify the structures of PUFAs.

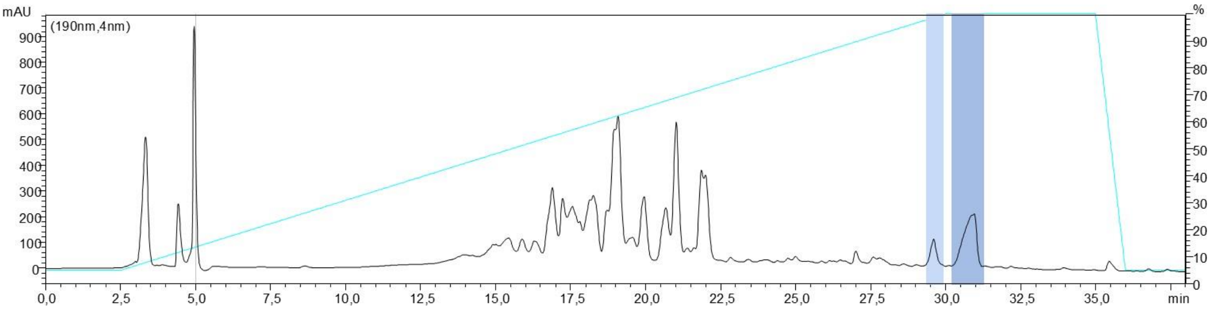

**Figure S11. Large-scale chitinimine production.** UV chromatogram from HPLC analysis (monitoring absorbance at 190 nm) of ethyl acetate extracts of agar plates grown with WT *C. koreensis* DSM 17726. The peaks that correspond to chitnimine II and chitinimine I/III are indicated with a light and dark blue box, respectively. The applied solvent gradient is shown as a blue line. The percentage of acetonitrile in water is indicated on the right y-axis.

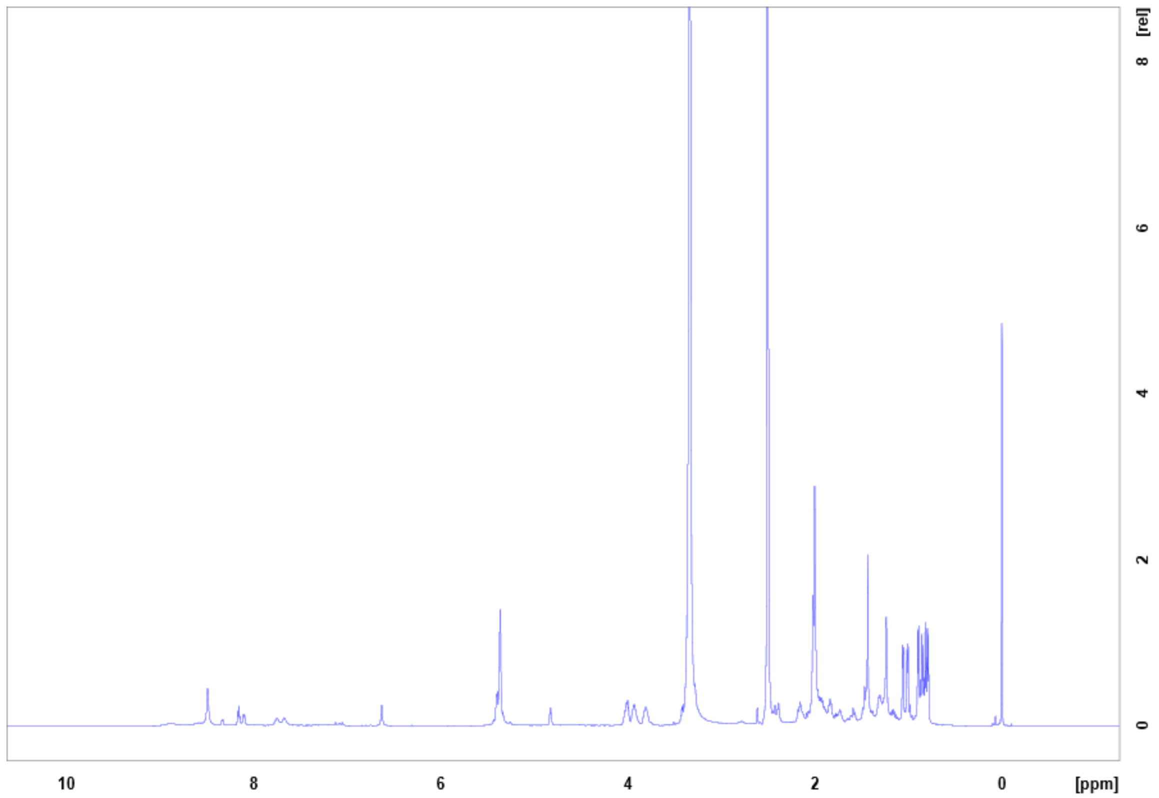

**Figure S12.** $^{1}H$ NMR spectrum of chitinimine I and III in DMSO-d6.

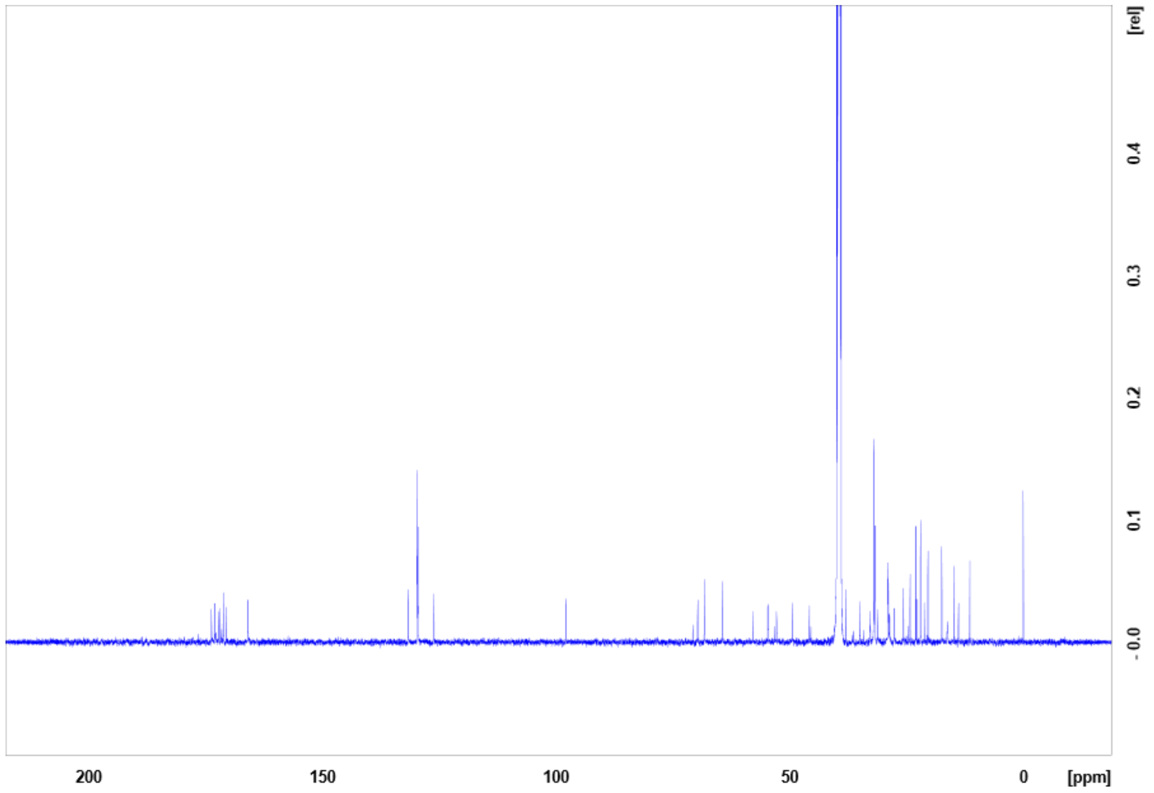

**Figure S13**. $^{13}C$ NMR spectrum of chitinimine I and III in DMSO-d6.

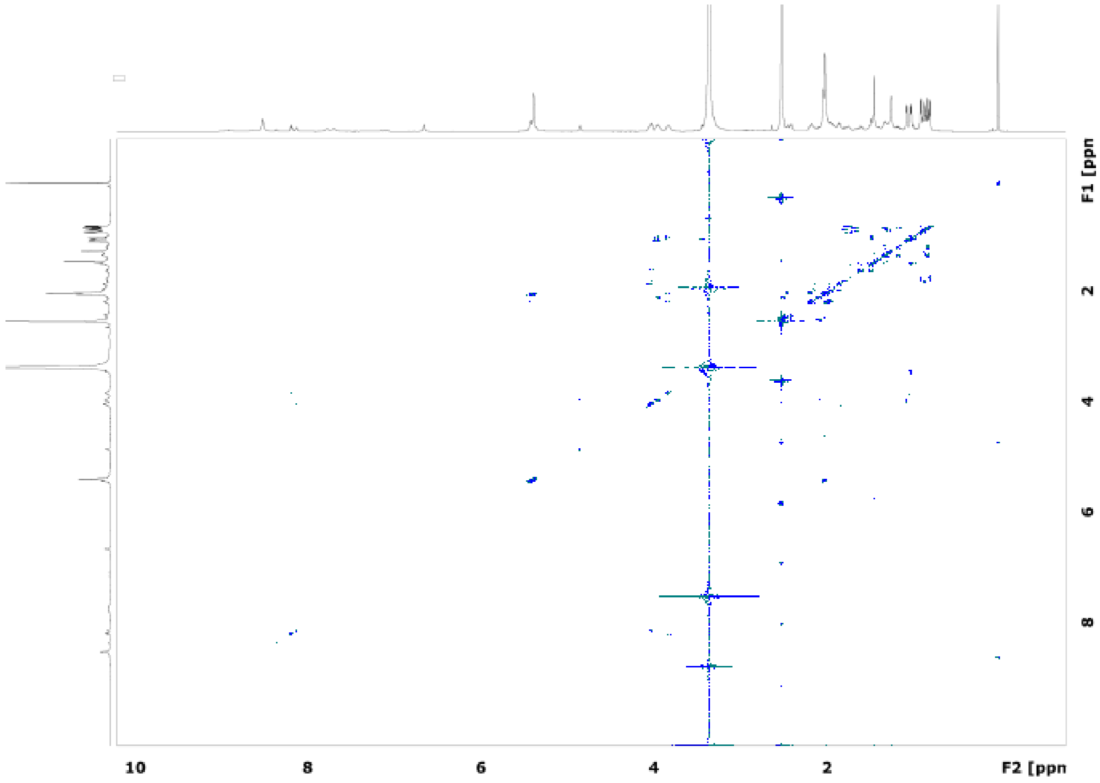

**Figure S14.** COSY spectrum of chitinimine I and III in DMSO-d6.

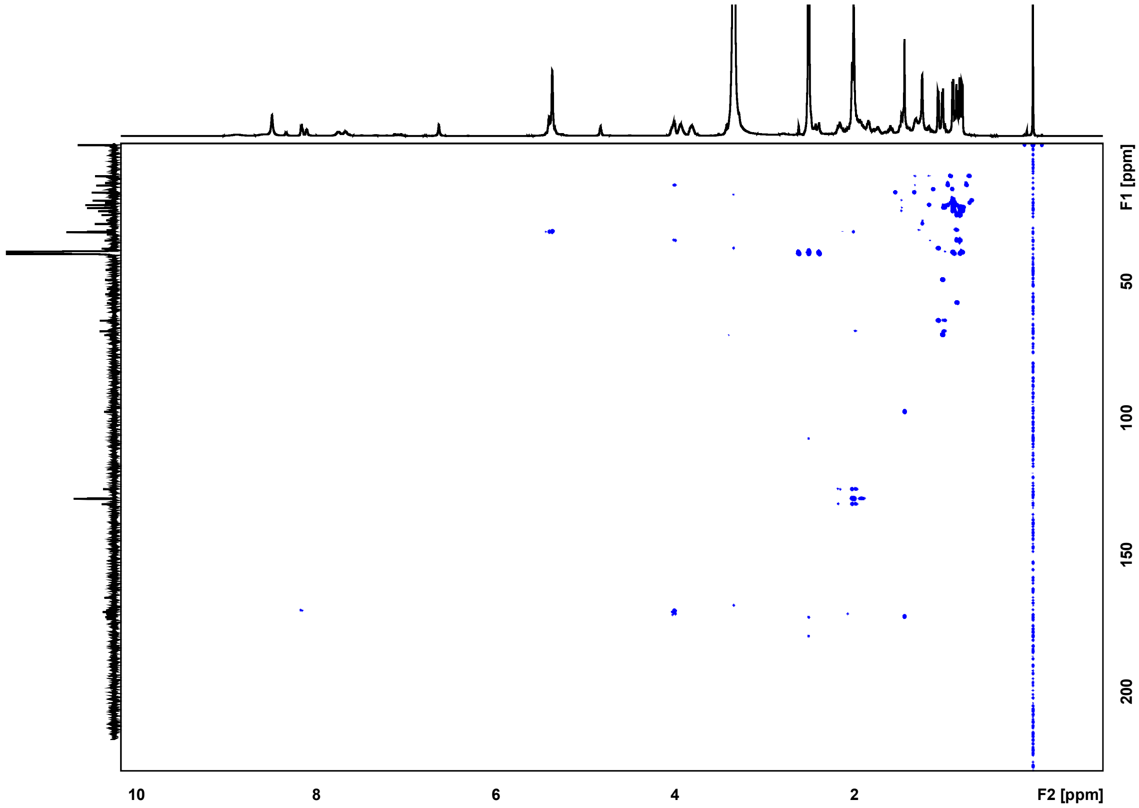

**Figure S15**. HMBC spectrum of chitinimine I and III in DMSO-d6.

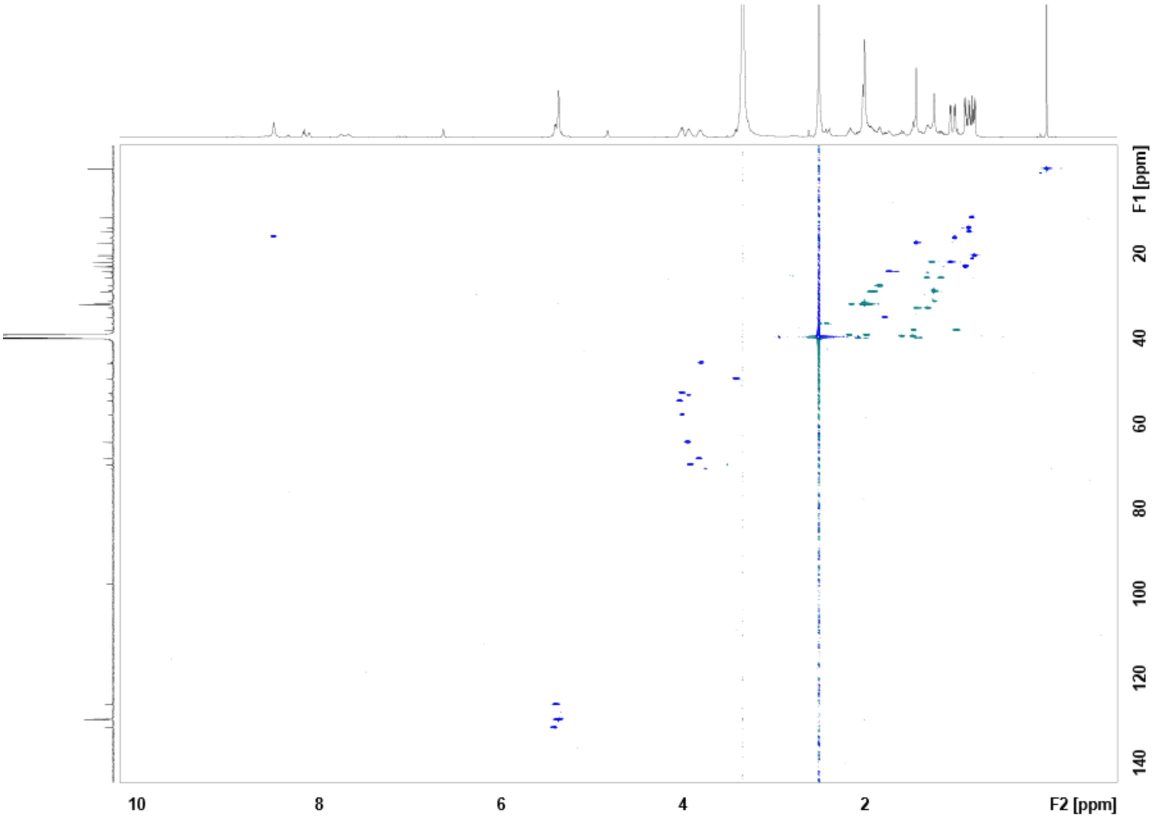

**Figure S16**. HSQC spectrum of chitinimine I and III in DMSO-d6.

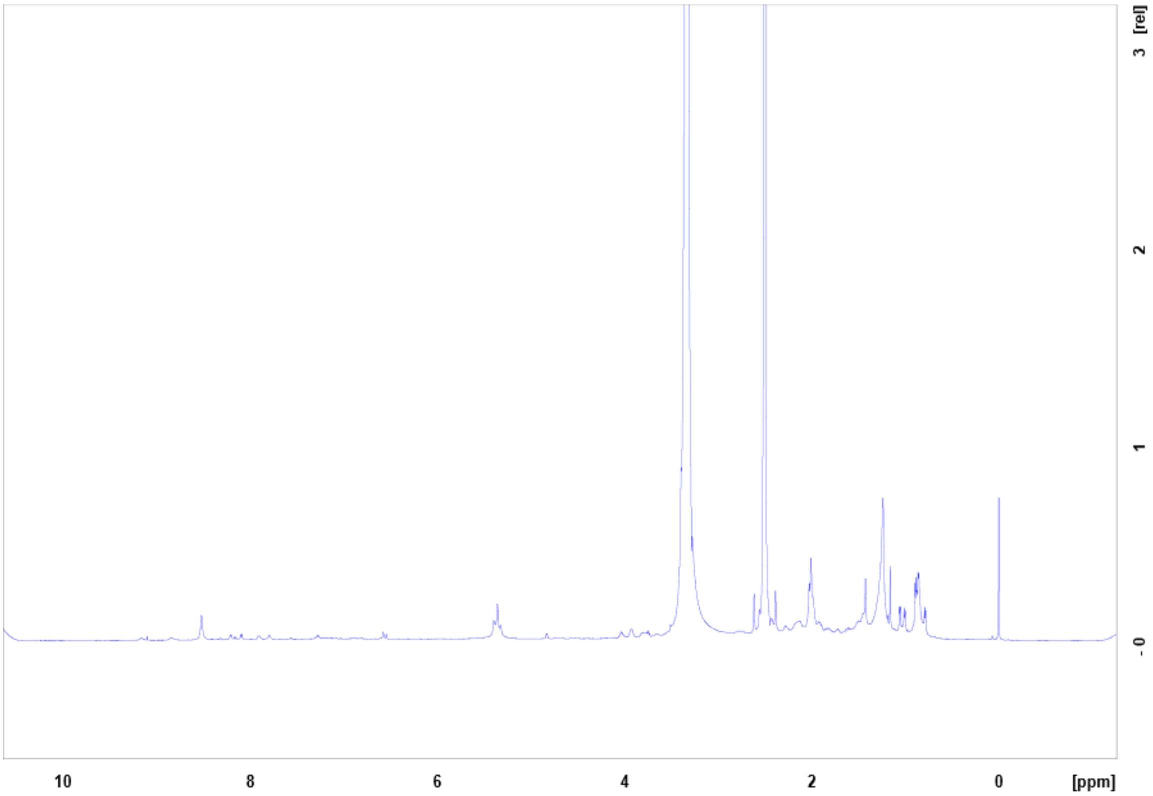

**Figure S17.** $^{1}$H NMR spectrum of chitinimine II in DMSO-d6.

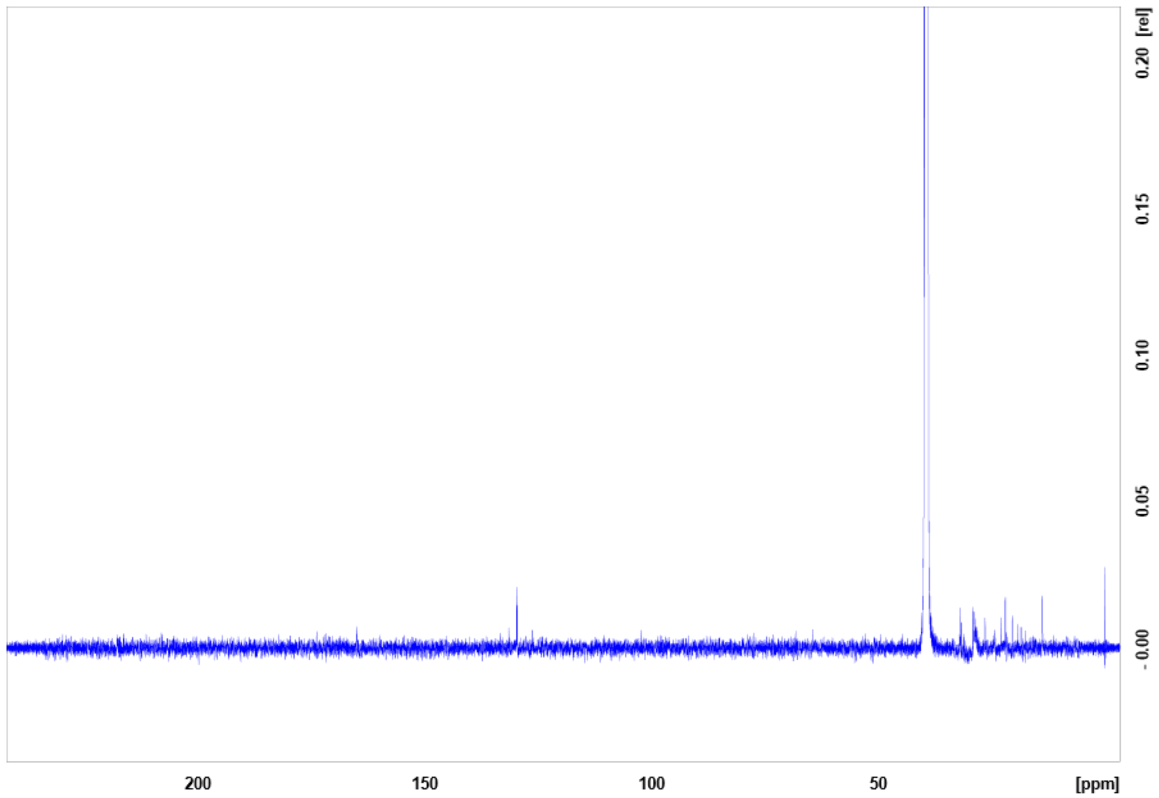

**Figure S18**. [13]C NMR spectrum of chitinimine II in DMSO-d6.

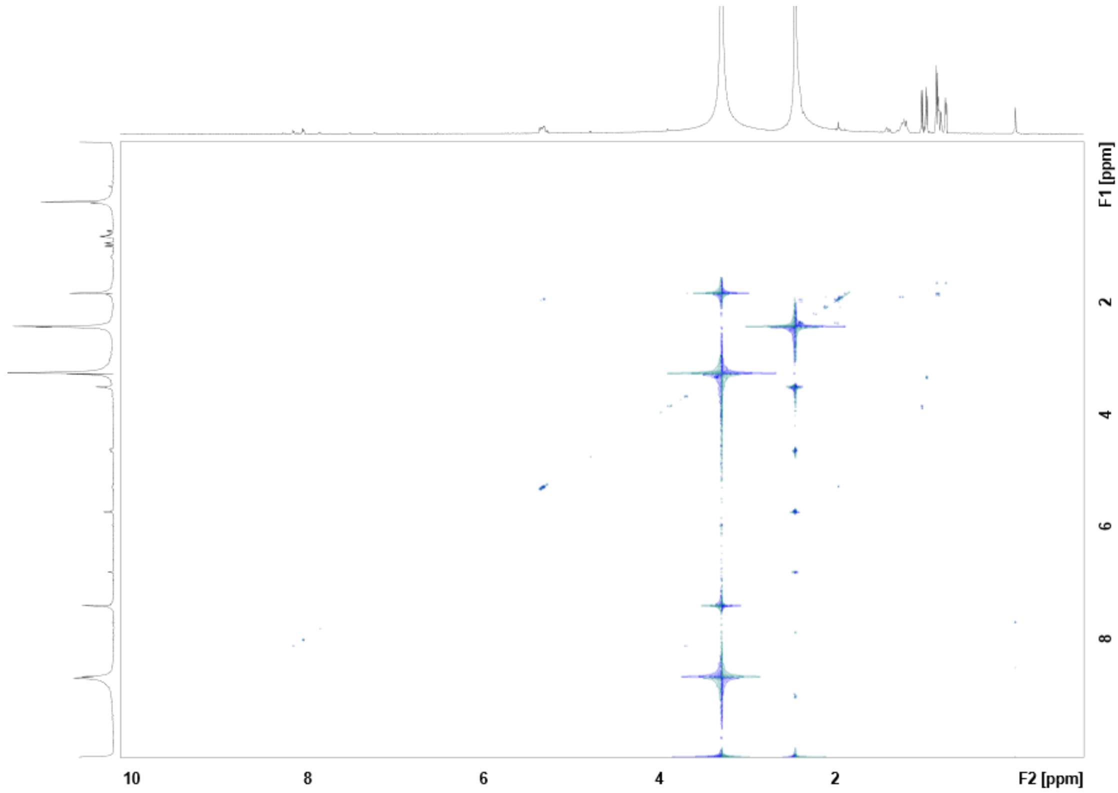

**Figure S19.** COSY spectrum of chitinimine II in DMSO-d6.

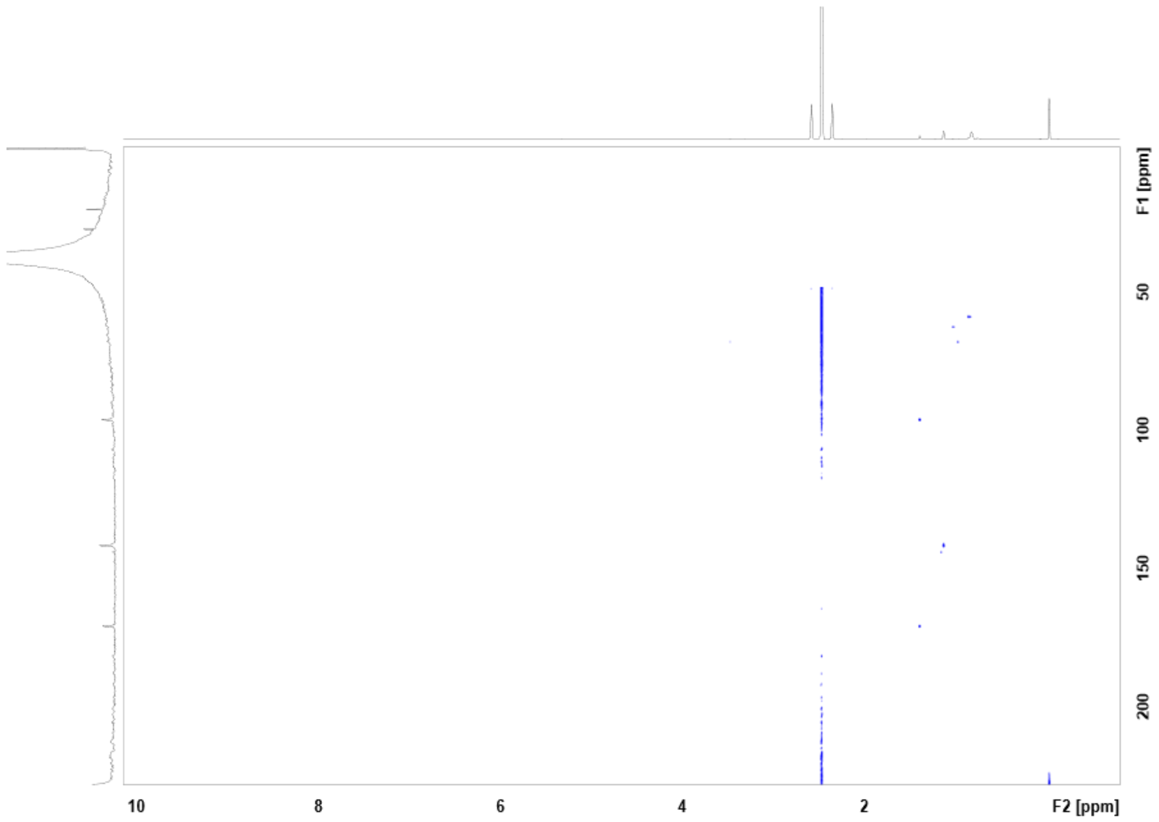

**Figure S20**. HMBC spectrum of chitinimine II in DMSO-d6.

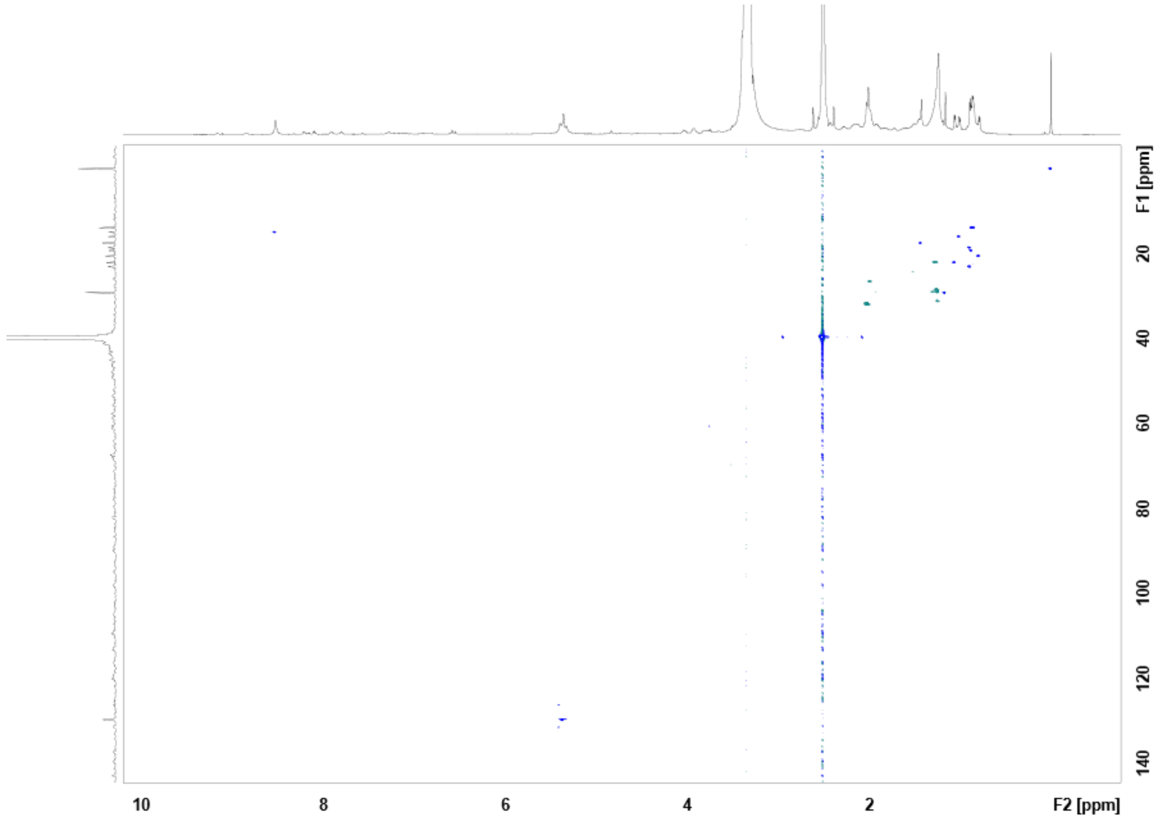

**Figure S21**. HSQC spectrum of chitinimine II in DMSO-d6.

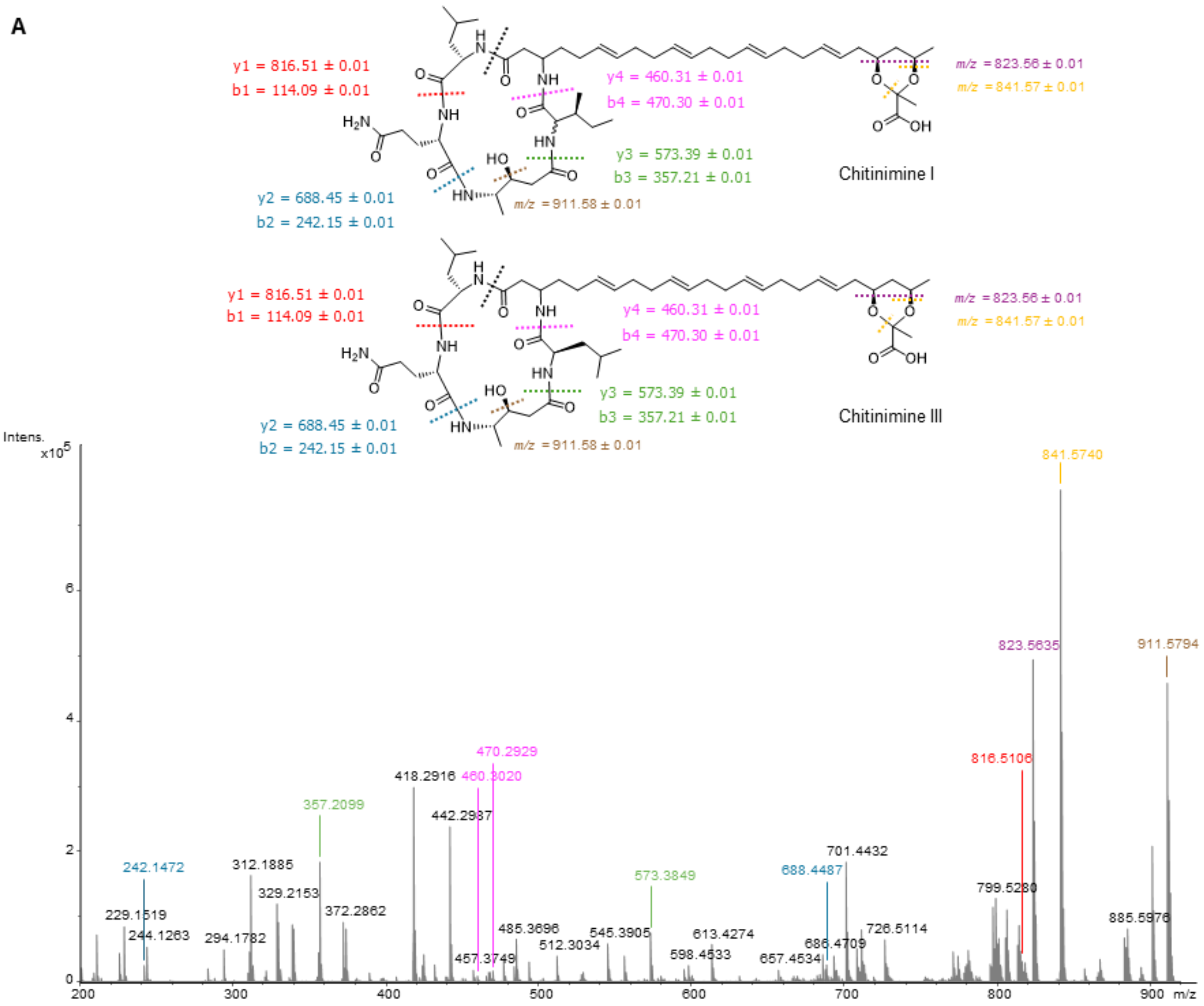

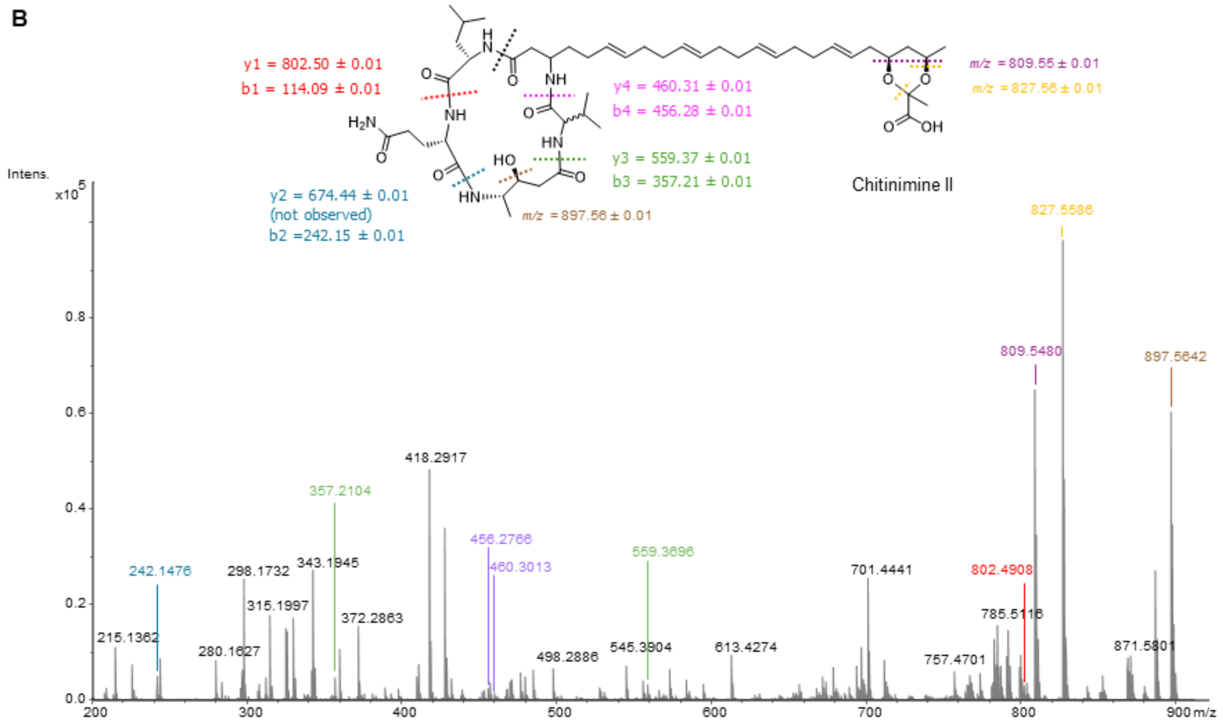

**Figure S22**. High-resolution LC-ESI-MS/MS spectra of (A) chitinimine I and III ($[M+H]^+$ = 929.5968 Da), and (B) chitinimine II ($[M+H]^+$ = 915.5815 Da). Selected b and y-fragment ions are indicated. Inset: structures of chitinimine I-III and detected fragment ions.

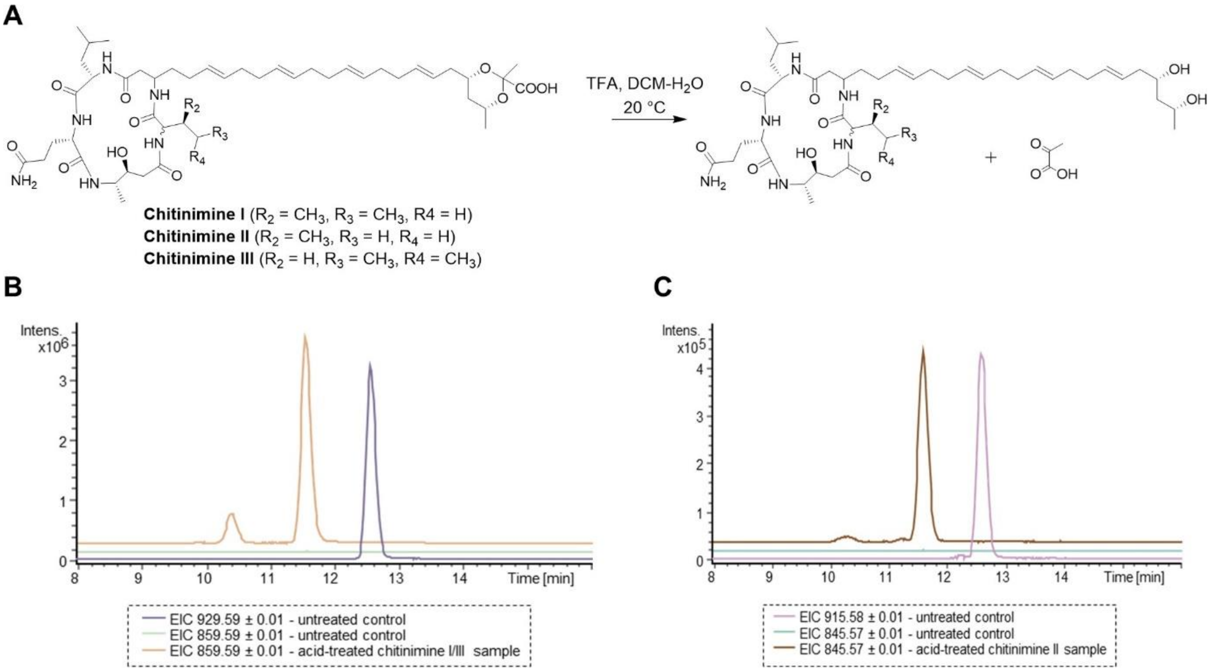

**Figure S23. Acid hydrolysis of the chitinimines.** A) Samples containing either chitinimine I/III or chitinimine II were hydrolyzed using TFA, releasing pyruvic acid. B) Extracted ion chromatograms at *m/z* = 929.5962 ± 0.01 (corresponding to the $[M+H]^+$ ion for chitinimine I/III) and *m/z* = 859.5902 ± 0.01 (corresponding to the $[M+H]^+$ ion for hydrolyzed chitinimine I/III) from UHPLC-ESI-Q-TOF-MS analyses of TFA-treated chitinimine I/III. C) Extracted ion chromatograms at *m/z* = 915.5879 ± 0.01 (corresponding to the $[M+H]^+$ ion for chitinimine II) and *m/z* = 845.5746 ± 0.01 (corresponding to the $[M+H]^+$ ion for hydrolyzed chitinimine II) from UHPLC-ESI-Q-TOF-MS analyses of TFA-treated chitinimine II.

A

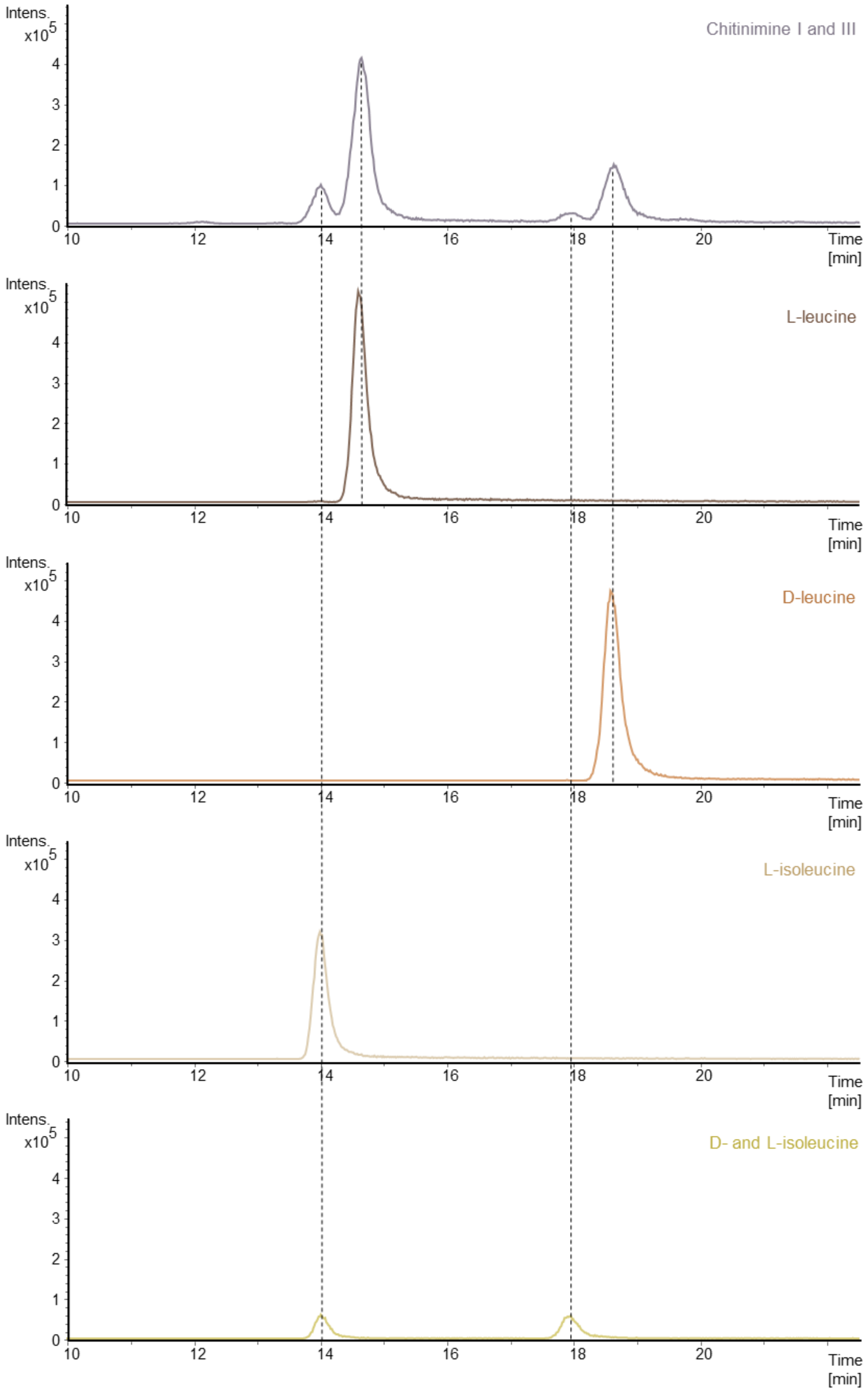

Intens. x10^5
Chitinimine I and III
L-leucine
D-leucine
L-isoleucine
D- and L-isoleucine
Time [min]

**B**

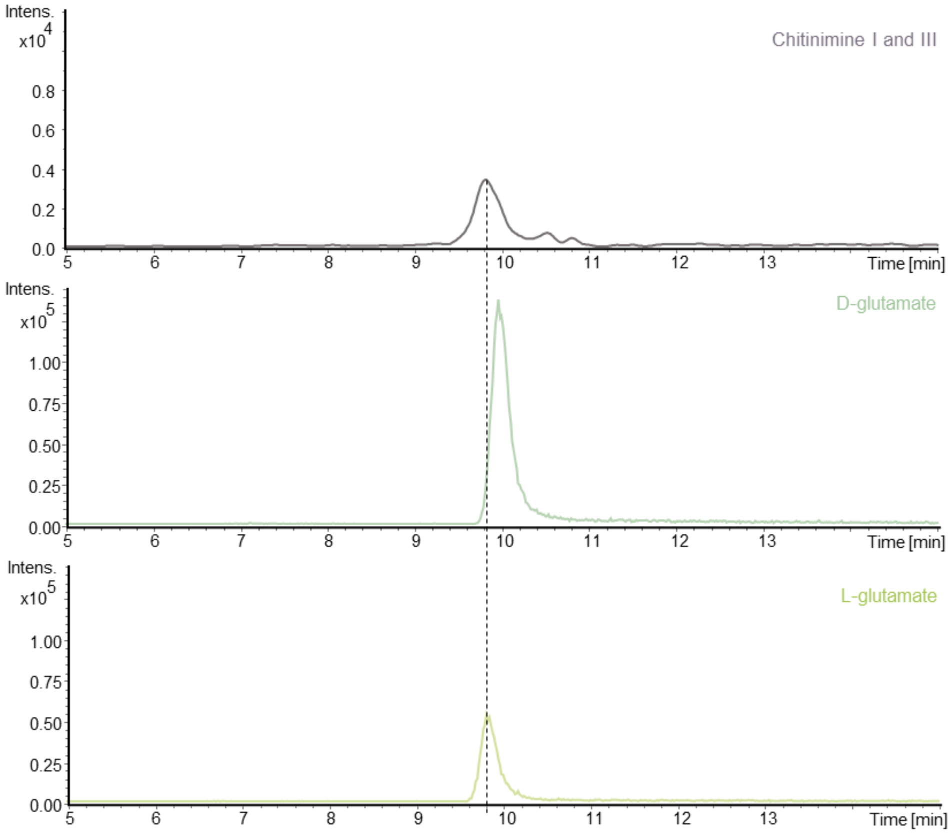
Intens.
x10^4
0.8
0.6
0.4
0.2
0.0
5 6 7 8 9 10 11 12 13 Time [min]
Chitinimine I and III
Intens.
x10^5
1.00
0.75
0.50
0.25
0.00
5 6 7 8 9 10 11 12 13 Time [min]
D-glutamate
Intens.
x10^5
1.00
0.75
0.50
0.25
0.00
5 6 7 8 9 10 11 12 13 Time [min]
L-glutamate

**C**

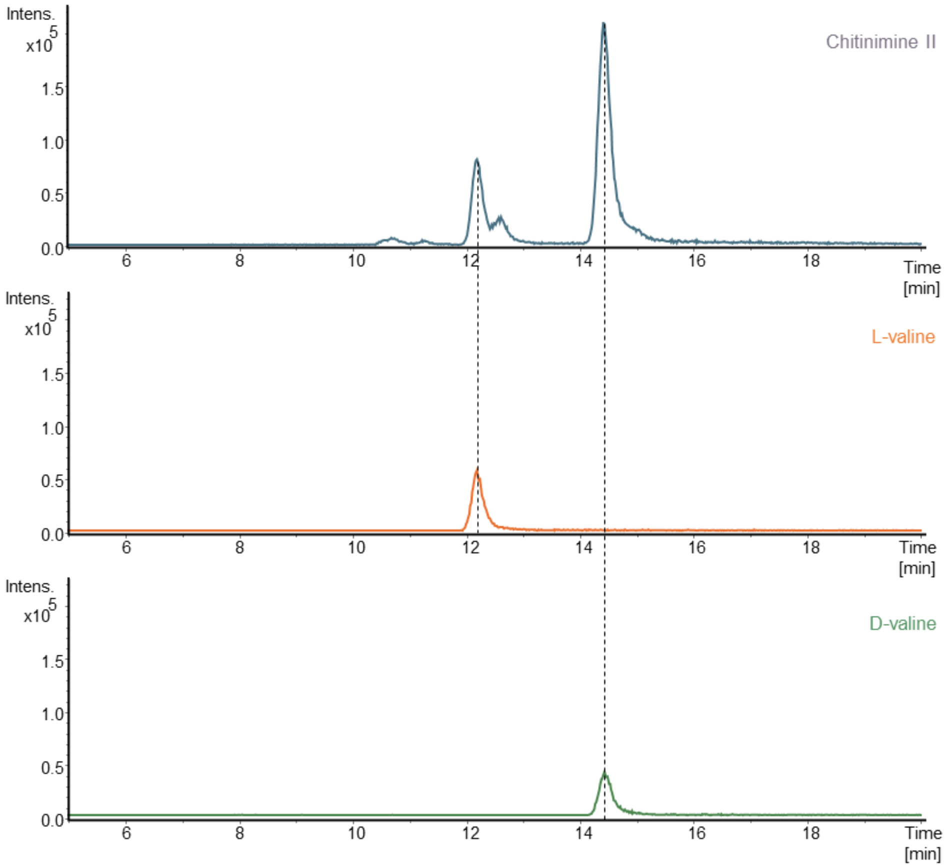
Intens.
x10^5
1.5
1.0
0.5
0.0
6 8 10 12 14 16 18 Time [min]
Chitinimine II
Intens.
x10^5
1.5
1.0
0.5
0.0
6 8 10 12 14 16 18 Time [min]
L-valine
Intens.
x10^5
1.5
1.0
0.5
0.0
6 8 10 12 14 16 18 Time [min]
D-valine

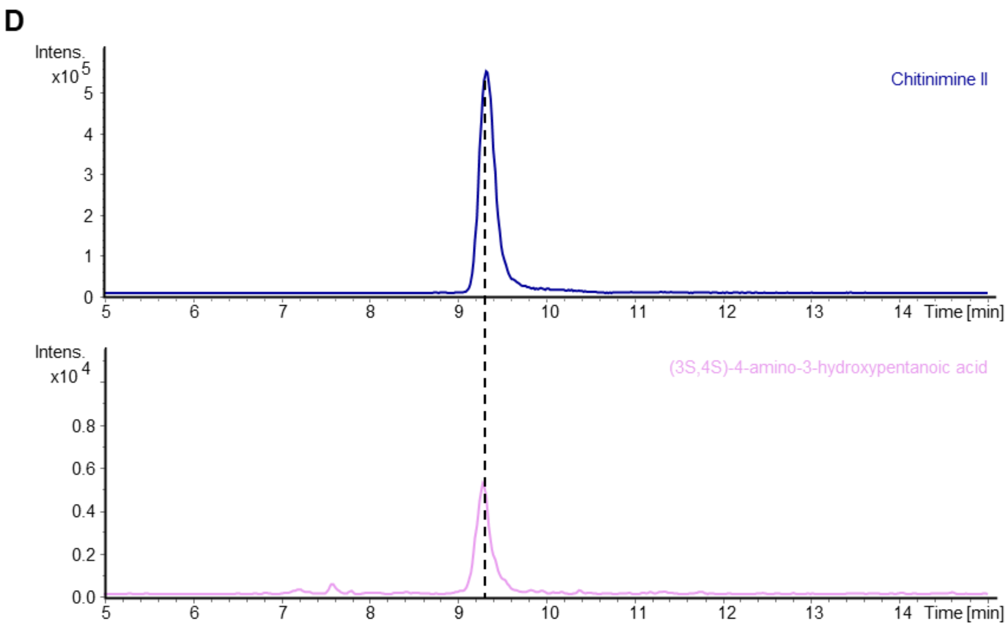

**Figure S24. Determination of the absolute stereochemistry of the chitinimine amino acids using Marfey's method.** (**A**) Extracted ion chromatograms at *m/z* = 384.1514 ± 0.01 (corresponding to $[M+H]^+$ for the Marfey's derivatives of Leu and Ile) from LC-MS analyses comparing the derivatives from chitinimine I and III hydrolysates with the derivatives of the authentic standards D/L-Leu and D/L-Ile. (**B**) Extracted ion chromatograms at *m/z* = 400.1099 ± 0.01 (corresponding to $[M+H]^+$ for the Marfey's derivative of Glu) from LC-MS analyses of Marfey's derivative of D-Glu and L-Glu and the derivative from the acid-hydrolysed chitinimine I and III. (C) Extracted ion chromatograms at *m/z* = 370.1357 ± 0.01 (corresponding to the $[M+H]^+$ for the Marfey's derivative of Val) from LC-MS analyses of Marfey's derivative of authentic standards of D-Val and L-Val, and Marfey's derivative from the acid-hydrolysed chitinimine II. (**D**) Extracted ion chromatograms at *m/z* = 386.1306 ± 0.01 (corresponding to the $[M+H]^+$ for the Marfey's derivative of 4-amino-3-hydroxypentanoic acid) from LC-MS analyses of Marfey's derivative of an authentic standard of (3S,4S)-4-amino-3-hydroxypentanoic acid and the derivative from the acid-hydrolysed chitinimine II.

```
ChtnE_AT4      LAAGVGPLLPGLVDALCANHRDAEADTRALYPRQLAPLAAEQ-----------------R 43
ChtnD_AT3      LFAGQGAQTVGMGARLCIDHPAARACFEA----FDAAGRAQGSAPISALVFPPDTFDAGR 56
ChtnA_AT1      --PGQGNPQPGAAAALYRDHAGFRAAIDR----CAALLDELLDVPLATLLFD-----A-- 47
ChtnB_A2       -FPGQGSQRAGMGHAIYAAEPAFRALADA----CLALLPAELAAQVRLAAFADPAEAG-- 53
                  * *    *    :   .   .*          *

ChtnE_AT4      AAIEEALLADNQALIKAGVIASWQYTRLLTERIGLAPAARFGHSLGQASMLFAAGAWLPD 103
ChtnD_AT3      AEAQRAALTQTQHAQAAIGAYNMALYGVL-RSAGFAPDMALGHSFGELSALWAAGALDDA 115
ChtnA_AT1      -ERGGPLLRQTRYAQPAQFALGWALAQLW-AGWGVRPAALAGHSLGEYVAAVLAGMAPLE 105
ChtnB_A2       -AEAEALLARTEVAQPALFVHQYALARLL-LDWGLRPAGLLGHSLGEIVAAALAGVLTPP 111
                      *  ..    *          :     *. *    ***:*:      **

ChtnE_AT4      DGWMQRLAALG----DSLGRLSGELPAVRAAWNLAEGEPLDWANLLVLAPAERVLAAARS 159
ChtnD_AT3      -GYRAAVLARGSALTPPAGREVGGLIAV-------------------SAPAEQVSALLPQ 155
ChtnA_AT1      -QVLPLVARRGALMQETAD--GAAMLLV-------------------RAGAADVAALVEA 143
ChtnB_A2       -DAIQLVVRRARLMQDSAE--GAMLQAE--------------------LPA---AELAAL 145
                     :   .           . :                         *

ChtnE_AT4      E-PRAYVSLVNSPAEATLVGDRAACGRILAAIGA---EAVPAPDSLVMHCAPAAGERAAI 215
ChtnD_AT3      L-PGLALANLNSPRQTVAGGSAAAVEAALPRLAEAGLQAVRLPVAAAFHTGLV-DYAAAP 213
ChtnA_AT1      RPAELALAADNGPASCVLAGTAAAIEAAAAGLAGRGIRSRLLDVAAAFHSPLM-EPIVLR 202
ChtnB_A2       LPDGLAIAACNGPALTVAAGPVAQVEAFAAQLDARGVAWQRLRGRHAFHSPAM-APAAAA 204
                     ::  *.*   .  *  *        :              .:*        .

ChtnE_AT4      AARFTAPLGPRPEGLEFAGGVPAEWSPA---ALAERVAADLVAPLDFPALVERVYGRGVR 272
ChtnD_AT3      WQAALAGLPLAAPRLPVWANVSAQPYPADAAGIRALLARQPFEPVRFCEQVEAAYAAGGR 273
ChtnA_AT1      LAGLAAMVDWQAGGVPAIANLHGRRH--AGAPDAAYWAAHARGTVRYREGLEALLADGHR 260
ChtnB_A2       LRPWLATLTAHPPQWPLLSNLDGGWMSDAEAVDPQRWARQLCAPVQFAPALAELAGRPGA 264
                    * :          ..: .              * .    : :   :    .

ChtnE_AT4      LFLEIGPGANCSRWIA--------         288
```

```
ChtnD_AT3       IFVEIGPRGILSRLVGDILG----          293
ChtnA_AT1       LFLELGPRPALATLGPTLAGGAEA          284
ChtnB_A2        LLLEIGAGTTLAAFARQA------          282
                :::*:*     :

ChtnE_AT4       ------------------------------------------------------------ 0
ChtnD_AT3       LFAGQGAQTVGMGARLCIDHPAARACFEAFDAAGRAQGSAPISALVFPPDTFDAGRAEAQ 60
ChtnA_AT1       --PGQGNPQPGAAAALYRDHAGFRAAIDRCAALLDELLDVPLATLLFD-----A---ERG 50
ChtnB_AT2       -FPGQGSQRAGMGHAIYAAEPAFRALADACLALLPAELAAQVRLAAFADPAEAG---AEA 56
```

**Figure S25. Sequence alignment of acyltransferase (AT) domains from the chitinimines BGC.** The conserved catalytic triad GxSxG is highlighted in yellow, substrate specificity determinants in blue.[28–31]

```
TropR-I_jweed     MEESKVSMMNCNNEGRWSLKGTTALVTGGSKGIGYAIVEELAG-LG-ARVYTCSRN-EK-      56
TropR-II_jweed    ------------MAGRWNLEGCTALVTGGSRGIGYGIVEELAS-LG-ASVYTCSRN-QK-      44
GlcDh_bacme       --------------MYKDLEGKVVVITGSSTGLGKSMAIRFAT-EK-AKVVVNYRSKED-      43
KR-fas_brana      --------------------SPVVVVTGASRGIGKAIALSLGK-AG-CKVLVNYARSAK-      37
KR-fas_ecoli      ----------------MNFEGKIALVTGASRGIGRAIAETLAA-RG-AKVIGTATS-EN-      40
KR1_ave           ---------------------GTTLITGGTGALATHLTHHLTTHQPTQHLLLTSRTGPHT      39
KR1_ery           ---------------------GTVLVTGGTGGVGGQIARWLAR-RGAPHLLLVSRSGPDA      38
KR2_ery           ---------------------GTILVTGGTAGLGAEVARWLAG-RGAEHLALVSRRGPDT      38
ChtnD_KR          ----------------------LVLVTGGARGVTARCIEALAA-RVPARFVLIGRSAPMA      37
ChtnB_KR          ----------------------VTLITGGFGRVGQAFARRLAA-RPGARLVLLGRQVPAG      37
                                          ::**.   :       :         .

TropR-I_jweed     ----------------------------------------------ELDECLEIWREKGL      70
TropR-II_jweed    ----------------------------------------------ELNDCLTQWRSKGF      58
GlcDh_bacme       ----------------------------------------------EANSVLEEIKKVGG      57
KR-fas_brana      ----------------------------------------------AAEEVSKQIEAYGG      51
KR-fas_ecoli      ----------------------------------------------GAQAISDYLGA---      51
KR1_ave           PHAQHLT--T------------------------------------------QLQQKGI       54
KR1_ery           DGAGELV--A------------------------------------------ELEALGA       53
KR2_ery           EGVGDLT--A------------------------------------------ELTRLGA       53
ChtnD_KR          ADPAWAAGIAEPAALRARALQQLREEGTAPTPRLVEARCQAVLAGREVRQTLQTLAAHGA      97
ChtnB_KR          DDPRLLELRA-----------------------------------------------LGA      50

TropR-I_jweed     NVEGSVCDLLSRTERDKLMQTVAHVFDGKLNILVNNAGVVI---HKEAKDFTEKDYNIIM      127
TropR-II_jweed    KVEASVCDLSSRSERQELMNTVANHFHGKLNILVNNAGIVI---YKEAKDYTVEDYSLIM      115
GlcDh_bacme       EAIAVKGDVTVESDVINLVQSAIKE-FGKLDVMINNAGLEN---PVSSHEMSLSDWNKVI      113
KR-fas_brana      QAITFGGDVSKEADVEAMMKTAIDA-WGTIDVVVNNAGITR---DTLLIRMKKSQWDEVI      107
KR-fas_ecoli      NGKGLMLNVTDPASIESVLEKIRAE-FGEVDILVNNAGITR---DNLLMRMKDEEWNDII      107
KR1_ave           HLTITTCDTSTPRPTHNNSLNTIPP-QHPVTTVIHTGGILD---DATLTNLTPTQLNNVL      110
KR1_ery           RTTVAACDVTDRESVRELLGGIGDD-VPL-SAVFHAAATLD---DGTVDTLTGERIERAS      108
KR2_ery           RVSVHACDVSSREPVRELVHGLIEQ-GDVVRGVVHAAGLPQ---QVAINDMDEAAFDEVV      109
ChtnD_KR          QADYLPLDLGDAAATRAAIAALTAR-HGRVAALVHGAGALA---DRRIADKTAQDIETVF      153
ChtnB_KR          EVLALAGDIAADGVAHAAVQAALGR-FGRLDCVIHAAGVAGEAAHRPLLECGRAERAAIQ      109
                  .      :                        :.: ..

TropR-I_jweed     GTNFEAAYHLSQIAYPLL-KASQNGNVIFLSSIAGFSALPSVSLYSASKGAINQMTKSLA      186
TropR-II_jweed    SINFEAAYHLSVLAHPFL-KASERGNVVFISSVSGALAVPYEAVYGATKGAMDQLTRCLA      174
GlcDh_bacme       DTNLTGAFLGSREAIKYFVENDIKGTVINMSSVHEKIPWPLFVHYAASKGGMKLMTETLA      173
KR-fas_brana      DLNLTGVFLCTQAATKIMMK-KRKGRIINIASVVGLIGNIGQANYAAAKAGVIGFSKTAA      166
KR-fas_ecoli      ETNLSSVFRLSKAVMRAMMK-KRHGRIITIGSVVGTMGNGGQANYAAAKAGLIGFSKSLA      166
KR1_ave           RAKAHSAHLLHQLTQHTP-----LTAFVLYSSAAATFGAPGQANYAAANAYLDALAHHR-      164
KR1_ery           RAKVLGARNLHELTRELD-----LTAFVLFSSFASAFGAPGLGGYAPGNAYLDGLAQQR-      162
KR2_ery           AAKAGGAVHLDELCS--D-----AELFLLFSSGAGVWGSARQGAYAAGNAFLDAFARHR-      161
ChtnD_KR          RPKLDGLLTLLEALDPAP-----PARVLLFSSTAGFSGNAGQADYAMANEALAKLAFQLP      208
ChtnB_KR          AAKLDGTRRLAAALDGVA-----VRRVLLCSSLSTVLGGLGFAAYAAGNRALEVLAERQ-      163
                    :  .                  .:  .*            *.  :  :  ::
```

```
TropR-I_jweed       CEWAKDNIRVNSVAPGVILTPLVETAIKKNPHQKEEIDNFIVKTPMGRAGKPQEVSALIA      246
TropR-II_jweed      FEWAKDNIRVNGVGPGVIATSLVEMTIQ-DPEQKENLNKLIDRCALRRMGEPKELAAMVA      233
GlcDh_bacme         LEYAPKGIRVNNIGPGAINTPINAEKFA-DPEQRAD---VESMIPMGYIGEPEEIAAVAA      229
KR-fas_brana        REGASRNINVNVVCPGFIASDMTAKL---GEDMEKK---ILGTIPLGRTGQPENVAGLVE      220
KR-fas_ecoli        REVASRGITVNVVAPGFIETDMTRAL---SDDQRAG---ILAQVPAGRLGGAQEIANAVA      220
KR1_ave             -HT--HHLPATSIAWGTWQG----------------------------------------      181
KR1_ery             -RS--DGLPATAVAWGTWA-----------------------------------------      178
KR2_ery             -RG--RGLPATSVAWGLWA-----------------------------------------      177
ChtnD_KR            LRW--RGVRAVALAWGPWA-----------------------------------------      225
ChtnB_KR            -SR--DGAQWLALGYDGW------------------------------------------      178
                                :  .

TropR-I_jweed       FLCF-PAASYITGQIIWADGGFTANGGF-----    273
TropR-II_jweed      FLCF-PAASYVTGQIIYVDGGLMANCGF-----    260
GlcDh_bacme         WLAS-SEASYVTGITLFADGGMTQYPSFQAGRG    261
KR-fas_brana        FLALSPAASYITGQAFTIDGGIAI---------    244
KR-fas_ecoli        FLAS-DEAAYITGETLHVNGGMYMV--------    244
KR1_ave             ---------------------------------    181
KR1_ery             ---------------------------------    178
KR2_ery             ---------------------------------    177
ChtnD_KR            ---------------------------------    225
ChtnB_KR            ---------------------------------    178
```

**Figure S26. Sequence alignment of the ketoreductase (KR) domains from the chitinimine BGC with representative A- and B-type KR domains from other organisms.** B-type KR domains generate D-β-hydroxyl groups and their classification in *cis*-AT PKSs is based on the presence of a conserved Leu-Asp-Asp-like motif, which is highlighted in yellow.[32,33] The Rossman fold involved in NADP(H)-binding GGxG(xxG) is marked in blue, the catalytic triad SYN/K in pink. [34] As a reference; the following KR domains were used: TropR-I_jweed, tropinone reductase-I from jimsonweed (*Datura stramonium*) (type A); TropR-II_jweed, tropinone reductase-II from jimsonweed (type A); GlcDh_bacme, glucose dehydrogenase from *Bacillus megaterium* (type A); KR-fas_brana, β-keto ACP reductase from *Brassicanapus* (type B); KR-fas_ecoli, β-keto ACP reductase from *E. coli* (type B); KR1_ave from the avermectin BGC from *Streptomyces avermitilis* (type B); KR1_ery from the erythromycin BGC from *Saccharopolyspora erythraea* (type B); KR2_ery from the erythromycin BGC from *Saccharopolyspora erythraea* (type A).[35]

```
ChtnE_DH1      EARAADARLAAGAAAAQVEAAAAPREVLFDEADVMEFAEGRVANVLGPHYAPVDALPRRV 60
ChtnE_DH2      -------RNRRPPAAQPFAAPLAPPGRPLERADLNALARGAIAEVLSPAHAAGGR-NPAL 52
                         *     **  . *  **    ::.**:  :*.* :*:**.* :*  .     :

ChtnE_DH1      RIPGPPFMAVSRITHLSGTYGQLEGSRIRTEYDIPDNAWNAVDGQ--------------- 105
ChtnE_DH2      RIPPPVIQFIDRVVRIDAAGGACGLGRSEAEWRIDPQHWAIRAHFKDDPVFPGPCMLEGA 112
                  *** * :  :.*:.::..: *    .* .:*: *  : *

ChtnE_DH1      ASYLSLDAQGVLFLAGWLGIDFENRGNRAYRWLDAQLTYLGPMPRAGQRVEYDIHIHQA- 164
ChtnE_DH2      VQLLQLHALALGLQTAVAGARFQPVAGRPIL-----VRFRAQVVPRNQLFTYRADIVEIG 167
                  .. *.*.* .: : :.  *  *:  ..*        : : . :   .* . *  .* :

ChtnE_DH1      -----FRNGDATLF         173
ChtnE_DH2      LGPEPYLIADIDLI         181
                       :  .*  *:
```

**Figure S27. Sequence alignment of dehydratase (DH) domains from the chitinimine BGC.** In DH1, the active site aspartate residue is present (DxxxH), whereas it is absent in DH2. The catalytic histidine in DH1 is located within an HxxxVxxxP instead of HxxxGxxxxP motif, while in DH2 it is included in a HxxxGxxP motif.[36]

```
ChtnE_KS5      ------------------------------------------------------------ 0
ChtnE_KS4      --IIGLGCLVPDAGDPATFWANLCAGRRSIRDADARDWGVEPTRFLAPGRGVADHVSSLE 58
ChtnD_KS3      IAVIGMAAMLPKAHDLAEYWRNIVDGTDCLEPLPADRWRSEHYYDADPKA---ADRAYAE 57
ChtnA_KS1      IAIIGAACRFPGADSPDRLAELLFDGREAIGPVAALR---PA-----------IAAGGIE 46
ChtnB_KS2      IAVVGMAGRFPGAADVEALWQLLLEGRSGVREIGRDEALADG---ADPAL--LDHPGYVP 55

ChtnE_KS5      ---------AIDALRLRVPPNDIDRMYPQQLLMLAVGDAALRDAGIEPGSR--------- 42
ChtnE_KS4      LG--KPRDFVFDPSGYLLPADFLAAQDRCIQWPLEAARQALLDAGLQPGE-------DLG 109
ChtnD_KS3      RGGF-VPDLWFDPLRYGMPPNTLASTDAAQLYALAVGRQALLDAGYDPDPAGTGRRLPAG 116
```

```
ChtnA_KS1      RAGLIAAPELFDPQFFGIAQREADQMDPQQRLALELAVEALEAAGLPRAG------LAGS 100
ChtnB_KS2      FAGTLDGIDAFDERLFGYSPADAALIDPQGRIFLECAHEALASAGIDPAR-------CGG 108
                            :*                     *  .  **  **

ChtnE_KS5      -TAVIVAGAMDHA-------GHRLMARWEAAWRLEDNLDAAGFDLSAEERTQLAALVRDA 94
ChtnE_KS4      RVGLVLGSYAWAAGSASDALTR---PLYDQALAQ------AFA---EAAGDPDRDPLRLT 157
ChtnD_KS3      RAGIVLGVSGNTMKLSSEMGKRADIAKWIDALRQ------AGA---GAA---LIETVAGA 164
ChtnA_KS1      RTGVYLGISTYDYSRLQMRRGD-----------------------GGE---L------- 126
ChtnB_KS2      RIGVYAGASVSSYALAALRGPA-------------------LA---DTE---L------- 136
                    .:  .

ChtnE_KS5      LHQPV---DAVVMLSYVGSLLASRIAATWDLSGPAFMLTGDETGALRALDLGAKLLASDE 151
ChtnE_KS4      VGRPTP--STHPESARISGGITTTVARALGLGGPRYAIDAACATSLYAIHLAALHLAAGE 215
ChtnD_KS3      MRRHYPDWTEDTFPGFLANLVAGRIANRFGLGAASHTVDAACASSLAAVRLACLELRSGA 224
ChtnA_KS1      ----Y------AGTGNAFSIAANRISYWLNLAGPSMAVDTACSSSLTAVHLAVRALRAGE 176
ChtnB_KS2      ----FR-----ALFANDKDYLASRVAYKLGLKGPAVGVQTACSTSLVAVAMAVRALRSGE 187
                                .   .  :  ::   .* .    :    : :* *: :.   * :.

ChtnE_KS5      ADAVLVGAVDLAGAIENLMVRQAQG-------------VDRAAPVGEGAVAVVLEPAAAV 198
ChtnE_KS4      ADAMLVVAANAFDTLYATFGFAATQALPDGSPNRPFDARSDGVAPADGAVALVLRRSGSR 275
ChtnD_KS3      ADLMLTGGVDTDNSNVAFLSFSKTPALSRSGRVRAFDAAADGTMISEGVGMLVLKRLDDA 284
ChtnA_KS1      IDLALVGGVGLLLSGELMQVFAGAGMLAPDGRCKTFDAAADGYVRGEGGGMVVLRRAAEA 236
ChtnB_KS2      CDAVLAGGASVSVPQRVGYLYEDGSILSPDGRCRAFGIDAAGTVPGNGVGVVVLKRLSRA 247
                   *  *. ...                               .   .:*   :**.

ChtnE_KS5      -------------RERGGAAYAEWRGAGFGEQP--------------AEAARQAHAATGI 231
ChtnE_KS4      GHERPGLSGREGPDLCGREAYGVIRAIGLSSDGRG-QTLTAPNPKGQQLACERAYARSGI 334
ChtnD_KS3      -------------LAAGDRVYGLIRGLGASTDGAG-GAIFAPHAAGQARALEAAYADAGI 330
ChtnA_KS1      -------------AAAGDRVLALIAGSAVNQDGR-SNGLTAPSGPAQSAVLRAALADAGL 282
ChtnB_KS2      -------------LADGDPIRAVIRAVALNNDGADKVGFSAPSVGGQEEVLQAALREAGL 294
                                  *    .   . . . :                 . . *   :*:

ChtnE_KS5      AAAEPGLVEAGDTLPA---AAEL------------AGQPALASAAAVFGHARMAAPLLAA 276
ChtnE_KS4      DPASVAYVECHATGTKLGDRVELETVGRVFGP-----GQPVGSVKSNVGHLLTAAGVAGL 389
ChtnD_KS3      DPASVGLIEAHATGTVVGDAVEIESLESVLGG--AAAPVALGSVKAQIGHAKAAAGAASL 388
ChtnA_KS1      APAEVDAVELHGTGTPLGDPIEAQALGEVY-AAGRAAPLAVGSIKTNIGHLEAAAGIAGL 341
ChtnB_KS2      DAADIGYVETHGTGTRLGDEVELSALAGAFGGAGQGARCLIGSLKSNLGHLDAAAGVAGL 354
                    *.   :*   *        *                  :.*  : .**   **   .

ChtnE_KS5      LHAALALNARQLPAWAGWREAAAAHSLDGRAGYVPTEPRPWLRPRHGCRIAAVLARDGDG 336
ChtnE_KS4      VKTLFALREGMIPATVGIGQSLAAEIAGQ--PRILTEPTPWP---GAQRRAAVNAFGFGG 444
ChtnD_KS3      IKTLLALYHKVIPPTLGVSAPNPRLDPRERPFYLPRRARPWLAPAAGPRRAGVSVFGFGG 448
ChtnA_KS1      IKAALALAARRLPPSLNFSRPNPDIDLAALGLAVATEAVPLD-PIGRPARVGVSSFGFGG 400
ChtnB_KS2      IKAVLTVERGIVPASLHVEQPNALLAAGGSRFALATATVAWP-DDGRPRRAGVSSFGIGG 413
                  ::: :::    :*                    :                ..*   . .*

ChtnE_KS5      SAARALLAEAP   347
ChtnE_KS4      VNAHLVVD---   452
ChtnD_KS3      ANVHLALEE--   457
ChtnA_KS1      SNAHVVLEAA-   410
ChtnB_KS2      TNAHCIVEQP-   423
                    .:  :
```

**Figure S28. Sequence alignment of ketosynthase (KS) domains from the chitinimines BGC.** The catalytic triad highlighted in yellow is present in all KS domains, except in ChtnE_KS5, which is a chain-length factor.[37]

```
ChtnD_ACP6      --LLLQTVADKTGYPVEMLSLDMRLEGDLGVDSIKRVEILAAMRDALGLAAD—GAGDGL  56
ChtnD_ACP4      ---LLRTVADKTGFPVELLAPGMRLEGDLGVDSIKRVEILAALRDALGLAAADGQAGEAL 57
ChtnD_ACP3      ---LLRTVADKTGFPVELLTLEMKLEADLGVDSIKRVEILAAMRDALGLTAAAGD-GEAL 56
ChtnD_ACP5      ---LLRTVADKTGFPVELLTLEMKLEADLGVDSIKRVEILAAVRDALGLTTAAGD-GEAL 56
ChtnA_ACP1      ---VVEQIARALGEAPERLPLDKPF-IEMGADSVMMAEAMRAIQARYGVRISAR---QLL 53
ChtnF_ACP7      -DWLMAQVAAQLEVEADDIDPRRTF-ESYALDSARALLVLTRLEARLGLRLSPT---LIW 55
ChtnC_PCP4      -EALAAVWRAVL--QCGELALEDDF-YALGGDSIMAMQVSMRLTRQ-GWTLRPQ---DML 52
ChtnA_PCP1      ETALLDYVRGTL--QVALRGIDHDF-FAAGGQSLAATQLIGWVQRQWSVKPALK---DFF 54
ChtnB_PCP3      ERRIAAIWCEVL--GLPAVDAARNF-FEAGGNSLLLMQVHARLKQAFSPAPRLA---DLF 54
ChtnB_PCP2      EQAIAALWQALL--GVERVGRHDDF-FALGGHSLLATRAAARLGRRFGLRLPMA---ALF 54
ChtnC_PCP5      ELELARLWEDLL--GLAPIGRDDGF-FALGGHSLLVLELMARLRARFGRAVPFA---AFL 54
ChtnB_ACP2      EAAVAQAIADTL--ALAAVGPDDDF-FALGGDSLVATRVIARLRGATGLPLSVG---LVL 54
                      :                    :    . .*         :    .

ChtnD_ACP6      RGAVTLAELAER----      68
ChtnD_ACP4      RNAATLADIAA-----      68
ChtnD_ACP3      RGAATLGEIAAL----      68
ChtnD_ACP5      RGAATLGEIAALL---      69
ChtnA_ACP1      NELDTVDALSAHL---      66
ChtnF_ACP7      N-YPTIEALAGRLAQ-      69
ChtnC_PCP4      R-QPTLAAQCALMRR-      66
ChtnA_PCP1      E-LPTVACLAALIEA-      68
ChtnB_PCP3      R-FPSVAALAAFLGRE      69
ChtnB_PCP2      D-APVLAALAARIEAA      69
ChtnC_PCP5      R-APTVAGLAALLG--      67
ChtnB_ACP2      Q-APTVRLLAAAVR--      67
                        :   .
```

**Figure S29. Sequence alignment of acyl (ACP) and peptidyl carrier protein (PCP) domains from the chitinimine BGC.** The highly conserved 4’-phosphopantetheine binding motif (LGG(H/D)S(L/I)) is highlighted in yellow.[38]

```
ChtnC_E        --LPVTPIQAWFFALELAHPQ---HWNQAVRLALDP-AAAGRLEPALAALEAAHEALRLR 54
ChtnA_C1       ATLPLSDSQRQIWLATELDPAGAGAYCETVALEVDGELNPALLERALAQLALRHEALRTV 60
ChtnC_C5       DLLPVTPQQRGMLLESQAHAG-LGLHVEQFVATFDGAFDRAALEAAWARLVARHDTLRSG 59
ChtnC_C4       APVPLSIEQRAVWLAEQHGG--GRAFLIPGALRLRGRLDAAALRLALQALVDRHEAFRTA 58
ChtnB_C2       -----TPAQQRLWLLCGLGED-AANYVIAGALRLQGALDPARLEAALNDCLARHESLRTG 54
ChtnB_C3       -PAPLSASQLGLWFVQQLDPA-STAYVLSGAIRIDGALSRSLLSRTLDLLQARHHALRTR 58
                      :  *  .                     .      . *  :       *.::*

ChtnC_E       FAPGETGWTMRVAPAGAPPLRRVRAEDA------GEAL-VQIEAAQRSLDLAAGPVWRAL  107
ChtnA_C1      LDAGG--AGQTVQPALKPPLSYGE---------HDDVAGWLRGFVEAAFDPAAGGPLRAA  109
ChtnC_C5      FAWRQEHAPLCLVHARAEP-AWQHLDWRGEAVDEARIAAWLERDRLAGFDGARP-PLRFA  117
ChtnC_C4      IRLVGDEPMQCVQSAVHFALPEHDLSALMPAERDGALERLLAAEAARPFELARAPLLRAV  118
ChtnB_C2      FAEIDGVPQQAIEPQAALALPLTELDALPAEARLEAALATAGALARQPFDLARPPLLRAR  114
ChtnB_C3      FDAAGGVPSQTVLAPSGIALPIDDLSRLAPAMREAELQRRLDAEARRPFELPGPPPVRAR  118
                 :          :                                    ::       *

ChtnC_E       LIDGPHDGWPHLVLIAHHLVVDGVSWRLLVDDLAQALAGGE-----PAPAALGYADWAAH  162
ChtnA_C1      LLKRSDGRH-VLALRAHHVLVDGWSLALIVDELGRLYHGGA-----ALAEAQPFRRQLDW  163
ChtnC_C5      TLRLDASRW-LFVWTYHHALLDGWSVARLLAEALAPA---------ADEAPPAARDHARW  167
ChtnC_C4      LVRLAAEEH-VLALTPHHIVADGWSIDVMVRELSLFYRPDGSP-PPPLPAPLGYPDFAAW  176
ChtnB_C2      LFRLGADDW-LLALAIHHIVCDGPSLGILIADLAAAYAGRGQGRPLPAAPALQFADFAEW  173
ChtnB_C3      LLVLGPDCH-VLSLSLHHIVADGWSLGLVWRDLVQGYRALREGGAPDWTPLPQYADYARG  177
                  .         :    ** : ** *   :  :

ChtnC_E       LAGLPPQPPR--------AAVPPAPPPIARPDGDDFEAQTRI----------ATLALPMD  204
ChtnA_C1      LAGAQ----DEAAERFWRDRVAEPPAPLALPGQRLFDAA---ARPRWEGE-RVRLALPAE  215
ChtnC_C5      LAGQD----RAAAAAFWRAELAGAEVP--TPAGRVDP-----ALAPGRGHADRCRRIDAA  216
ChtnC_C4      QAGRIAAGADAADLDYWRATLAELPPPLELPAARQAAAGTDGIAAGYAGA-AVQATLPAA  235
ChtnB_C2      QQEQLARPETERLLAAAAARLAGVPD-LLVPTDRPRP-----ALRAGRGA-RHAFALDAV  226
ChtnB_C3      EAARLAAPAAAAELDYWRRALDALPP-LELPTDFARP-----PLPGYRGA-QYRFTVPAR  230
                                     :         *                         :

ChtnC_E       ATAALLGPANQPYRSEPTELLLAGLLLGLQAAHGRSALAVALERHGRDVDGVELAGTVGW  264
ChtnA_C1      L-AEKLAALAAGRRVTPFTAALALTLGWLHRLCDRDDILIGVPAHGRPD---GLERMVGQ  271
ChtnC_C5      T-VAQLETLARRHRLTPALLAQGLWGLALAWASGRREVTLARTVAGRPAEVDGSERWVGL  275
ChtnC_C4      T-ARGLRRLAAESGTTLFSALAALFAALLQRYGGRRELVLGTASAGRGRI--ELEEVVGL  292
ChtnB_C2      L-MDAVTLRARALGSTPFAVLLAAWGIVLAGWSGQDDFAIGTPVSGRADP--ALAEVVGL  283
ChtnB_C3      L-CGSADRLARAAGASRFMVLLAAFQALLARWSGQRDFAVGVPVSGREAL--DWQETVGC  287
                                       .     *    .:  . :.    **          **

ChtnC_E       FTAIVPLLLAL    275
ChtnA_C1      AVQLMPLRSRI    282
ChtnC_C5      FINSLPLRLSL    286
ChtnC_C4      FAGRLPLRLDF    303
ChtnB_C2      FAETAALRFRC    294
ChtnB_C3      FVNTVAIRAEL    298
                       :
```

**Figure S30. Sequence alignment of the condensation (C) domains from the chitinimine NRPS modules.** The active site HHxxxDG motif is highlighted in yellow and present in all C domains.[39] E domain-specific signature motifs are highlighted in green.

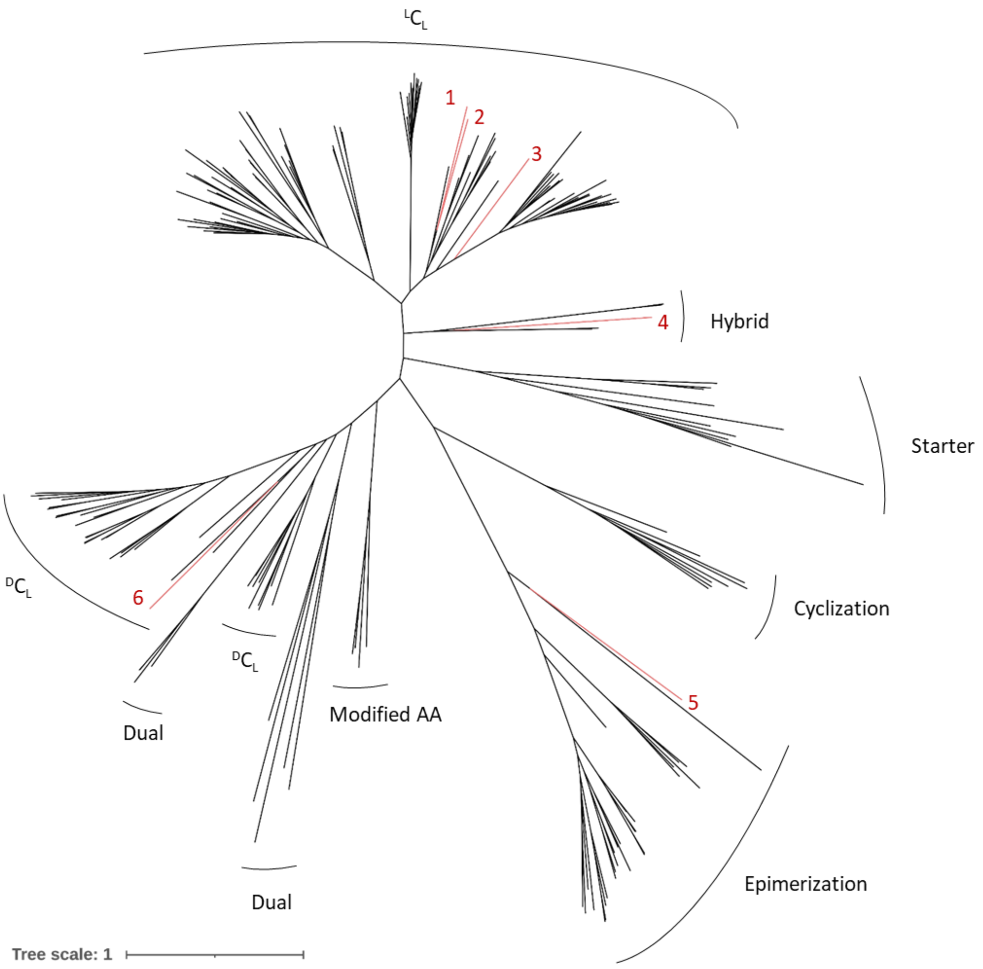

**Figure S31. Neighbor joining phylogenetic tree of condensation (C) domains from the chitinimine BGC and reference C domains from the NaPDoS database.** The phylogenetic tree was generated with NaPDoS2 and visualized with iTOL. Red branches represent C domains from the chitinimine BGC in *C. koreensis*. Functional classifications are based on NaPDoS annotations: 'Starter' C domains are typically not present in the first module of an NRPS, but when present, link an amino acid to a fatty acid, polyketide, or another molecule; '$^{L}C_{L}$' domains catalyze peptide bond formation between two L-configured amino acids; '$^{D}C_{L}$' domains couple a D-configured amino acid to an L-configured amino acid; 'Cyclization' domains catalyse chain elongation with cysteine, serine or threonine residues, followed by a two-step cyclodehydration reaction to form five-membered thiazoline or (methyl)oxazoline rings, respectively; 'Epimerization' domains act on elongated PCP-bound peptidyl intermediates and isomerize the $\alpha$-position of the C-terminal amino acid residue; 'Modified AA' domains modify the newly incorporated amino acid residue; 'Dual' domains have both condensation and epimerization abilities; 'Hybrid' domains add an amino acid to a growing polyketide chain. 1: ChtnB_C3 ($^{L}C_{L}$); 2: ChtnC_C4 ($^{L}C_{L}$); 3: ChtnB_C2 ($^{L}C_{L}$); 4: ChtnA_C1 (Hybrid: immediately downstream of a PKS module); 5: ChtnC_E (Epimerase); 6: ChtnC_C5($^{D}C_{L}$).

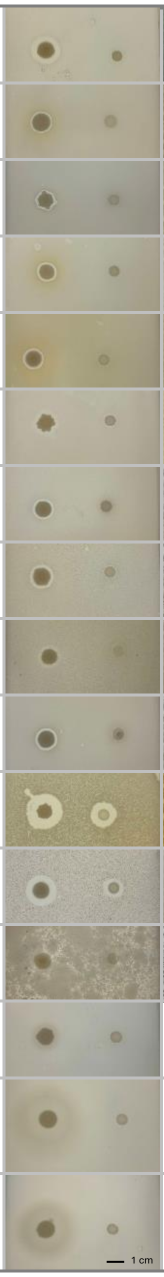

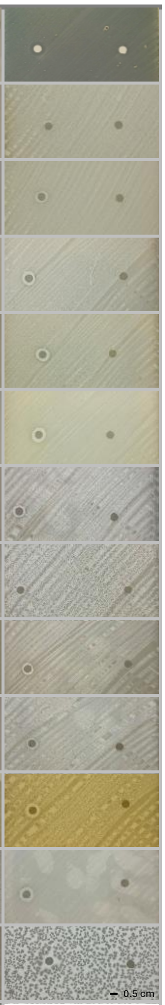

| | A | B |
|---|---|---|
| *Enterococcus faecium* DSM 25390 | | |
| *Staphylococcus aureus* DSM 21979 | | |
| *Staphylococcus aureus* RN4220 | | |
| *Staphylococcus aureus* ATCC 6538 | | |
| *Staphylococcus aureus* StaAu068 | | |
| *Staphylococcus aureus* Sa9 | | |
| *Staphylococcus capitis* StaCa010 | | |
| *Staphylococcus haemolyticus* StaHa024 | | |
| *Staphylococcus hominis* StaHo017 | | |
| *Staphylococcus lugdunensis* StaLu018 | | |
| *Bacillus cereus* DSM 31/ATCC 14579 | | |
| *Bacillus subtilis* ATCC 9799 | | |
| *Mycobacterium smegmatis* MC2-155 | | |
| *Salmonella newport* C487 | | No bioactivity |
| *Salmonella enterica* 14029 (+GFP) | | No bioactivity |
| *Salmonella enteritidis* ATCC 13046 | | No bioactivity |

**Figure S32. Antibacterial activity of the chitinimines and the *C. koreenis* DSM 17726 WT and *∆chtnA* mutant. A.** Soft agar halo assays comparing the antibacterial activity of the *C. koreensis* WT (left) and ∆*chtnA* mutant (right) strain. **B.** Plate lawn assays with 10 µL of a 1 mg/mL purified chitinimine mixture (left) and a DMSO solvent control (right).

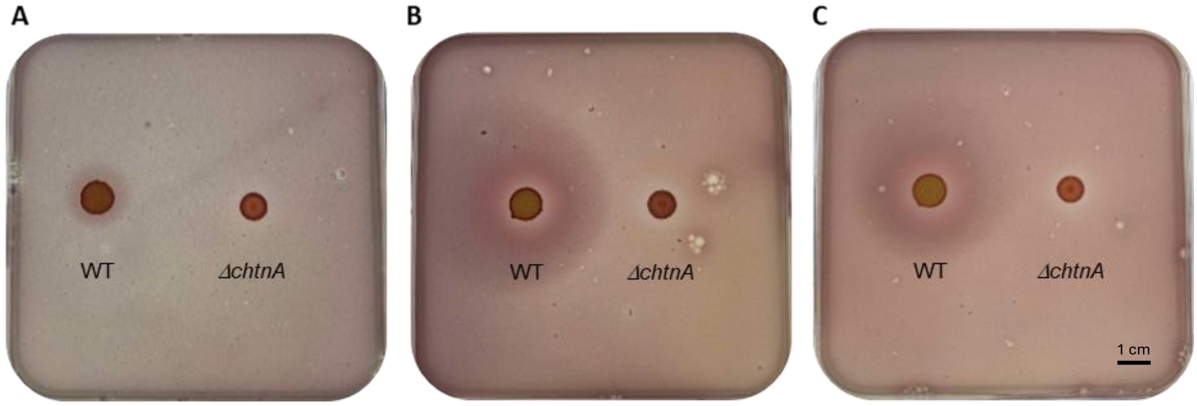

**Figure S33. Growth-promoting effects of *C. koreenis* DSM 17726 WT and the *ΔchtnA* mutant on *Salmonell*a species.** WT (left) and *ΔchtnA* mutant (right) cultures of *C. koreensis* were spotted onto BSM agar plates supplemented with glucose as a sole carbon source. Following incubation for four days at 28°C, the cells were inactivated by chloroform vapours and overlaid by soft agar inoculated with *Salmonella newport* C487 (A), S*almonella enterica* 14029 (B), or *Salmonella enteritidis* ATCC 13046 (C). The soft agar was also supplemented with iodonitrotetrazolium chloride to enhance visualisation of *Salmonella* growth. After 24 hours of incubation, growth-promoting effects were observed in the presence of WT *C. koreenis* DSM 17726.

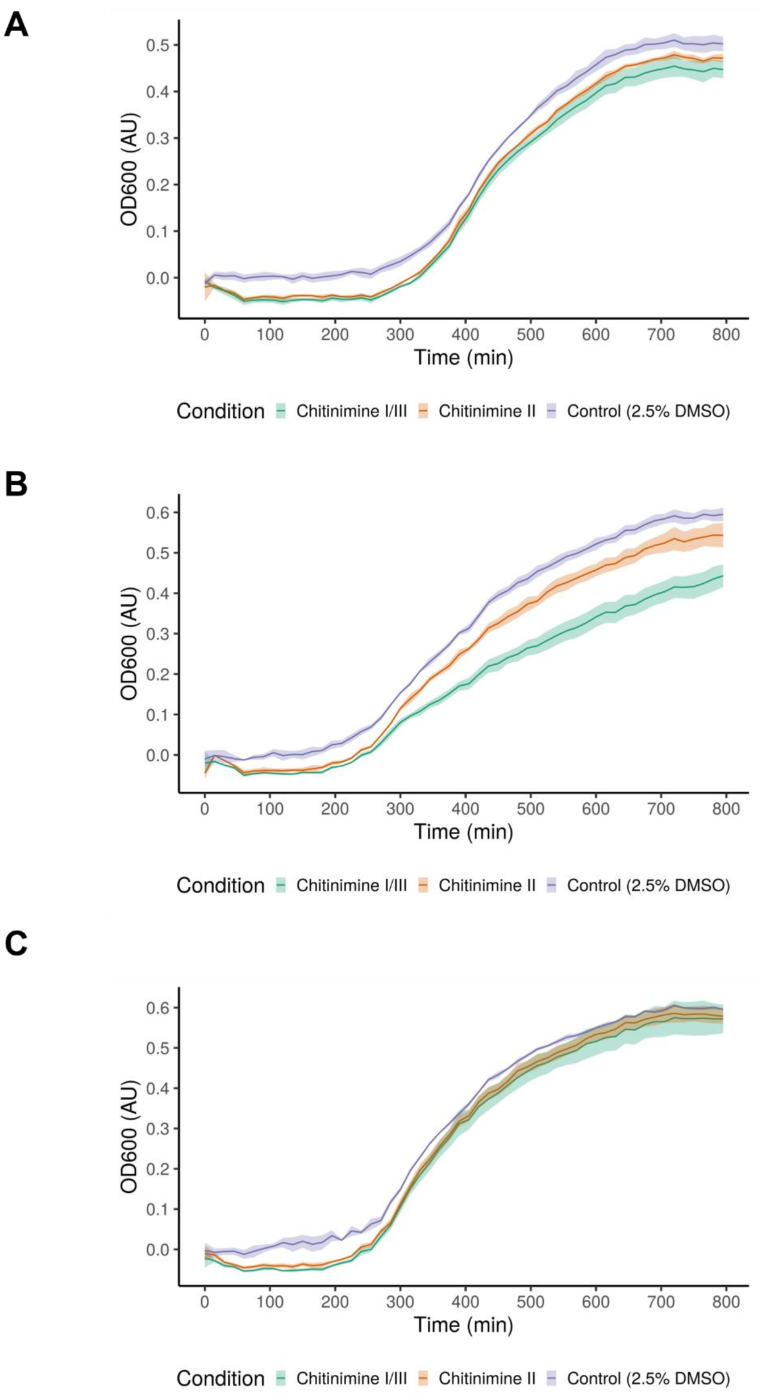

**Figure S34. Growth curves of *Salmonella* strains in the presence of chitinimines.** *Salmonella newport* C487 (A), *Salmonella enterica* 14029 (B), and *Salmonella enteritidis* ATCC 13046 (C) were exposed to chitinimine I/III or chitinimine II (125 µg/mL in 2.5% DMSO), and the optical density at 600 nm was monitored for 13 hours. No growth-promoting effect was observed compared to cultures supplemented only with 2.5% DMSO, indicating that the chitinimines alone do not have growth-promoting activity. Growth curves represent the mean of three replicates, while shaded areas represent standard deviations.

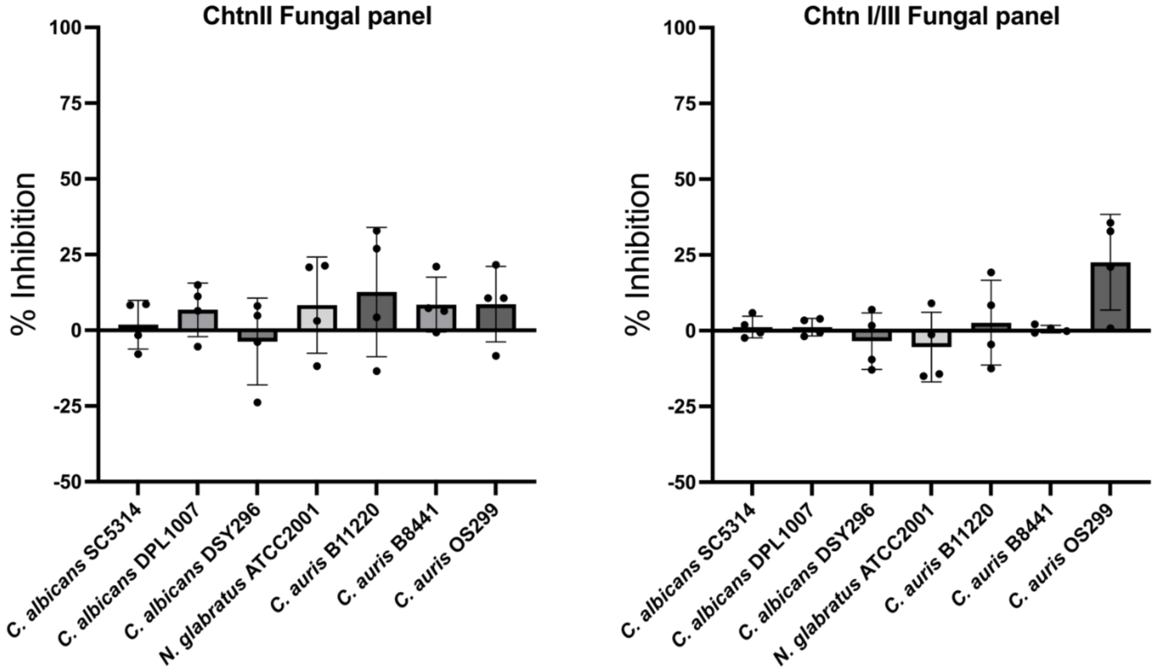

**Figure S35. Antifungal activity of the chitinimine II (left) and chitinimines I/III against a panel of *Candida albicans*, *C. glabrata* and *C. auris* strains.** Growth inhibition (%) was calculated relative to the untreated control for each strain.

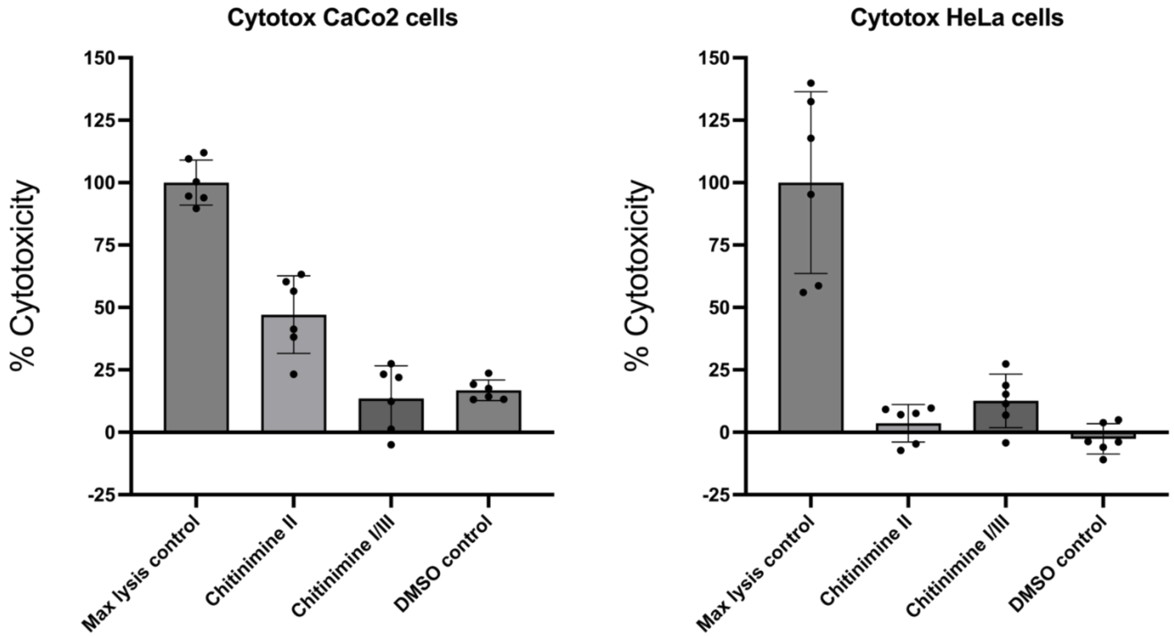

**Figure S36. Cytotoxicity of chitinimine I/III and chitinimine II against HeLa and CaCo-2 cells, as determined by the lactate dehydrogenase (LDH) cytotoxicity assay**. Maximum LDH release was obtained by lysing cells with the supplied lysis buffer (‘max lysis control’). Cells treated with 2% DMSO served as negative controls. Error bars represent the standard deviation of three independent experiments.

### Supplementary Tables

**Table S1**. **Summary of genome mining results for candidate clusters encoding hybrid PKS-NRPS-PUFA synthase-like biosynthetic pathways.** Compilation of NCBI GenBank accession numbers, species and strain names, bacterial orders, if the clusters were retained or discarded, and – if applicable – the reason for exclusion. For each entry, an image of the corresponding cluster-containing region from antiSMASH is included.

| GenBank accession no. | Species and strain | Order | Discarded? | Reason for exclusion |
|---|---|---|---|---|
| **Gram-negative bacteria** | | | | |
| AP009552.1 | *Microcystis aeruginosa* NIES-843 | Chroococcales | No | / |

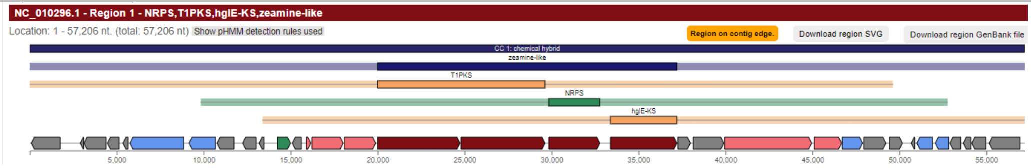

| GenBank accession no. | Species and strain | Order | Discarded? | Reason for exclusion |
|---|---|---|---|---|
| CP097576.1 | *Microcystis aeruginosa* Chao 1910 | Chroococcales | No | / |

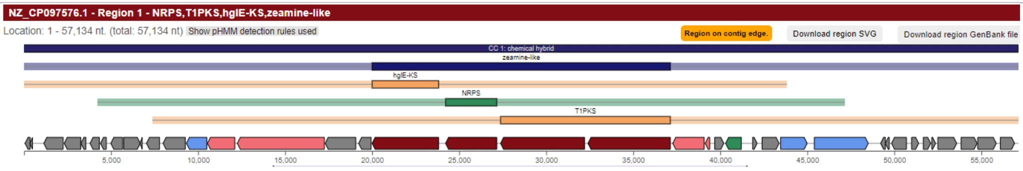

| GenBank accession no. | Species and strain | Order | Discarded? | Reason for exclusion |
|---|---|---|---|---|
| CP003552.1 | *Nostoc* sp. PCC 7524 (ATCC 29411) | Nostocales | Yes | PfaBC homolog misannotated as T1PKS |

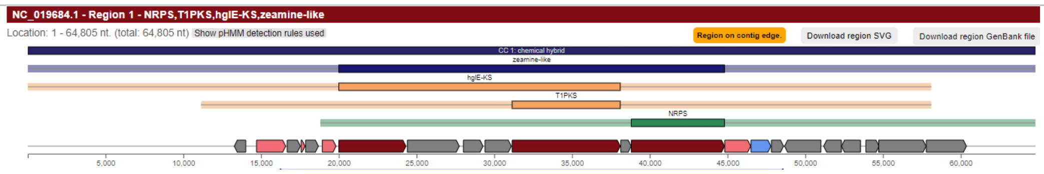

| GenBank accession no. | Species and strain | Order | Discarded? | Reason for exclusion |
|---|---|---|---|---|
| AP018216.1 | *Trichormus variabilis* NIES-23 | Nostocales | No | / |

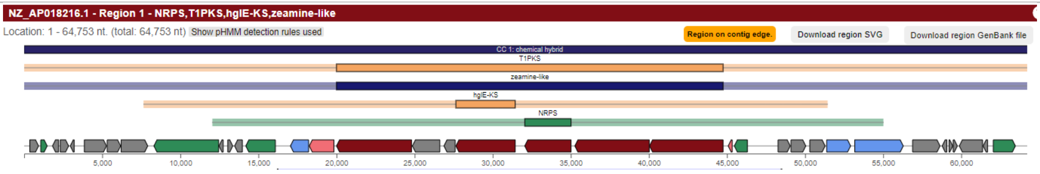

| GenBank accession no. | Species and strain | Order | Discarded? | Reason for exclusion |
|---|---|---|---|---|
| AP018316.1 | *Dolichospermum compactum* NIES-806 | Nostocales | Yes | 85% of genes from heterocyst glycolipid BGC share at least 30% identity to this *pfa* operon and 100% of genes from the anabaenopeptin NZ857/nostamide A BGC share at least 30% similarity to the NRPS genes. |

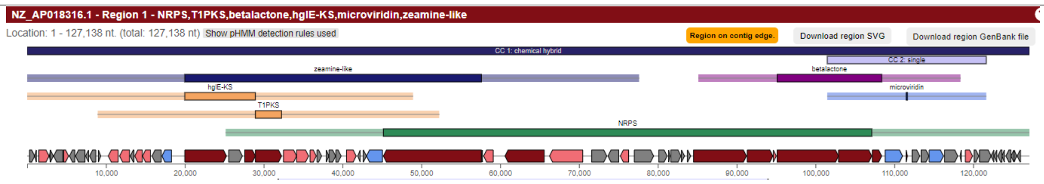

| GenBank accession no. | Species and strain | Order | Discarded? | Reason for exclusion |
|---|---|---|---|---|
| BDUD01000001.1 | *Nostoc commune* NIES-4072 | Nostocales | No | / |

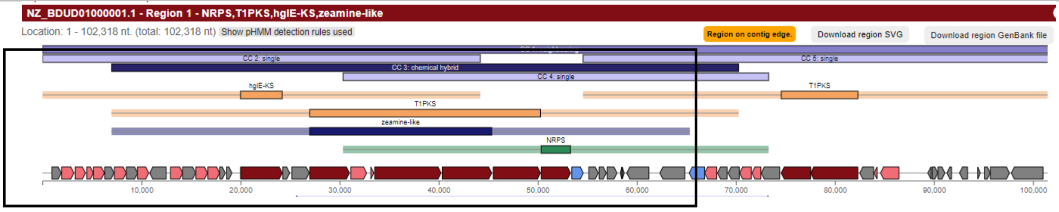

| GenBank accession no. | Species and strain | Order | Discarded? | Reason for exclusion |
|---|---|---|---|---|
| CP024792.1 | *Nostoc flagelliforme* CCNUN1 | Nostocales | No | / |

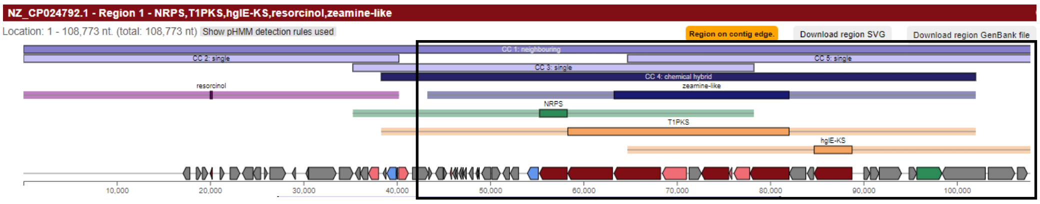

| | | | | |
|---|---|---|---|---|
| CP045227.1 | *Nostoc sphaeroides* CCNUC1 | Nostocales | No | / |

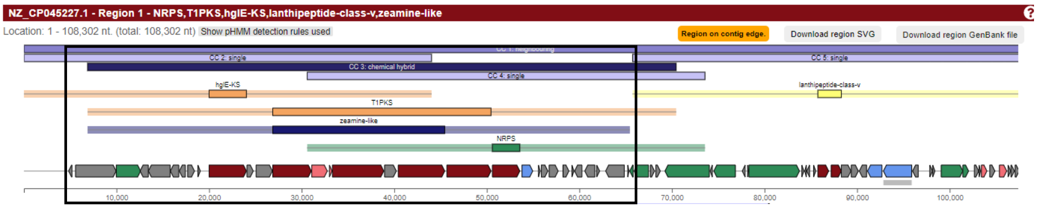

| | | | | |
|---|---|---|---|---|
| CP034058.1 | *Anabaena* sp. YBS01 | Nostocales | No | / |

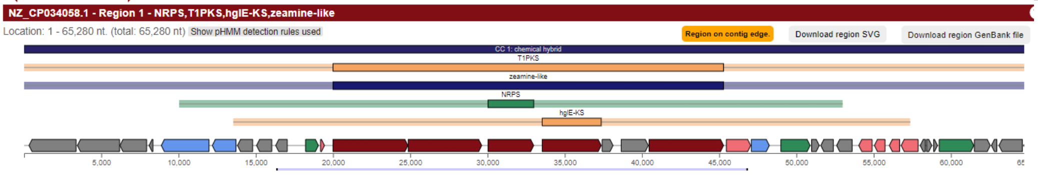

| | | | | |
|---|---|---|---|---|
| JH992901.1 | *Mastigocladopsis repens* PCC 10914 | Nostocales | Yes | 85% of genes from heterocyst glycolipid BGC share at least 30% identity to this *pfa* operon |

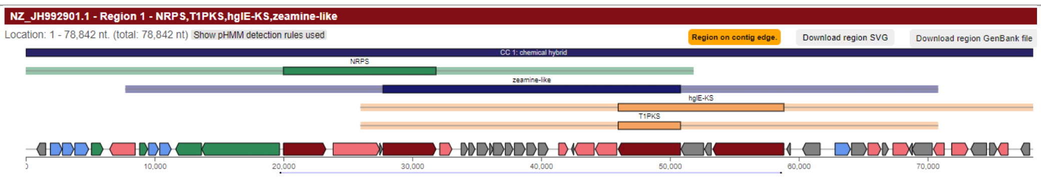

| | | | | |
|---|---|---|---|---|
| KQ976354.1 | *Scytonema hofmannii* PCC 7110 | Nostocales | No | / |

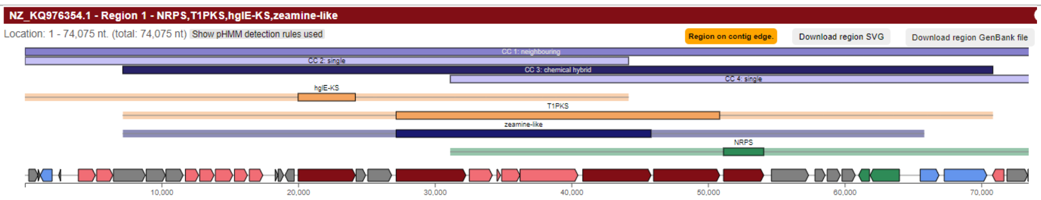

| | | | | |
|---|---|---|---|---|
| VILF01000001.1 | *Dolichospermum flos-aquae* UHCC 0037 | Nostocales | Yes | 85% of genes from heterocyst glycolipid BGC share at least 30% identity to this *pfa* operon |

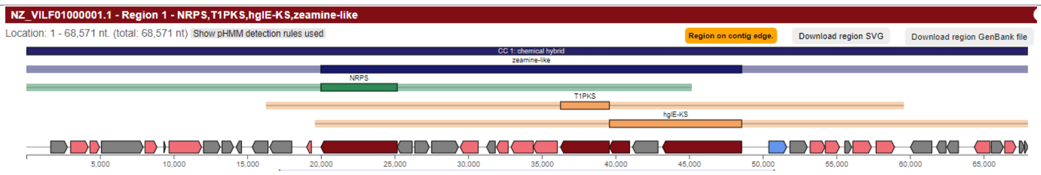

| | | | | |
|---|---|---|---|---|
| CP009962.1 | *Collimonas arenae* Cal35 | Burkholderiales | No | / |

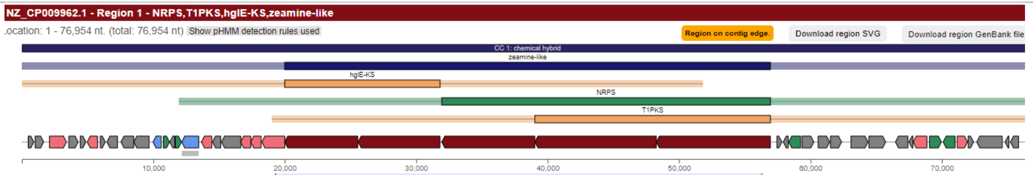

| | | | | |
|---|---|---|---|---|
| CP025429.1 | *Chromobacterium* sp. ATCC 53434 | Burkholderiales | No | / |

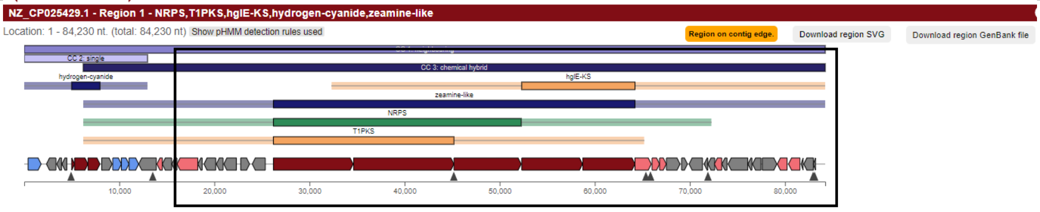

| | | | | |
|---|---|---|---|---|
| CP041743.1 | *Paraburkholderia megapolitana* LMG 23650 | Burkholderiales | No | / |

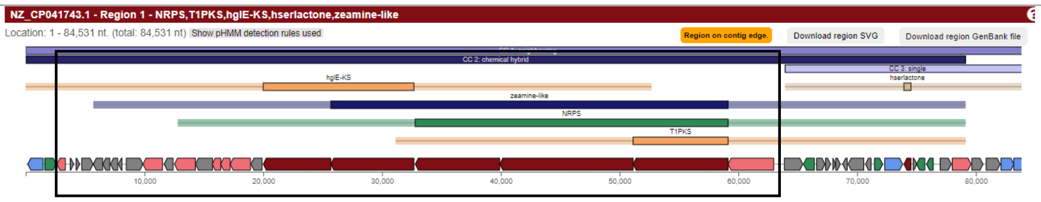

| | | | | |
|---|---|---|---|---|
| KE386747.1 | *Chitinimonas koreensis* DSM 17726 | Burkholderiales | No | / |

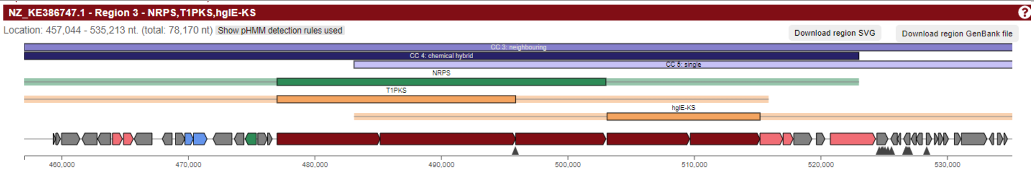

| | | | | |
|---|---|---|---|---|
| AUGX01000034.1 | *Ottowia thiooxydans* DSM 14619 | Burkholderiales | No | / |

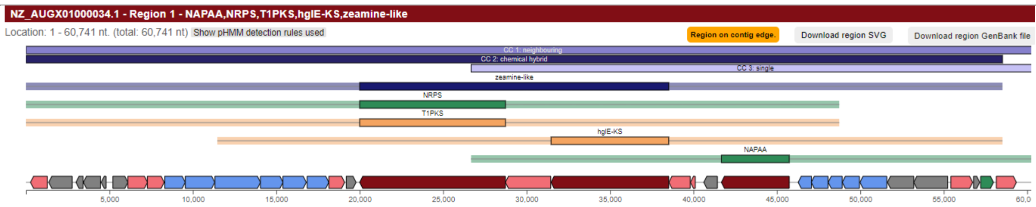

.

| | | | | |
|---|---|---|---|---|
| FWZX01000003.1 | *Tistlia consotensis* USBA 355 | Rhodospirillales | No | / |

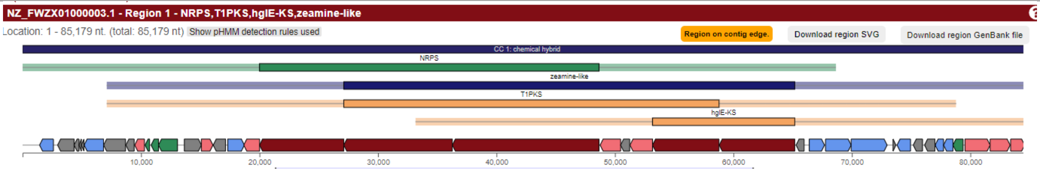

| | | | | |
|---|---|---|---|---|
| SNZH01000009.1 | *Tahibacter aquaticus* DSM 21667 | Lysobacterales | No | / |

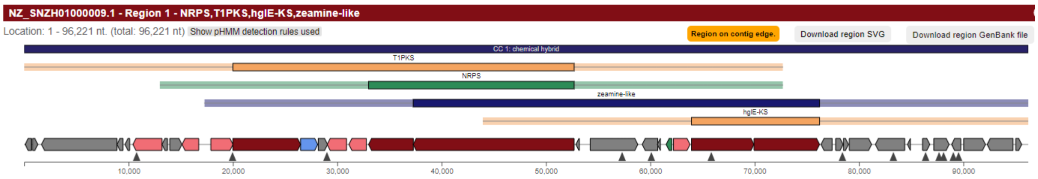

| | | | | |
|---|---|---|---|---|
| CP053590.1 | *Aquimarina* sp. TRL1 | Flavobacteriales | No | / |

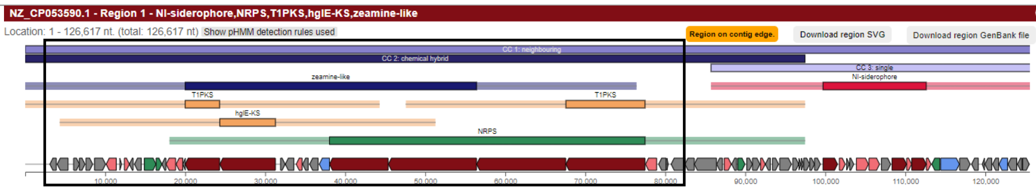

| | | | | |
|---|---|---|---|---|
| CP110012.1 | *Flavobacterium* sp. N502540 | Flavobacteriales | No | / |

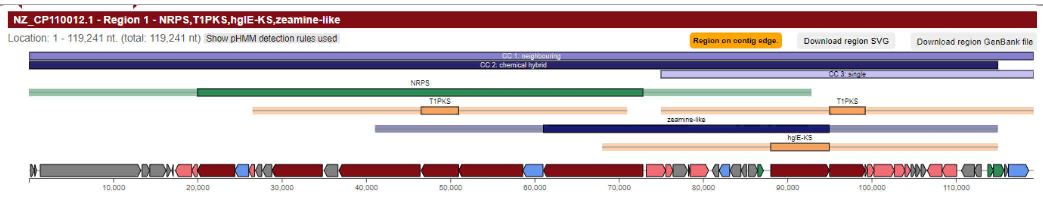

| | | | | |
|---|---|---|---|---|
| CP087134.1 | *Flavobacterium* sp. F-323 | Flavobacteriales | Yes | PfaBC homolog misannotated as T1PKS |

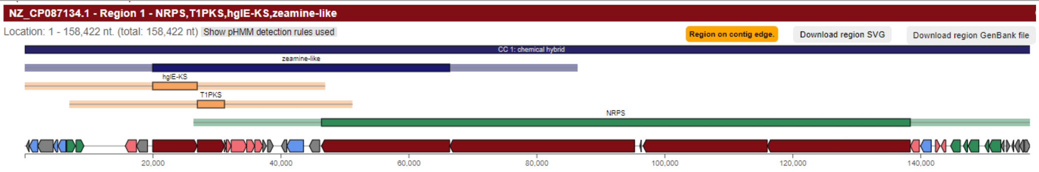

| | | | | |
|---|---|---|---|---|
| KB900627.1 | *Methylosinus* sp. LW4 | Hyphomicrobiales | Yes | 85% of genes from the phenolic lipids BGC share at least 30% identity to this *pfa* operon |

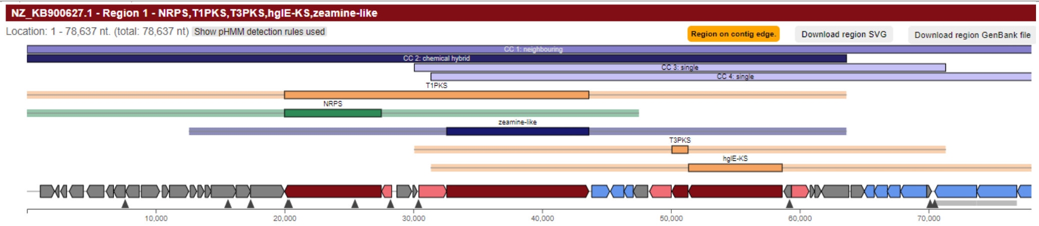

| | | | | |
|---|---|---|---|---|
| SOEJ01000010.1 | *Methylosinus* sp. sav-2 | Hyphomicrobiales | Yes | 85% of genes from the phenolic lipids BGC share at least 30% identity to this *pfa* operon |

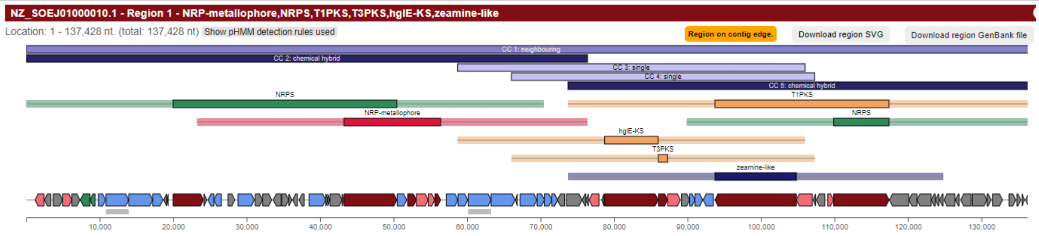

| | | | | |
|---|---|---|---|---|
| AP022333.1 | *Methylosinus* sp. C49 | Hyphomicrobiales | Yes | 85% of genes from the phenolic lipids BGC share at least 30% identity to this *pfa* operon |

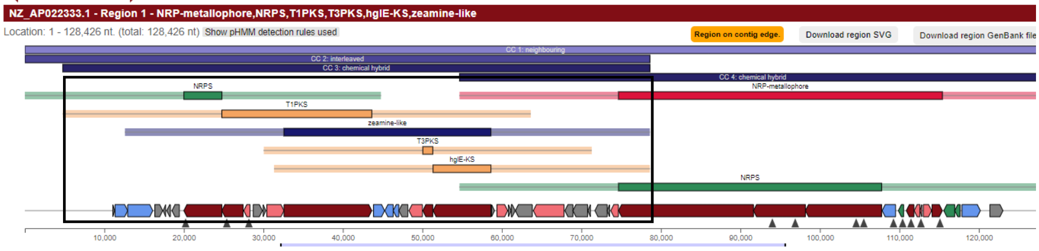

| | | | | |
|---|---|---|---|---|
| AZUO01000004.1 | *Methylosinus* sp. LW3 | Hyphomicrobiales | Yes | 85% of genes from the phenolic lipids BGC share at least 30% identity to this *pfa* operon |

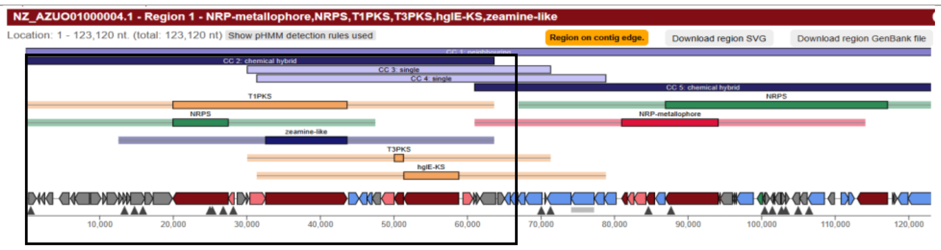

| | | | | |
|---|---|---|---|---|
| JACHHY010000001.1 | *Chitinivorax tropicus* DSM 27165 | Betaproteobacteria | No | / |

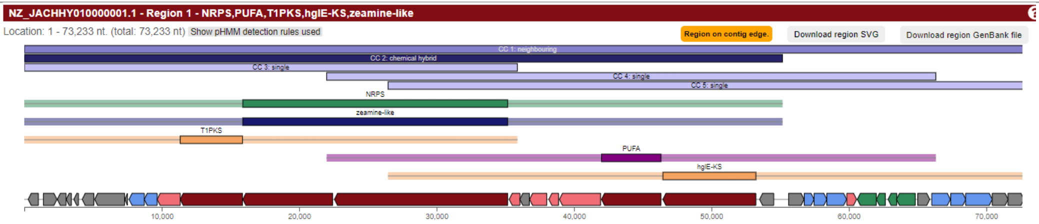

| | | | | |
|---|---|---|---|---|
| CP069161.1 | *Paludibacterium paludis* BCRC 80514 | Neisseriales | No | / |

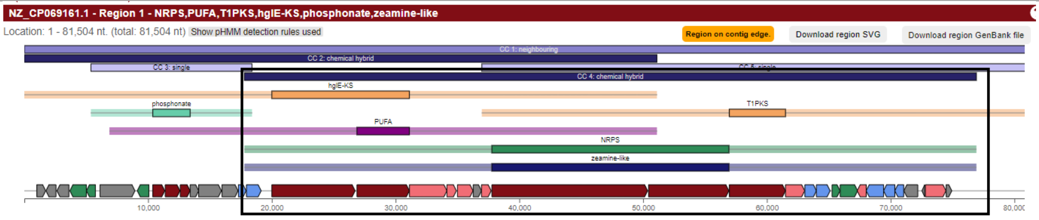

| | | | | |
|---|---|---|---|---|
| PEBU01000004.1 | *Bowmanella denitrificans* JL63 | Alteromonadales | No | / |

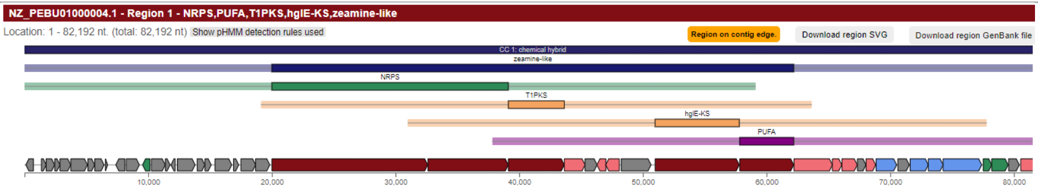

**Gram-positive bacteria**

| | | | | |
|---|---|---|---|---|
| CP001700.1 | *Catenulispora acidiphila* DSM 44928 | Catenulisporales | Yes | PfaBC homolog misannotated as T1PKS. The co-localization and synteny of the three types of biosynthetic genes are not conserved among closely-related strains |

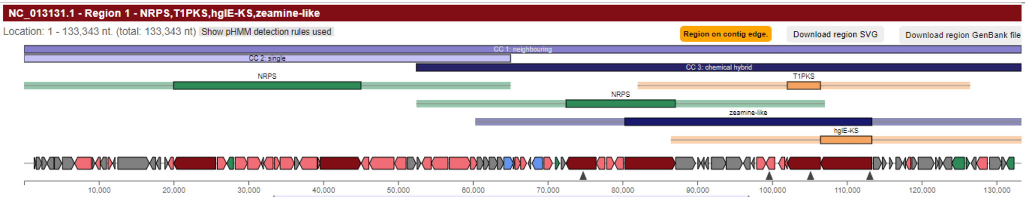

| | | | | |
|---|---|---|---|---|
| CP002047.1 | *Streptomyces bingchenggensis* BCW-1 | Kitasatosporales | Yes | The co-localization and synteny of the three types of biosynthetic genes are not conserved among closely-related strains. The *pfa*-like and the PKS-NRPS genes are separated by considerable distance with many intervening genes and no clear operon-like organization. |

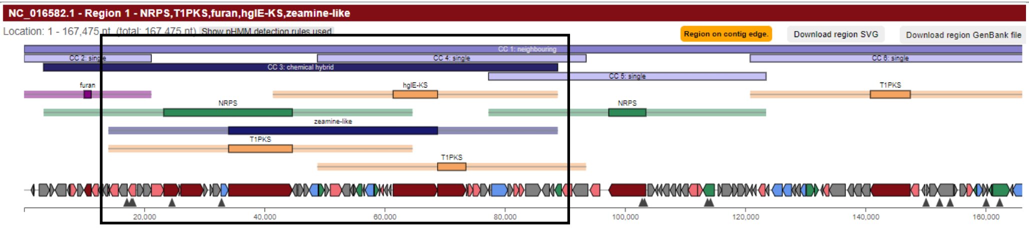

| | | | | |
|---|---|---|---|---|
| BMQK01000010.1 | *Streptomyces ruber* JCM 3131 | Kitasatosporales | Yes | PfaBC homolog misannotated as T1PKS. The co-localization and synteny of the three types of biosynthetic genes are not conserved among closely-related strains. |

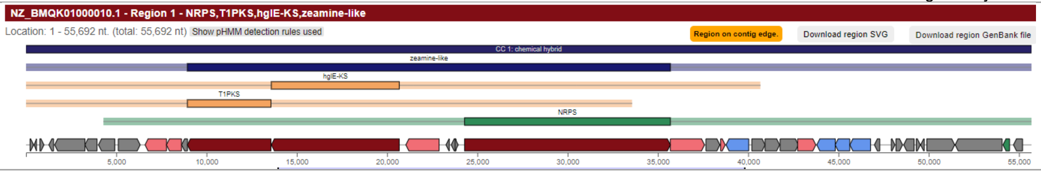

| | | | | |
|---|---|---|---|---|
| BNBJ01000010.1 | *Streptomyces griseus* JCM 4516 | Kitasatosporales | No | / |

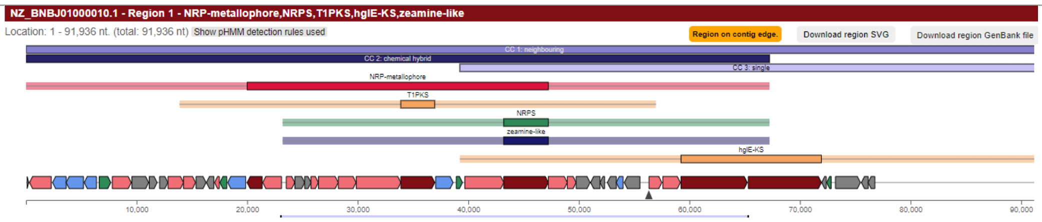

| | | | | |
|---|---|---|---|---|
| CP015849.1 | *Streptomyces* sp. SAT1 | Kitasatosporales | Yes | 100% of genes from the bacillibactin BGC share at least 30% identity to the NRPS operon |

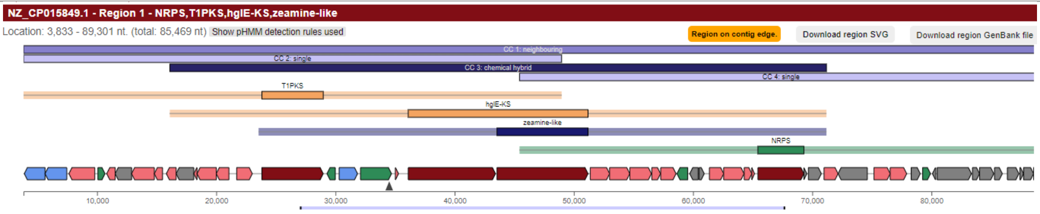

| | | | | |
|---|---|---|---|---|
| CP106840.1 | *Streptomyces* sp. Je 1-4 | Kitasatosporales | Yes | PfaBC homolog misannotated as T1PKS. The co-localization and synteny of the three types of biosynthetic genes are not conserved among closely-related strains. |

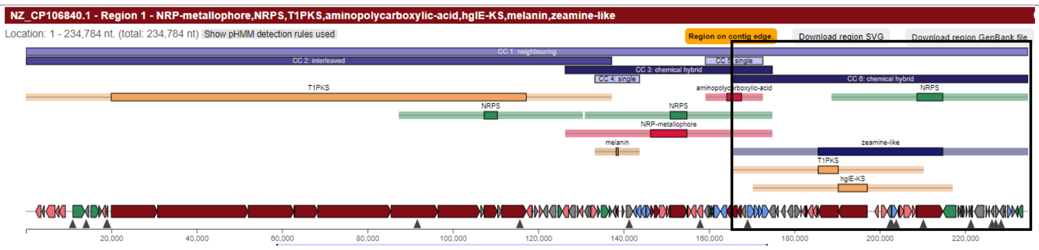

| | | | | |
|---|---|---|---|---|
| JAMFLF010000009.1 | *Streptomyces* sp. 43Y-GA-1 | Kitasatosporales | No | / |

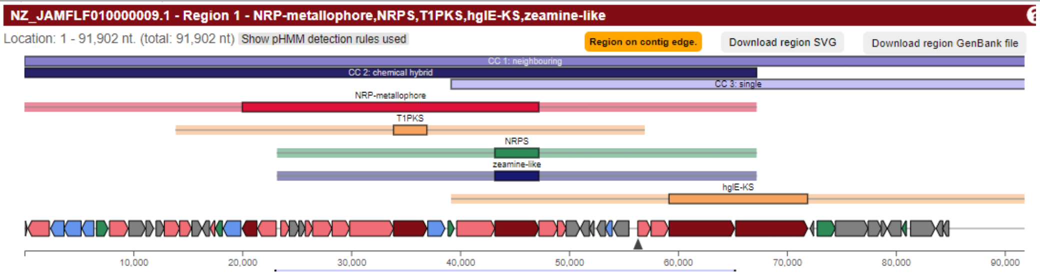

| | | | | |
|---|---|---|---|---|
| KB892157.1 | *Streptomyces* sp. PsTaAH-124 | Kitasatosporales | Yes | 100% of genes from the bacillibactin BGC share at least 30% identity to the NRPS operon. |

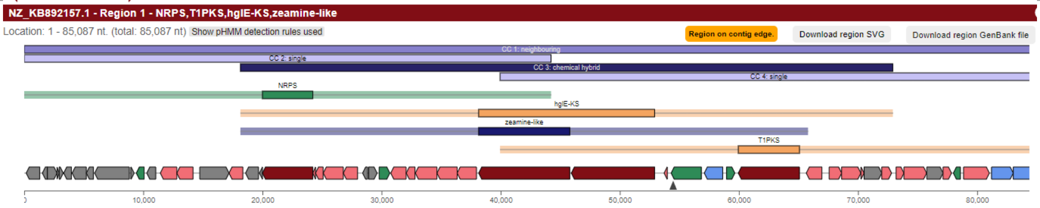

| | | | | |
|---|---|---|---|---|
| KL573545.1 | *Streptomyces baarnensis* NRRL B-2842 | Kitasatosporales | No | / |

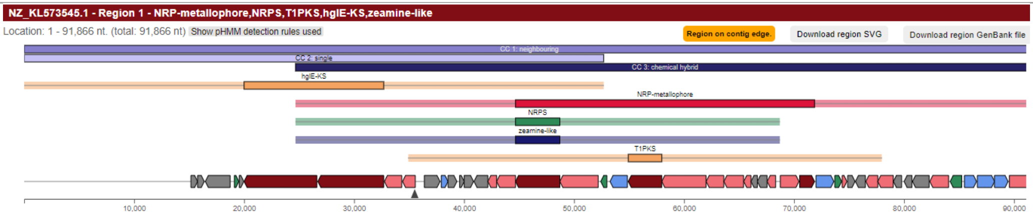

| | | | | |
|---|---|---|---|---|
| ML123034.1 | *Streptomyces* sp. ADI96-02 | Kitasatosporales | Yes | PfaBC homolog misannotated as T1PKS. The co-localization and synteny of the three types of biosynthetic genes are not conserved among closely-related strains. |

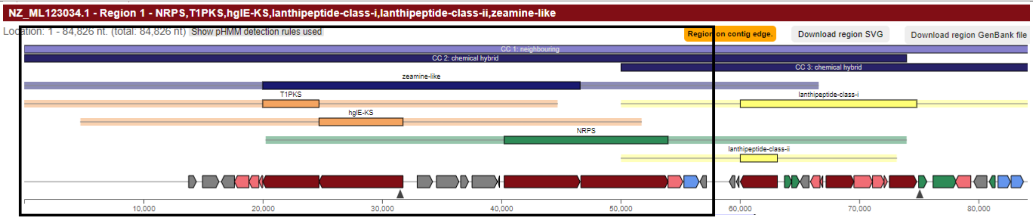

| | | | | |
|---|---|---|---|---|
| CP049782.1 | *Streptomyces* sp. ID38640 | Kitasatosporales | Yes | PfaBC homolog misannotated as T1PKS. The co-localization and synteny of the three types of biosynthetic genes are not conserved among closely-related strains. |

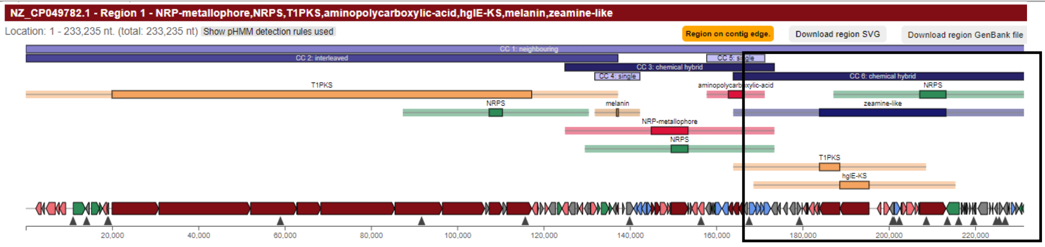

| | | | | |
|---|---|---|---|---|
| CP051486.1 | *Streptomyces pratensis* S10 | Kitasatosporales | Yes | PfaBC homolog misannotated as T1PKS. The co-localization and synteny of the three types of biosynthetic genes are not conserved among closely-related strains. |

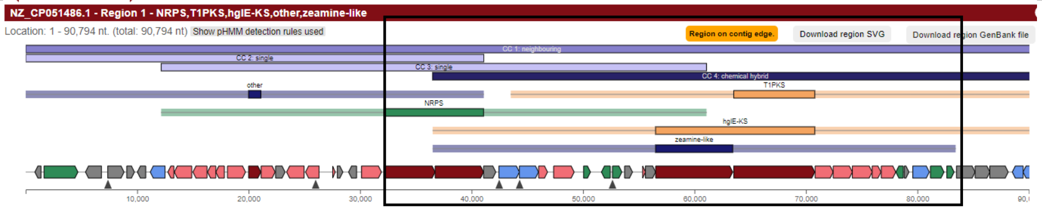

| | | | | |
|---|---|---|---|---|
| CP079114.1 | *Streptomyces* sp. WY228 | Kitasatosporales | No | / |

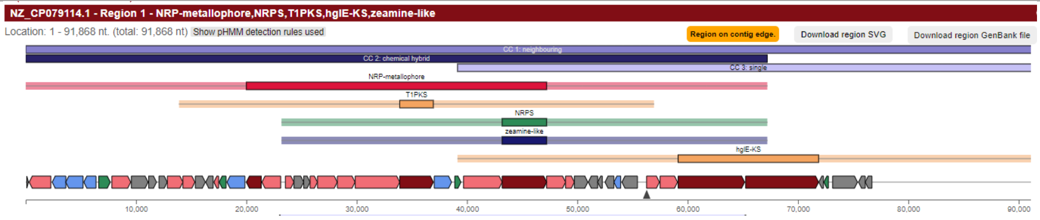

| | | | | |
|---|---|---|---|---|
| CP081496.1 | *Streptomyces* sp. BHT-5-2 | Kitasatosporales | Yes | The co-localization and synteny of the three types of biosynthetic genes are not conserved among closely-related strains. No clear operon-like organization. |

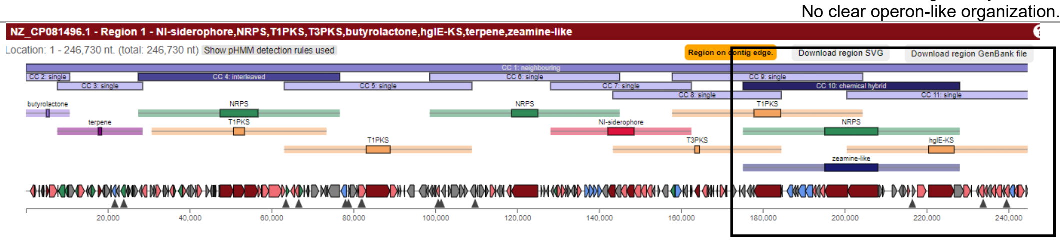

| | | | | |
|---|---|---|---|---|
| FNIX01000003.1 | *Lentzea jiangxiensis* CGMCC 4.6609 | Pseudonocardiales | Yes | The co-localization and synteny of the three types of biosynthetic genes are not conserved among closely-related strains. |

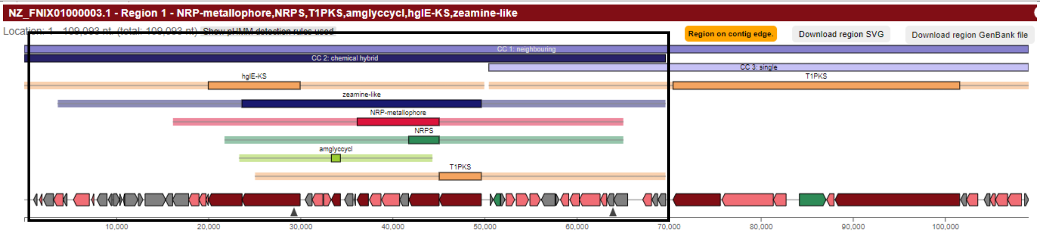

| | | | | |
|---|---|---|---|---|
| JAJCXE010000056.1 | *Lentzea* sp. CC55 | Pseudonocardiales | Yes | The co-localization and synteny of the three types of biosynthetic genes are not conserved among closely-related strains. |

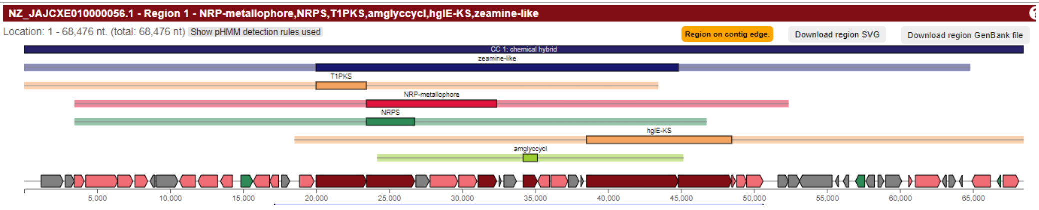

| | | | | |
|---|---|---|---|---|
| CP031142.1 | *Saccharopolyspora pogona* NRRL30141 | Pseudonocardiales | Yes | 82% of genes from the A83543A BGC share at least 30% identity to the T1PKS region |

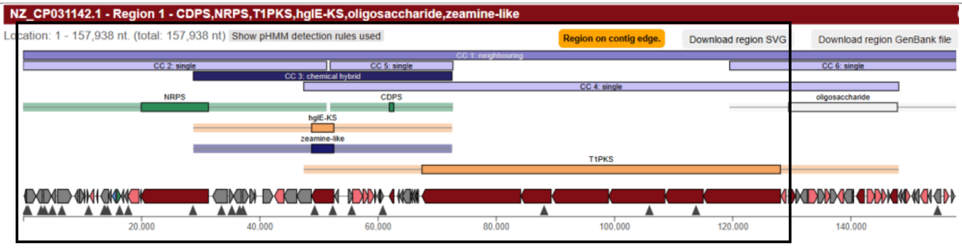

| | | | | |
|---|---|---|---|---|
| JADG01000010.1 | *Actinomadura oligospora* ATCC 43269 | Streptosporangiales | Yes | PfaBC homolog misannotated as T1PKS. |

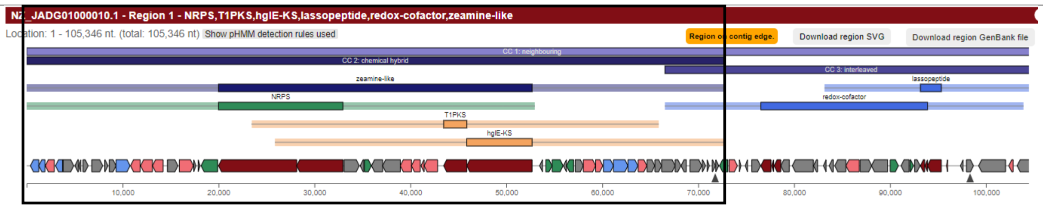

| | | | | |
|---|---|---|---|---|
| MPKW01000003.1 | *Mycolicibacterium* sp. CBMA 234 | Mycobacteriales | No | / |

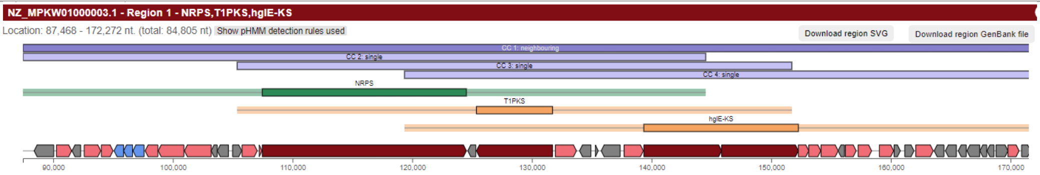

| | | | | |
|---|---|---|---|---|
| VOMB01000005.1 | *Mycobacterium fortunisiensis* TNTM28 | Mycobacteriales | No | / |

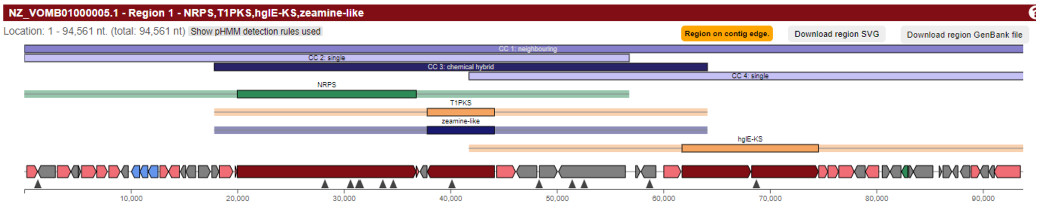

| | | | | |
|---|---|---|---|---|
| OCTY01000002.1 | *Mycobacterium simulans* FB-527 | Mycobacteriales | Yes | PfaBC homolog misannotated as T1PKS. |

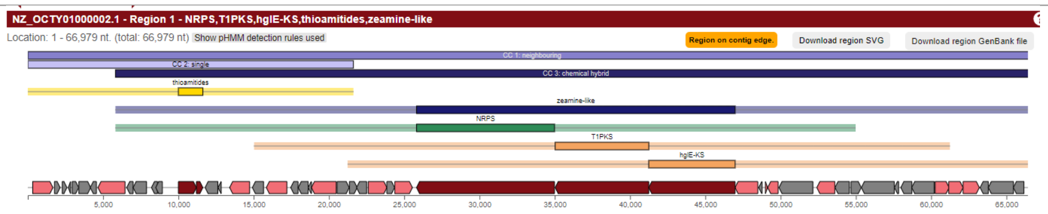
NZ_OCTY01000002.1 - Region 1 - NRPS,T1PKS,hglE-KS,thioamitides,zeamine-like
Location: 1 - 66,979 nt. (total: 66,979 nt)
Show pHMM detection rules used
Region on contig edge.
Download region SVG
Download region GenBank file
CC 1: neighbouring
CC 2: single
CC 3: chemical hybrid
thioamitides
zeamine-like
NRPS
T1PKS
hglE-KS
5,000
10,000
15,000
20,000
25,000
30,000
35,000
40,000
45,000
50,000
55,000
60,000
65,000

**Table S2. Predicted amino acid substrate specificities of the adenylation (A) domains in the zeamine/fabclavine BGCs from *Serratia plymuthica* RVH1, *Xenorhabdus hominickii* ANU1, *Bowmanella denitrificans* JL63, *Paludibacterium paludis* BCRC 80514 and *Chitinivorax tropicus* DSM 27165.** Overview of the predicted substrate binding residues in each A domain, along with specificity predictions obtained using antiSMASH and PARAS (Predictive Algorithm for Resolving Adenylation domain Selectivity).[20] The second amino acid incorporated (underlined) is distinct in zeamine and fabclavine compounds, and therefore, is used as a parameter to predict which out of the two metabolites *B. denitrificans, P. paludis* and *C. tropicus* are expected to produce.[40,41]

| | | |
|---|---|---|
| ***Serratia plymuthica* RVH1 (zeamines)** | | |
| **Zmn16_A1** | Predicted binding residues | DPRHLALLAK |
| | AntiSMASH prediction | Asp |
| | PARAS prediction | Asp (0.637) |
| | Actual amino acid incorporated | Asp |
| **Zmn16_A2** | Predicted binding residues | DTWTIASVSK |
| | AntiSMASH prediction | His |
| | PARAS prediction | His (0.674) |
| | Actual amino acid incorporated | His |
| **Zmn16_A3** | Predicted binding residues | DATKVGEVGK |
| | AntiSMASH prediction | Asn |
| | PARAS prediction | Asn (0.968) |
| | Actual amino acid incorporated | Asn |
| **Zmn16_A4** | Predicted binding residues | DATKVGEVGK |
| | AntiSMASH prediction | Asn |
| | PARAS prediction | Asn (0.960) |
| | Actual amino acids incorporated | Asn |
| **Zmn17_A5** | Predicted binding residues | DFWNIGMVHK |
| | AntiSMASH prediction | Thr |
| | PARAS prediction | Thr (0.939) |
| | Actual amino acids incorporated | Thr |
| **Zmn17_A6** | Predicted binding residues | DALFIGGTFK |
| | AntiSMASH prediction | Val |
| | PARAS prediction | Val (0.782) |
| | Actual amino acids incorporated | Val |
| ***Xenorhabdus hominickii* ANU1 (fabclavines)** | | |
| **Xbud_02640_A1** | Predicted binding residues | DPRHLSLLAK |
| | AntiSMASH prediction | X |
| | PARAS prediction | Asp (0.249) |
| | Actual amino acid incorporated | Asp |
| **Xbud_02640_A2** | Predicted binding residues | DTWTIASVGK |
| | AntiSMASH prediction | Phe |
| | PARAS prediction | Phe (0.636) |
| | Actual amino acid incorporated | Phe (Other variants might include His or Ala) |
| **Xbud_02640_A3** | Predicted binding residues | DATKVGEVGK |
| | AntiSMASH prediction | Asn |
| | PARAS prediction | Asn (0.813) |
| | Actual amino acid incorporated | |
| **Xbud_02640_A4** | Predicted binding residues | DATKVGEVGK |
| | AntiSMASH prediction | Asn |
| | PARAS prediction | Asn (0.965) |
| | Actual amino acids incorporated | |
| **Xbud_02641_A5** | Predicted binding residues | DFWNIGMVHK |
| | AntiSMASH prediction | Thr |
| | PARAS prediction | Thr (1.000) |
| | Actual amino acids incorporated | |
| **Xbud_02641_A6** | Predicted binding residues | DALFIGGTFK |
| | AntiSMASH prediction | Val |
| | PARAS prediction | Val (0.476) |
| | Actual amino acids incorporated | Val (Other variants might include Pro or Thr) |
| ***Bowmanella denitrificans* JL63** | | |
| **CSR04_RS09940_A1** | Predicted binding residues | DPRHLALLAK |
| | AntiSMASH prediction | Asp |
| | PARAS prediction | Asp (0.449) |

| | | |
|---|---|---|
| **CSR04_RS09940_A2** | Predicted binding residues | DTWTIASVSK |
| | AntiSMASH prediction | His |
| | PARAS prediction | His (0.417) |
| **CSR04_RS09940_A3** | Predicted binding residues | DATKVGEVGK |
| | AntiSMASH prediction | Asn |
| | PARAS prediction | Asn (0.672) |
| **CSR04_RS09940_A4** | Predicted binding residues | DATKVGEVGK |
| | AntiSMASH prediction | Asn |
| | PARAS prediction | Asn (0.870) |
| **CSR04_RS09945_A5** | Predicted binding residues | DFWNVGMVHK |
| | AntiSMASH prediction | Thr |
| | PARAS prediction | Thr (1.000) |
| **CSR04_RS09945_A6** | Predicted binding residues | DAFFLGATFK |
| | AntiSMASH prediction | X |
| | PARAS prediction | Val (0.374) |
| ***Paludibacterium paludis* BCRC 80514** | | |
| **JNO50_RS03090_A1** | Predicted binding residues | DPRHAALLAK |
| | AntiSMASH prediction | X |
| | PARAS prediction | Asp (0.247) |
| **JNO50_RS03090_A2** | Predicted binding residues | DSWTIASVSK |
| | AntiSMASH prediction | X |
| | PARAS prediction | His (0.379) |
| **JNO50_RS03090_A3** | Predicted binding residues | DATKVGEVGK |
| | AntiSMASH prediction | Asn |
| | PARAS prediction | Asn (0.941) |
| **JNO50_RS03090_A4** | Predicted binding residues | DATKVGEVGK |
| | AntiSMASH prediction | Asn |
| | PARAS prediction | Asn (0.854) |
| **JNO50_RS03095_A5** | Predicted binding residues | DFWNIGMVHK |
| | AntiSMASH prediction | Thr |
| | PARAS prediction | Thr (0.995) |
| **JNO50_RS03095_A6** | Predicted binding residues | DAMFIGGTFK |
| | AntiSMASH prediction | Val |
| | PARAS prediction | Val (0.577) |
| ***Chitinivorax tropicus* DSM 27165** | | |
| **HNQ59_RS00070_A1** | Predicted binding residues | DPRHAALLAK |
| | AntiSMASH prediction | X |
| | PARAS prediction | Asp (0.262) |
| **HNQ59_RS00070_A2** | Predicted binding residues | DTWTIASVSK |
| | AntiSMASH prediction | His |
| | PARAS prediction | His (0.448) |
| **HNQ59_RS00070_A3** | Predicted binding residues | DATKVGEVGK |
| | AntiSMASH prediction | Asn |
| | PARAS prediction | Asn (0.977) |
| **HNQ59_RS00070_A4** | Predicted binding residues | DATKVGEVGK |
| | AntiSMASH prediction | Asn |
| | PARAS prediction | Asn (0.854) |
| **HNQ59_RS00070_A5** | Predicted binding residues | DFWNVGMVHK |
| | AntiSMASH prediction | Thr |
| | PARAS prediction | Thr (1.000) |
| **HNQ59_RS00070_A6** | Predicted binding residues | DAFFFGGTFK |
| | AntiSMASH prediction | Val |
| | PARAS prediction | Val (0.533) |

**Table S3. Putative functions and closest homologs of the genes within and surrounding the chitinimine BGC.** Overview of genes, the putative function and predicted size of the proteins they encode, alongside the most similar known protein (protein ID, species and identity percentage), as determined by BlastP.

| Gene identifier | Putative function of gene product | Size (aa) | Top BlastP Hit | % Identity |
|---|---|---|---|---|
| *F559_RS0116810* | Type II toxin-antitoxin system VapB family antitoxin | 65 | MDO9386543.1 (*Thiobacillus* sp.) | 77.78 |
| *F559_RS0116815* | Type II toxin-antitoxin system VapC family toxin | 124 | WP_099407028.1 (*Chitinimonas* sp. BJB300) | 67.74 |
| *F559_RS26255* | Exonuclease domain-containing protein | 482 | GLR12994.1 (*Chitinimonas prasina*) | 66.95 |
| *F559_RS26260* | Lytic murein transglycosylase B | 396 | WP_331985645.1 (*Chitinimonas* sp.) | 73.80 |
| *F559_RS0116830* | Heat-inducible transcriptional repressor | 338 | WP_290334435.1 (*Chitinimonas viridis*) | 89.35 |
| *F559_RS0116835* | Transporter substrate-binding domain-containing protein | 259 | WP_290334434.1 (*Chitinimonas viridis*) | 79.10 |
| *F559_RS0116840* | Transporter substrate-binding domain-containing protein | 259 | WP_099405814.1 (*Chitinimonas sp. BJB300*) | 75.68 |
| *F559_RS26265* | SGNH/GDSL hydrolase family protein (carbohydrate modifying enzyme) | 471 | WP_335709779.1 (*Chitinimonas* sp. JJ19) | 89 |
| *F559_RS26270* | TOBE domain-containing protein (ATP-binding) | 259 | WP_331989049.1 (*Chitinimonas* sp.) | 67.57 |
| *F559_RS0116855* | Molybdate ABC transporter substrate-binding protein | 246 | WP_367788558.1 (Ch*itinivorax* sp. PXF-14) | 75.83 |
| *F559_RS0116860* | Molybdate ABC transporter permease subunit | 225 | WP_367788557.1 (Chitinivorax sp. PXF-14) | 77.52 |
| *F559_RS0116865* | Molybdenum ABC transporter ATP-binding protein | 357 | WP_331989043.1 (*Chitinimonas sp.*) | 69.86 |
| *F559_RS0116870* | AMP nucleosidase | 497 | WP_367788685.1 (*Chitinivorax* sp. PXF-14) | 82.49 |
| *F559_RS0116875* | ABC transporter substrate-binding protein | 254 | WP_035055253.1 (*Andreprevotia chitinilytica*) | 69.88 |
| *F559_RS0116880* | LysR family transcriptional regulator | 295 | WP_128683155.1 (*Pseudomonas aeruginosa*) | 70.34 |
| *F559_RS0116885* | CTP synthase | 251 | WP_199693560.1 (*Sorangium cellulosum*) | 66.38 |
| *F559_RS0116890* | Antibiotic biosynthesis monooxygenase | 121 | WP_263150353.1 (*Pseudomonas tohonis*) | 82.37 |
| *F559_RS26275 = chtnA* | Hybrid NRPS-type I PKS | 2715 | WP_099405553.1 (*Chitinimonas* sp. BJB300) | 65.99 |
| *F559_RS28020 = chtnB* | Hybrid NRPS-type I PKS | 3568 | WP_099405554.1 (*Chitinimonas* sp. BJB300) | 64.02 |
| *F559_RS0116905 = chtnC* | NRPS | 2379 | WP_099405555.1 (*Chitinimonas* sp. BJB300) | 62.48 |
| *F559_RS0116910 = chtnD* | Type I PKS | 2176 | WP_158228936.1 (*Chitinimonas* sp. BJB300) | 60.38 |
| *F559_RS0116915 = chtnE* | β-ketoacyl synthase N-terminal-like domain-containing protein | 1837 | WP_099405557.1 (*Chitinimonas* sp. BJB300) | 76.58 |
| *F559_RS26285 = chtnF* | PfaD family polyunsaturated fatty acid/polyketide biosynthesis protein | 602 | WP_099405558.1 (*Chitinimonas* sp. BJB300) | 72.67 |
| *F559_RS0116930 = chtnG* | Thioesterase II family protein | 265 | WP_325666666.1 (*Collimonas* sp.) | 63.95 |
| *F559_RS0116935 = chtnH* | Polysaccharide pyruvyl transferase family protein | 477 | WP_047243952.1 (*Chromobacterium subtsugae*) | 68.55 |
| *F559_RS26290* | Hypothetical protein | 220 | MBV8657729.1 (*Burkholderiales* bacterium) | 62.11 |
| *F559_RS26295* | EAL domain-containing protein, partial | 1190 | WP_331987312.1 (*Chitinimonas* sp.) | 72.51 |
| *F559_RS29640* | Hypothetical protein | 291 | WP_296888885.1 (*Thiobacillus* sp.) | 58.94 |
| *F559_RS0116950* | HEPN domain-containing protein | 172 | HST20945.1 (*Blastocatellia bacterium)* | 45.78 |
| *F559_RS0116955* | Nucleotidyltransferase domain-containing protein | 93 | MCB2263725.1 (Candidatus Thiosymbion ectosymbiont of *Robbea hypermnestra*) | 53.76 |
| *F559_RS29645* | Hypothetical protein | 170 | WP_084091439.1 (*Andreprevotia lacus*) | 59.17 |
| *F559_RS0116960* | Hypothetical protein | 134 | - | - |
| *F559_RS26300* | GNAT family N-acetyltransferase | 160 | OWQ46416.1 (*Roseateles noduli*) | 66.67 |
| *F559_RS0116970* | Hypothetical protein | 145 | WP_285234984.1 (*Paucibacter sediminis*) | 67.20 |
| *F559_RS29650* | AbrB/MazE/SpoVT family DNA-binding domain-containing protein | 91 | WP_251972849.1 (*Sphaerotilus microaerophilus*) | 39.78 |
| *F559_RS0116980* | Type II toxin-antitoxin system VapC family toxin | 128 | MDX2043469.1 (*Acidobacteriota* bacterium) | 46.51 |
| *F559_RS0116985* | VOC (vicinal oxygen chelate) family protein Glyoxalase | 125 | HTE15913.1 (*Burkholderiales* bacterium) | 74.40 |
| *F559_RS0116990* | DUF421 domain-containing protein | 171 | HEX5128416.1 (*Usitatibacter* sp.) | 57.89 |
| *F559_RS0116995* | AsmA family protein | 671 | WP_378161007.1 (*Chitinimonas lacunae*) | 54.02 |
| *F559_RS30135* | Hypothetical protein | 111 | - | - |
| *F559_RS30140* | Glycine zipper 2TM domain-containing protein | 145 | WP_300111117.1 (*Rhodoferax* sp.) | 40.58 |
| *F559_RS26310* | Hypothetical protein | 91 | WP_378161013.1 (*Chitinimonas lacunae*) | 62.34 |

**Table S4**. Overview of NMR signals observed for chitinimine I isolated from *C. koreensis* DSM 17726 (DMSO-d6, $^{1}H$ 600 MHz, $^{13}C$ 151 MHz). nd = not determinable.

| Position | $^{1}H$ (ppm) | $^{13}C$ (ppm) |
|---|---|---|
| **Ile** | | |
| C=O | - | 170.5 |
| α | 4.01 | 57.9 |
| β | 1.78 | 35.0 |
| β-$CH_3$ | 0.86 | 14.8 |
| γ | 1.32,1.16 | 25.6 |
| δ | 0.82 | 11.4 |
| NH | 8.11 | - |
| **4-amino-3-hydroxypentanoic acid** | | |
| 1” | - | 171.8 |
| 2” | 2.50, 2.08 | 39.9 |
| 3” | 3.92 | 69.6 |
| 3”-OH | 4.83 | - |
| 4” | 3.41 | 49.4 |
| 4”-NH | 7.75 | - |
| 5” | 1.01 | 14.8 |
| **Gln** | | |
| C=O | - | 171.1 |
| α | 4.03 | 54.6 |
| β | 1.84 | 27.6 |
| γ | 2.15, 1.95 | 32.0 |
| δ | - | 173.7 |
| δ-$NH_2$ | 7.67, 6.63 | - |
| NH | 8.58 | - |
| **Leu** | | |
| C=O | - | 172.2 |
| α | 4.01 | 52.8 |
| β | 1.59, 1.47 | 39.4 |
| γ | 1.73 | 24.2 |
| γ-$CH_3$ | 0.80 | 20.4 |
| γ-$CH_3$ | 0.90 | 23.0 |
| NH | 8.89 | - |
| **Specialized lipid** | | |
| 1 | - | 172.9 |
| 2 | 2.50, 2.42 | 36.4 |
| 3 | 3.80 | 45.7 |
| 3-NH | 8.17 | - |
| 4 | 1.44 | 32.7 |
| 5 | 1.92 | 29.0 |
| 6 | 5.36 | 129.7-129.5 |
| 7 | 5.36 | 129.7-129.5 |
| 8 | 2.01 | 31.8-31.2 |
| 9 | 2.01 | 31.8-31.2 |
| 10 | 5.36 | 129.7-129.5 |
| 11 | 5.36 | 129.7-129.5 |
| 12 | 2.01 | 31.8-31.2 |
| 13 | 2.01 | 31.8-31.2 |
| 14 | 5.36 | 129.7-129.5 |
| 15 | 5.36 | 129.7-129.5 |
| 16 | 2.01 | 31.8-31.2 |
| 17 | 2.01 | 31.8-31.2 |
| 18 | 5.39 | 126.1 |
| 19 | 5.42 | 131.6 |
| 20 | 2.17, 1.98 | 39.1 |
| 21 | 3.82 | 68.2 |
| 21-OH | nd | - |
| 22 | 1.46, 1.00 | 37.9 |
| 23 | 3.95 | 64.4 |

| | | |
|---|---|---|
| 24 | 1.06 | 22.0 |
| **Pyruvate** | | |
| 1’ | - | 172.8 |
| 2’ | - | 97.8 |
| 3’ | 1.44 | 17.4 |
| 2’-OH | nd | - |
| 2’-OH | nd | - |

**Table S5**. Overview of NMR signals observed for chitinimine II isolated from *C. koreensis* DSM 17726 (DMSO-d6, $^{1}H$ 600 MHz, $^{13}C$ 151 MHz).

| Position | $^{1}H$ (ppm) | $^{13}C$ (ppm) |
|---|---|---|
| **Val** | | |
| C=O | - | nd |
| $\alpha$ | 3.74 | 60.5 |
| $\beta$ | 1.91 | 28.7 |
| $\gamma$ | 0.90, 0.88 | 18.6, 19.3 |
| NH | 8.21 | - |
| **4-amino-3-hydroxypentanoic acid** | | |
| 1” | - | nd |
| 2” | 2.50, 2.03 | 40.0 |
| 3” | 3.92 | 69.6 |
| 3”-OH | 4.03 | - |
| 4” | 3.41 | 49.4 |
| 4”-NH | 7.93 | - |
| 5” | 1.01 | 14.8 |
| **Gln** | | |
| C=O | - | nd |
| $\alpha$ | 4.03 | 54.6 |
| $\beta$ | 1.84 | 28.0 |
| $\gamma$ | 2.20, 1.92 | 32.2 |
| $\delta$ | - | nd |
| $\delta$-$NH_2$ | 7.81, 6.60 | - |
| NH | 8.88 | - |
| **Leu-2** | | |
| C=O | - | nd |
| $\alpha$ | 4.03 | 52.9 |
| $\beta$ | 1.59, 1.47 | 39.4 |
| $\gamma$ | 1.73 | 24.2 |
| $\gamma$-$CH_3$ | 0.80 | 20.4 |
| $\gamma$-$CH_3$ | 0.90 | 23.0 |
| NH | 9.20 | - |
| **Specialized lipid** | | |
| 1 | - | nd |
| 2 | 2.52, 2.44 | 36.2 |
| 3 | 3.78 | 45.5 |
| 3-NH | 8.09 | - |
| 4 | 1.44 | 32.7 |
| 5 | 1.92 | 29.0 |
| 6 | 5.36 | 129.7-129.5 |
| 7 | 5.36 | 129.7-129.5 |
| 8 | 2.01 | 31.8-31.2 |
| 9 | 2.01 | 31.8-31.2 |
| 10 | 5.36 | 129.7-129.5 |
| 11 | 5.36 | 129.7-129.5 |
| 12 | 2.01 | 31.8-31.2 |
| 13 | 2.01 | 31.8-31.2 |
| 14 | 5.36 | 129.7-129.5 |
| 15 | 5.36 | 129.7-129.5 |

| | | |
|---|---|---|
| 16 | 2.01 | 31.8-31.2 |
| 17 | 2.01 | 31.8-31.2 |
| 18 | 5.39 | 126.1 |
| 19 | 5.42 | 131.6 |
| 20 | 2.17, 1.98 | 39.1 |
| 21 | 3.82 | 68.2 |
| 21-OH | nd | - |
| 22 | 1.46, 1.00 | 37.9 |
| 23 | 3.95 | 64.4 |
| 24 | 1.06 | 22.0 |
| **Pyruvate** | | |
| 1’ | - | nd |
| 2’ | - | 97.8 |
| 3’ | 1.44 | 17.4 |
| 2’-OH | nd | - |
| 2’-OH | nd | - |

**Table S6**. Overview of NMR signals observed for chitinimine III isolated from *C. koreensis* DSM 17726 (DMSO-d6, $^{1}H$ 600 MHz, $^{13}C$ 151 MHz).

| Position | $^{1}H$ (ppm) | $^{13}C$ (ppm) |
|---|---|---|
| **Leu-1** | | |
| C=O | - | nd |
| $\alpha$ | 3.96 | 53.2 |
| $\beta$ | 1.44, 1.39 | 39.8 |
| $\gamma$ | 1.66 | 24.2 |
| $\gamma$-$CH_3$ | 0.82 | 21.2 |
| $\gamma$-$CH_3$ | 0.88 | 22.8 |
| NH | 8.34 | - |
| **4-amino-3-hydroxypentanoic acid** | | |
| 1’’ | - | 171.5 |
| 2’’ | 2.45, 2.00 | 40.0 |
| 3’’ | 3.90 | 69.6 |
| 3”-OH | 4.82 | - |
| 4’’ | 3.41 | 49.4 |
| 4”-NH | 7.74 | - |
| 5’’ | 1.01 | 14.8 |
| **Gln** | | |
| C=O | - | 171.1 |
| $\alpha$ | 4.03 | 54.6 |
| $\beta$ | 1.84 | 27.6 |
| $\gamma$ | 2.15, 1.95 | 32.0 |
| $\delta$ | - | 173.7 |
| $\delta$-$NH_2$ | 7.67, 6.63 | - |
| NH | 8.58 | - |
| **Leu-2** | | |
| C=O | - | 172.2 |
| $\alpha$ | 4.01 | 52.7 |
| $\beta$ | 1.59, 1.47 | 39.4 |
| $\gamma$ | 1.73 | 24.2 |
| $\gamma$-$CH_3$ | 0.80 | 20.4 |
| $\gamma$-$CH_3$ | 0.90 | 23.0 |
| NH | 8.89 | - |
| **Specialized lipid** | | |
| 1 | - | 172.9 |
| 2 | 2.50, 2.40 | 36.6 |
| 3 | 3.79 | 45.5 |

| | | |
|---|---|---|
| 3-NH | 8.16 | - |
| 4 | 1.44 | 32.8 |
| 5 | 1.92 | 29.0 |
| 6 | 5.36 | 129.7-129.5 |
| 7 | 5.36 | 129.7-129.5 |
| 8 | 2.01 | 31.8-31.2 |
| 9 | 2.01 | 31.8-31.2 |
| 10 | 5.36 | 129.7-129.5 |
| 11 | 5.36 | 129.7-129.5 |
| 12 | 2.01 | 31.8-31.2 |
| 13 | 2.01 | 31.8-31.2 |
| 14 | 5.36 | 129.7-129.5 |
| 15 | 5.36 | 129.7-129.5 |
| 16 | 2.01 | 31.8-31.2 |
| 17 | 2.01 | 31.8-31.2 |
| 18 | 5.39 | 126.1 |
| 19 | 5.42 | 131.6 |
| 20 | 2.17, 1.98 | 39.1 |
| 21 | 3.82 | 68.2 |
| 21-OH | nd | - |
| 22 | 1.46, 1.00 | 37.9 |
| 23 | 3.95 | 64.4 |
| 24 | 1.06 | 22.0 |
| **Pyruvate** | | |
| 1’ | - | 172.8 (b) |
| 2’ | - | 97.8 |
| 3’ | 1.44 | 17.4 |
| 2’-OH | nd | - |
| 2’-OH | nd | - |

**Table S7. Predicted amino acid substrate specificities of the adenylation (A) domains in the chitinimine NRPS modules.** Overview of the predicted substrate binding residues in each A domain, along with specificity predictions obtained using antiSMASH and PARAS.[20] Where multiple predictions were given, the actual amino acid incorporated into the chitinimines is highlighted in bold.

| | | |
|---|---|---|
| **ChtnA_A1** | Predicted binding residues | DIWQFGLILK |
| | AntiSMASH prediction | X |
| | PARAS prediction | 1. Ala (0.131) **2. Leu (0.130)** |
| | Actual amino acid incorporated | Leu |
| **ChtnB_A2** | Predicted binding residues | DAQDLGVVDK |
| | AntiSMASH prediction | Gln |
| | PARAS prediction | **1. Gln (0.494)** |
| | Actual amino acid incorporated | Gln |
| **ChtnB_A3** | Predicted binding residues | DVWHFSLIEK |
| | AntiSMASH prediction | Ser |
| | PARAS prediction | **1. Ala (0.478)** |
| | Actual amino acid incorporated | Ala |
| **ChtnC_A4** | Predicted binding residues | DALFMGVVLK |
| | AntiSMASH prediction | Ile |
| | PARAS prediction | **1. Leu (0.338) 2. Valine (0.183) 3. Isoleucine (0.156)** |
| | Actual amino acids incorporated | Leu, Val and Ile |

**Table S8. Condensation (C) domains used for phylogenetic analysis of the C domains within ChtnA-C.** Compilation of 172 C domains from the NaPDoS database, including the name and class of each C domain, their associated protein, MiBiG and PubMed IDs, the metabolic product and type of their cognate BGC, species name and strain, and gene product annotation.[21] na = not annotated.

| BGC product | BGC type | Domain name | Domain class | Genbank protein ID | MIBiG ID | PubMed ID | Species name | Strain | Gene product name |
|---|---|---|---|---|---|---|---|---|---|
| actinomycin | NRPS | actinomycin_C04_DCL | DCL | ADG27359 | BGC0000296 | 20304989 | *Streptomyces anulatus* | ATCC 11523 | *AcmC* |
| actinomycin | NRPS | actinomycin_C03_epimerization | epimerization | ADG27358 | BGC0000296 | 20304989 | *Streptomyces anulatus* | ATCC 11523 | *AcmB* |
| actinomycin | NRPS | actinomycin_C02_LCL | LCL | ADG27358 | BGC0000296 | 20304989 | *Streptomyces anulatus* | ATCC 11523 | *AcmB* |
| actinomycin | NRPS | actinomycin_C05_LCL | LCL | ADG27359 | BGC0000296 | 20304989 | *Streptomyces anulatus* | ATCC 11523 | *AcmC* |
| actinomycin | NRPS | actinomycin_C06_LCL | LCL | ADG27359 | BGC0000296 | 20304989 | *Streptomyces anulatus* | ATCC 11523 | *AcmC* |
| actinomycin | NRPS | actinomycin_C01_starter | starter | ADG27358 | BGC0000296 | 20304989 | *Streptomyces anulatus* | ATCC 11523 | *AcmB* |
| anabaenopeptilide | NRPS | anabaenopeptilide_C01_LCL | LCL | CAC01603 | na | 10931313 | *Anabaena sp.* | str 90 | *AdpA* |
| anabaenopeptilide | NRPS | anabaenopeptilide_C02_LCL | LCL | CAC01604 | na | 10931313 | *Anabaena sp.* | str 90 | *AdpB* |
| anabaenopeptilide | NRPS | anabaenopeptilide_C03_LCL | LCL | CAC01604 | na | 10931313 | *Anabaena sp.* | str 90 | *AdpB* |
| anabaenopeptilide | NRPS | anabaenopeptilide_C04_LCL | LCL | CAC01604 | na | 10931313 | *Anabaena sp.* | str 90 | *AdpB* |
| anabaenopeptilide | NRPS | anabaenopeptilide_C05_LCL | LCL | CAC01604 | na | 10931313 | *Anabaena sp.* | str 90 | *AdpB* |
| anabaenopeptilide | NRPS | anabaenopeptilide_C06_LCL | LCL | CAC01606 | na | 10931313 | *Anabaena sp.* | str 90 | *AdpD* |
| arthrofactin | NRPS | arthrofactin_C02_DCL | DCL | BAC67534 | BGC0000305 | 14522057 | *Pseudomonas sp.* | MIS38 | *ArfA* |
| arthrofactin | NRPS | arthrofactin_C03_DCL | DCL | BAC67535 | BGC0000305 | 14522057 | *Pseudomonas sp.* | MIS38 | *ArfB* |
| arthrofactin | NRPS | arthrofactin_C04_DCL | DCL | BAC67535 | BGC0000305 | 14522057 | *Pseudomonas sp.* | MIS38 | *ArfB* |
| arthrofactin | NRPS | arthrofactin_C05_DCL | DCL | BAC67535 | BGC0000305 | 14522057 | *Pseudomonas sp.* | MIS38 | *ArfB* |
| arthrofactin | NRPS | arthrofactin_C06_DCL | DCL | BAC67535 | BGC0000305 | 14522057 | *Pseudomonas sp.* | MIS38 | *ArfB* |
| arthrofactin | NRPS | arthrofactin_C07_DCL | DCL | BAC67536 | BGC0000305 | 14522057 | *Pseudomonas sp.* | MIS38 | *ArfC* |
| arthrofactin | NRPS | arthrofactin_C09_DCL | DCL | BAC67536 | BGC0000305 | 14522057 | *Pseudomonas sp.* | MIS38 | *ArfC* |
| arthrofactin | NRPS | arthrofactin_C08_LCL | LCL | BAC67536 | BGC0000305 | 14522057 | *Pseudomonas sp.* | MIS38 | *ArfC* |
| arthrofactin | NRPS | arthrofactin_C10_LCL | LCL | BAC67536 | BGC0000305 | 14522057 | *Pseudomonas sp.* | MIS38 | *ArfC* |
| arthrofactin | NRPS | arthrofactin_C11_LCL | LCL | BAC67536 | BGC0000305 | 14522057 | *Pseudomonas sp.* | MIS38 | *ArfC* |
| arthrofactin | NRPS | arthrofactin_C01_starter | starter | BAC67534 | BGC0000305 | 14522057 | *Pseudomonas sp.* | MIS38 | *ArfA* |
| bacillibactin | NRPS | bacillibactin_C02_LCL | LCL | CAB15186 | BGC0000309 | 11112781 | *Bacillus subtilis subsp. subtilis* | str 168 | *DhbF* |
| bacillibactin | NRPS | bacillibactin_C01_starter | starter | CAB15186 | BGC0000309 | 11112781 | *Bacillus subtilis subsp. subtilis* | str 168 | *DhbF* |
| bacitracin | NRPS | bacitracin_C01_cyclization | cyclization | AAC06346 | BGC0000310 | 9427658 | *Bacillus licheniformis* | ATCC 10716 | *BacA* |
| bacitracin | NRPS | bacitracin_C05_DCL | DCL | AAC06346 | BGC0000310 | 9427658 | *Bacillus licheniformis* | ATCC 10716 | *BacA* |
| bacitracin | NRPS | bacitracin_C09_DCL | DCL | AAC06348 | BGC0000310 | 9427658 | *Bacillus licheniformis* | ATCC 10716 | *BacC* |
| bacitracin | NRPS | bacitracin_C12_DCL | DCL | AAC06348 | BGC0000310 | 9427658 | *Bacillus licheniformis* | ATCC 10716 | *BacC* |
| bacitracin | NRPS | bacitracin_C15_DCL | DCL | AAC06348 | BGC0000310 | 9427658 | *Bacillus licheniformis* | ATCC 10716 | *BacC* |
| bacitracin | NRPS | bacitracin_C04_epimerization | epimerization | AAC06346 | BGC0000310 | 9427658 | *Bacillus licheniformis* | ATCC 10716 | *BacA* |
| bacitracin | NRPS | bacitracin_C08_epimerization | epimerization | AAC06347 | BGC0000310 | 9427658 | *Bacillus licheniformis* | ATCC 10716 | *BacB* |
| bacitracin | NRPS | bacitracin_C11_epimerization | epimerization | AAC06348 | BGC0000310 | 9427658 | *Bacillus licheniformis* | ATCC 10716 | *BacC* |
| bacitracin | NRPS | bacitracin_C14_epimerization | epimerization | AAC06348 | BGC0000310 | 9427658 | *Bacillus licheniformis* | ATCC 10716 | *BacC* |
| bacitracin | NRPS | bacitracin_C02_LCL | LCL | AAC06346 | BGC0000310 | 9427658 | *Bacillus licheniformis* | ATCC 10716 | *BacA* |
| bacitracin | NRPS | bacitracin_C03_LCL | LCL | AAC06346 | BGC0000310 | 9427658 | *Bacillus licheniformis* | ATCC 10716 | *BacA* |
| bacitracin | NRPS | bacitracin_C06_LCL | LCL | AAC06347 | BGC0000310 | 9427658 | *Bacillus licheniformis* | ATCC 10716 | *BacB* |
| bacitracin | NRPS | bacitracin_C07_LCL | LCL | AAC06347 | BGC0000310 | 9427658 | *Bacillus licheniformis* | ATCC 10716 | *BacB* |
| bacitracin | NRPS | bacitracin_C10_LCL | LCL | AAC06348 | BGC0000310 | 9427658 | *Bacillus licheniformis* | ATCC 10716 | BacC |
| balhimycin | NRPS | balhimycin_C01_DCL | DCL | CAC48360 | BGC0000311 | 11932455 | *Amycolatopsis balhimycina* | DSM 5908 | BpsA |
| balhimycin | NRPS | balhimycin_C03_DCL | DCL | CAC48360 | BGC0000311 | 11932455 | *Amycolatopsis balhimycina* | DSM 5908 | BpsA |
| balhimycin | NRPS | balhimycin_C04_DCL | DCL | CAC48361 | BGC0000311 | 11932455 | *Amycolatopsis balhimycina* | DSM 5908 | BpsB |

| | | | | | | | | | |
|---|---|---|---|---|---|---|---|---|---|
| balhimycin | NRPS | balhimycin_C06_DCL | DCL | CAC48361 | BGC0000311 | 11932455 | *Amycolatopsis balhimycina* | DSM 5908 | BpsB |
| balhimycin | NRPS | balhimycin_C08_DCL | DCL | CAC48361 | BGC0000311 | 11932455 | *Amycolatopsis balhimycina* | DSM 5908 | BpsB |
| balhimycin | NRPS | balhimycin_C09_DCL | DCL | CAC48362 | BGC0000311 | 11932455 | *Amycolatopsis balhimycina* | DSM 5908 | BpsC |
| balhimycin | NRPS | balhimycin_C02_epimerization | epimerization | CAC48360 | BGC0000311 | 11932455 | *Amycolatopsis balhimycina* | DSM 5908 | BpsA |
| balhimycin | NRPS | balhimycin_C05_epimerization | epimerization | CAC48361 | BGC0000311 | 11932455 | *Amycolatopsis balhimycina* | DSM 5908 | BpsB |
| balhimycin | NRPS | balhimycin_C07_epimerization | epimerization | CAC48361 | BGC0000311 | 11932455 | *Amycolatopsis balhimycina* | DSM 5908 | BpsB |
| balhimycin | NRPS | balhimycin_C10_LCL | LCL | CAC48362 | BGC0000311 | 11932455 | *Amycolatopsis balhimycina* | DSM 5908 | BpsC |
| bleomycin | PKS-NRPS | bleomycin_C01_condensation | condensation | AAG02355 | BGC0000963 | 11048953 | *Streptomyces verticillus* | ATCC 15003 | BlmX |
| bleomycin | PKS-NRPS | bleomycin_C03_condensation | condensation | AAG02356 | BGC0000963 | 11048953 | *Streptomyces verticillus* | ATCC 15003 | BlmIX |
| bleomycin | PKS-NRPS | bleomycin_C04_condensation | condensation | AAG02358 | BGC0000963 | 11048953 | *Streptomyces verticillus* | ATCC 15003 | BlmVII |
| bleomycin | PKS-NRPS | bleomycin_C05_condensation | condensation | AAG02359 | BGC0000963 | 11048953 | *Streptomyces verticillus* | ATCC 15003 | BlmVI |
| bleomycin | PKS-NRPS | bleomycin_C08_condensation | condensation | AAG02364 | BGC0000963 | 11048953 | *Streptomyces verticillus* | ATCC 15003 | BlmIV |
| bleomycin | PKS-NRPS | bleomycin_C09_cyclization | cyclization | AAG02364 | BGC0000963 | 11048953 | *Streptomyces verticillus* | ATCC 15003 | BlmIV |
| bleomycin | PKS-NRPS | bleomycin_C10_cyclization | cyclization | AAG02364 | BGC0000963 | 11048953 | *Streptomyces verticillus* | ATCC 15003 | BlmIV |
| bleomycin | PKS-NRPS | bleomycin_C07_DCL | DCL | AAG02360 | BGC0000963 | 11048953 | *Streptomyces verticillus* | ATCC 15003 | BlmV |
| bleomycin | PKS-NRPS | bleomycin_C02_modifiedAA | modified amino acid | AAG02355 | BGC0000963 | 11048953 | *Streptomyces verticillus* | ATCC 15003 | BlmX |
| bleomycin | PKS-NRPS | bleomycin_C06_modifiedAA | modified amino acid | AAG02359 | BGC0000963 | 11048953 | *Streptomyces verticillus* | ATCC 15003 | BlmVI |
| C-1027 | PKS-NRPS | C1027_C02_LCL | LCL | AAL06678 | BGC0000965 | 12183628 | *Streptomyces globisporus* | C-1027 | SgcC5 |
| calcium-dependent antibiotic | NRPS | calciumdependent antibiotic_C05_DCL | DCL | CAB38518 | BGC0000315 | 12445768 | *Streptomyces coelicolor* | A3(2) | CdaPS1 |
| calcium-dependent antibiotic | NRPS | calciumdependent antibiotic_C08_DCL | DCL | CAB38517 | BGC0000315 | 12445768 | *Streptomyces coelicolor* | A3(2) | CdaPS2 |
| calcium-dependent antibiotic | NRPS | calciumdependent antibiotic_C12_DCL | DCL | CAD55498 | BGC0000315 | 12445768 | *Streptomyces coelicolor* | A3(2) | CdaPS3 |
| calcium-dependent antibiotic | NRPS | calciumdependent antibiotic_C04_epimerization | epimerization | CAB38518 | BGC0000315 | 12445768 | *Streptomyces coelicolor* | A3(2) | CdaPS1 |
| calcium-dependent antibiotic | NRPS | calciumdependent antibiotic_C11_epimerization | epimerization | CAB38517 | BGC0000315 | 12445768 | *Streptomyces coelicolor* | A3(2) | CdaPS2 |
| calcium-dependent antibiotic | NRPS | calciumdependent antibiotic_C02_LCL | LCL | CAB38518 | BGC0000315 | 12445768 | *Streptomyces coelicolor* | A3(2) | CdaPS1 |
| calcium-dependent antibiotic | NRPS | calciumdependent antibiotic_C03_LCL | LCL | CAB38518 | BGC0000315 | 12445768 | *Streptomyces coelicolor* | A3(2) | CdaPS1 |
| calcium-dependent antibiotic | NRPS | calciumdependent antibiotic_C06_LCL | LCL | CAB38518 | BGC0000315 | 12445768 | *Streptomyces coelicolor* | A3(2) | CdaPS1 |
| calcium-dependent antibiotic | NRPS | calciumdependent antibiotic_C07_LCL | LCL | CAB38518 | BGC0000315 | 12445768 | *Streptomyces coelicolor* | A3(2) | CdaPS1 |
| calcium-dependent antibiotic | NRPS | calciumdependent antibiotic_C09_LCL | LCL | CAB38517 | BGC0000315 | 12445768 | *Streptomyces coelicolor* | A3(2) | CdaPS2 |
| calcium-dependent antibiotic | NRPS | calciumdependent antibiotic_C10_LCL | LCL | CAB38517 | BGC0000315 | 12445768 | *Streptomyces coelicolor* | A3(2) | CdaPS2 |
| calcium-dependent antibiotic | NRPS | calciumdependent antibiotic_C13_LCL | LCL | CAD55498 | BGC0000315 | 12445768 | *Streptomyces coelicolor* | A3(2) | CdaPS3 |
| calcium-dependent antibiotic | NRPS | calciumdependent antibiotic_C01_starter | starter | CAB38518 | BGC0000315 | 12445768 | *Streptomyces coelicolor* | A3(2) | CdaPS1 |
| chloroeremomycin | NRPS | chloroeremomycin_C01_DCL | DCL | CAA11794 | na | 10716695 | *Amycolotopsis orientali* | PCZA363 | PCZA363-3 |
| chloroeremomycin | NRPS | chloroeremomycin_C02_DCL | DCL | CAA11794 | na | 10716695 | *Amycolotopsis orientali* | PCZA363 | PCZA363-3 |
| chloroeremomycin | NRPS | chloroeremomycin_C03_DCL | DCL | CAA11795 | na | 10716695 | *Amycolotopsis orientali* | PCZA363 | PCZA363-4 |
| chloroeremomycin | NRPS | chloroeremomycin_C04_DCL | DCL | CAA11795 | na | 10716695 | *Amycolotopsis orientali* | PCZA363 | PCZA363-4 |
| chloroeremomycin | NRPS | chloroeremomycin_C05_DCL | DCL | CAA11795 | na | 10716695 | *Amycolotopsis orientali* | PCZA363 | PCZA363-4 |
| chloroeremomycin | NRPS | chloroeremomycin_C06_DCL | DCL | CAA11796 | na | 10716695 | *Amycolotopsis orientali* | PCZA363 | PCZA363-5 |
| cinnabaramide | PKS-NRPS | cinnabaramide_C01_LCL | LCL | CBW54671 | BGC0000971 | 21387511 | *Streptomyces cinnabarigriseus* | JS360 | CinA |
| complestatin | NRPS | complestatin_C01_DCL | DCL | AAK81824 | BGC0000326 | 11447274 | *Streptomyces lavendulae* | | ComA |

| | | | | | | | | | |
|---|---|---|---|---|---|---|---|---|---|
| complestatin | NRPS | complestatin_C02_DCL | DCL | AAK81825 | BGC0000326 | 11447274 | *Streptomyces lavendulae* | | ComB |
| complestatin | NRPS | complestatin_C03_DCL | DCL | AAK81826 | BGC0000326 | 11447274 | *Streptomyces lavendulae* | | ComC |
| complestatin | NRPS | complestatin_C05_DCL | DCL | AAK81826 | BGC0000326 | 11447274 | *Streptomyces lavendulae* | | ComC |
| complestatin | NRPS | complestatin_C07_DCL | DCL | AAK81826 | BGC0000326 | 11447274 | *Streptomyces lavendulae* | | ComC |
| complestatin | NRPS | complestatin_C09_DCL | DCL | AAK81827 | BGC0000326 | 11447274 | *Streptomyces lavendulae* | | ComD |
| complestatin | NRPS | complestatin_C04_epimerization | epimerization | AAK81826 | BGC0000326 | 11447274 | *Streptomyces lavendulae* | | ComC |
| complestatin | NRPS | complestatin_C06_epimerization | epimerization | AAK81826 | BGC0000326 | 11447274 | *Streptomyces lavendulae* | | ComC |
| complestatin | NRPS | complestatin_C08_epimerization | epimerization | AAK81826 | BGC0000326 | 11447274 | *Streptomyces lavendulae* | | ComC |
| complestatin | NRPS | complestatin_C10_LCL | LCL | AAK81827 | BGC0000326 | 11447274 | *Streptomyces lavendulae* | | ComD |
| cyclomarin | NRPS | cyclomarin_C01_LCL | LCL | ABW00331 | BGC0000333 | 18331040 | *Salinispora arenicola* | CNS-205 | CymA |
| cyclomarin | NRPS | cyclomarin_C02_LCL | LCL | ABW00331 | BGC0000333 | 18331040 | *Salinispora arenicola* | CNS-205 | CymA |
| cyclomarin | NRPS | cyclomarin_C03_LCL | LCL | ABW00331 | BGC0000333 | 18331040 | *Salinispora arenicola* | CNS-205 | CymA |
| cyclomarin | NRPS | cyclomarin_C04_LCL | LCL | ABW00331 | BGC0000333 | 18331040 | *Salinispora arenicola* | CNS-205 | CymA |
| cyclomarin | NRPS | cyclomarin_C05_LCL | LCL | ABW00331 | BGC0000333 | 18331040 | *Salinispora arenicola* | CNS-205 | CymA |
| cyclomarin | NRPS | cyclomarin_C06_LCL | LCL | ABW00331 | BGC0000333 | 18331040 | *Salinispora arenicola* | CNS-205 | CymA |
| cyclomarin | NRPS | cyclomarin_C01_LCL | LCL | ABW00331 | BGC0000333 | 18331040 | *Salinispora arenicola* | CNS-205 | CymA |
| cyclosporin | NRPS | cyclosporin_C12_dual | dual | NA1 | BGC0000334 | 8376400 | *Tolypocladium inflatum* | NRRL8044 | na |
| cyclosporin | NRPS | cyclosporin_C01_LCL | LCL | NA1 | BGC0000334 | 8376400 | *Tolypocladium inflatum* | NRRL8044 | na |
| cyclosporin | NRPS | cyclosporin_C02_LCL | LCL | NA1 | BGC0000334 | 8376400 | *Tolypocladium inflatum* | NRRL8044 | na |
| cyclosporin | NRPS | cyclosporin_C03_LCL | LCL | NA1 | BGC0000334 | 8376400 | *Tolypocladium inflatum* | NRRL8044 | na |
| cyclosporin | NRPS | cyclosporin_C04_LCL | LCL | NA1 | BGC0000334 | 8376400 | *Tolypocladium inflatum* | NRRL8044 | na |
| cyclosporin | NRPS | cyclosporin_C05_LCL | LCL | NA1 | BGC0000334 | 8376400 | *Tolypocladium inflatum* | NRRL8044 | na |
| cyclosporin | NRPS | cyclosporin_C06_LCL | LCL | NA1 | BGC0000334 | 8376400 | *Tolypocladium inflatum* | NRRL8044 | na |
| cyclosporin | NRPS | cyclosporin_C07_LCL | LCL | NA1 | BGC0000334 | 8376400 | *Tolypocladium inflatum* | NRRL8044 | na |
| cyclosporin | NRPS | cyclosporin_C08_LCL | LCL | NA1 | BGC0000334 | 8376400 | *Tolypocladium inflatum* | NRRL8044 | na |
| cyclosporin | NRPS | cyclosporin_C09_LCL | LCL | NA1 | BGC0000334 | 8376400 | *Tolypocladium inflatum* | NRRL8044 | na |
| cyclosporin | NRPS | cyclosporin_C10_LCL | LCL | NA1 | BGC0000334 | 8376400 | *Tolypocladium inflatum* | NRRL8044 | na |
| cyclosporin | NRPS | cyclosporin_C11_LCL | LCL | NA1 | BGC0000334 | 8376400 | *Tolypocladium inflatum* | NRRL8044 | na |
| enniatin | NRPS | enniatin_C02_dual | dual | CAA79245 | BGC0000342 | 8483420 | *Fusarium scirpi* | ETH1536J5 | Esyn1 |
| enniatin | NRPS | enniatin_C01_LCL | LCL | CAA79245 | BGC0000342 | 8483420 | *Fusarium scirpi* | ETH1536J5 | Esyn1 |
| enterobactin | NRPS | enterobactin_C01_starter | starter | ADB98044 | BGC0000343 | 10688898 | *Escherichia coli* | chi7122 | EntF |
| exochelin | NRPS | exochelin_C01_DCL | DCL | AAC82549 | BGC0000351 | 9720878 | *Mycobacterium smegmatis* | MC2 155 | FxbB |
| exochelin | NRPS | exochelin_C02_DCL | DCL | AAC82549 | BGC0000351 | 9720878 | *Mycobacterium smegmatis* | MC2 155 | FxbB |
| exochelin | NRPS | exochelin_C03_DCL | DCL | AAC82550 | BGC0000351 | 9720878 | *Mycobacterium smegmatis* | MC2 155 | FxbC |
| exochelin | NRPS | exochelin_C04_DCL | DCL | AAC82550 | BGC0000351 | 9720878 | *Mycobacterium smegmatis* | MC2 155 | FxbC |
| exochelin | NRPS | exochelin_C05_LCL | LCL | AAC82550 | BGC0000351 | 9720878 | *Mycobacterium smegmatis* | MC2 155 | FxbC |
| fengycin | NRPS | fengycin_C01_DCL | DCL | NA1 | BGC0001095 | 10438779 | *Bacillus subtilis* | | na1 |
| fengycin | NRPS | fengycin_C05_DCL | DCL | NA2 | BGC0001095 | 10438779 | *Bacillus subtilis* | | na2 |
| fengycin | NRPS | fengycin_C09_DCL | DCL | NA4 | BGC0001095 | 10438779 | *Bacillus subtilis* | | na4 |
| fengycin | NRPS | fengycin_C12_DCL | DCL | NA5 | BGC0001095 | 10438779 | *Bacillus subtilis* | | na5 |
| fengycin | NRPS | fengycin_C04_epimerization | epimerization | NA1 | BGC0001095 | 10438779 | *Bacillus subtilis* | | na1 |
| fengycin | NRPS | fengycin_C08_epimerization | epimerization | NA3 | BGC0001095 | 10438779 | *Bacillus subtilis* | | na3 |
| fengycin | NRPS | fengycin_C11_epimerization | epimerization | NA4 | BGC0001095 | 10438779 | *Bacillus subtilis* | | na4 |
| fengycin | NRPS | fengycin_C14_epimerization | epimerization | NA5 | BGC0001095 | 10438779 | *Bacillus subtilis* | | na5 |
| fengycin | NRPS | fengycin_C02_LCL | LCL | NA1 | BGC0001095 | 10438779 | *Bacillus subtilis* | | na1 |
| fengycin | NRPS | fengycin_C03_LCL | LCL | NA1 | BGC0001095 | 10438779 | *Bacillus subtilis* | | na1 |

| | | | | | | | | | |
|---|---|---|---|---|---|---|---|---|---|
| fengycin | NRPS | fengycin_C07_LCL | LCL | NA3 | BGC0001095 | 10438779 | *Bacillus subtilis* | | na3 |
| fengycin | NRPS | fengycin_C10_LCL | LCL | NA4 | BGC0001095 | 10438779 | *Bacillus subtilis* | | na4 |
| fengycin | NRPS | fengycin_C13_LCL | LCL | NA5 | BGC0001095 | 10438779 | *Bacillus subtilis* | | na5 |
| fengycin | NRPS | fengycin_C06_starter | starter | NA3 | BGC0001095 | 10438779 | *Bacillus subtilis* | | na3 |
| fengycin | NRPS | fengycin_C01_DCL | DCL | NA1 | BGC0001095 | 10438779 | *Bacillus subtilis* | | na1 |
| gramicidin | NRPS | gramicidin_C02_DCL | DCL | CAA43838 | BGC0000367 | 1560782 | *Brevibacillus brevis* | ATCC 9999 | GrsB |
| gramicidin | NRPS | gramicidin_C01_epimerization | epimerization | CAA33603 | BGC0000367 | 1560782 | *Brevibacillus brevis* | ATCC 9999 | GrsA |
| gramicidin | NRPS | gramicidin_C03_LCL | LCL | CAA43838 | BGC0000367 | 1560782 | *Brevibacillus brevis* | ATCC 9999 | GrsB |
| gramicidin | NRPS | gramicidin_C04_LCL | LCL | CAA43838 | BGC0000367 | 1560782 | *Brevibacillus brevis* | ATCC 9999 | GrsB |
| gramicidin | NRPS | gramicidin_C05_LCL | LCL | CAA43838 | BGC0000367 | 1560782 | *Brevibacillus brevis* | ATCC 9999 | GrsB |
| HC-toxin | NRPS | HCtoxin_C02_dual | dual | AAA33023 | na | 1281482 | *Bipolaris zeicola* | SB111 | HTS1 |
| HC-toxin | NRPS | HCtoxin_C03_dual | dual | AAA33023 | na | 1281482 | *Bipolaris zeicola* | SB111 | HTS1 |
| HC-toxin | NRPS | HCtoxin_C04_dual | dual | AAA33023 | na | 1281482 | *Bipolaris zeicola* | SB111 | HTS1 |
| HC-toxin | NRPS | HCtoxin_C05_dual | dual | AAA33023 | na | 1281482 | *Bipolaris zeicola* | SB111 | HTS1 |
| HC-toxin | NRPS | HCtoxin_C01_epimerization | epimerization | AAA33023 | na | 1281482 | *Bipolaris zeicola* | SB111 | HTS1 |
| iturin | PKS-NRPS | iturin_C05_DCL | DCL | BAB69699 | BGC0001098 | 11591669 | *Bacillus subtilis* | RB14 | ItuB |
| iturin | PKS-NRPS | iturin_C07_DCL | DCL | BAB69699 | BGC0001098 | 11591669 | *Bacillus subtilis* | RB14 | ItuB |
| iturin | PKS-NRPS | iturin_C11_DCL | DCL | BAB69700 | BGC0001098 | 11591669 | *Bacillus subtilis* | RB14 | ItuC |
| iturin | PKS-NRPS | iturin_C04_epimerization | epimerization | BAB69699 | BGC0001098 | 11591669 | *Bacillus subtilis* | RB14 | ItuB |
| iturin | PKS-NRPS | iturin_C06_epimerization | epimerization | BAB69699 | BGC0001098 | 11591669 | *Bacillus subtilis* | RB14 | ItuB |
| iturin | PKS-NRPS | iturin_C10_epimerization | epimerization | BAB69700 | BGC0001098 | 11591669 | *Bacillus subtilis* | RB14 | ItuC |
| iturin | PKS-NRPS | iturin_C01_hybridC | hybrid C | BAB69698 | BGC0001098 | 11591669 | *Bacillus subtilis* | RB14 | ItuA |
| iturin | PKS-NRPS | iturin_C02_LCL | LCL | BAB69698 | BGC0001098 | 11591669 | *Bacillus subtilis* | RB14 | ItuA |
| iturin | PKS-NRPS | iturin_C03_LCL | LCL | BAB69698 | BGC0001098 | 11591669 | *Bacillus subtilis* | RB14 | ItuA |
| iturin | PKS-NRPS | iturin_C08_LCL | LCL | BAB69699 | BGC0001098 | 11591669 | *Bacillus subtilis* | RB14 | ItuB |
| iturin | PKS-NRPS | iturin_C09_LCL | LCL | BAB69699 | BGC0001098 | 11591669 | *Bacillus subtilis* | RB14 | ItuB |
| lichenysin | NRPS | lichenysin_C05_DCL | DCL | AAU39360 | BGC0000381 | 9864322 | *Bacillus licheniformis* | DSM 13 | LchAB |
| lichenysin | NRPS | lichenysin_C09_DCL | DCL | AAU39361 | BGC0000381 | 9864322 | *Bacillus licheniformis* | DSM 13 | LchAC |
| lichenysin | NRPS | lichenysin_C04_epimerization | epimerization | AAU39359 | BGC0000381 | 9864322 | *Bacillus licheniformis* | DSM 13 | LchAA |
| lichenysin | NRPS | lichenysin_C08_epimerization | epimerization | AAU39360 | BGC0000381 | 9864322 | *Bacillus licheniformis* | DSM 13 | LchAB |
| lichenysin | NRPS | lichenysin_C02_LCL | LCL | AAU39359 | BGC0000381 | 9864322 | *Bacillus licheniformis* | DSM 13 | LchAA |
| lichenysin | NRPS | lichenysin_C03_LCL | LCL | AAU39359 | BGC0000381 | 9864322 | *Bacillus licheniformis* | DSM 13 | LchAA |
| lichenysin | NRPS | lichenysin_C06_LCL | LCL | AAU39360 | BGC0000381 | 9864322 | *Bacillus licheniformis* | DSM 13 | LchAB |
| lichenysin | NRPS | lichenysin_C07_LCL | LCL | AAU39360 | BGC0000381 | 9864322 | *Bacillus licheniformis* | DSM 13 | LchAB |
| lichenysin | NRPS | lichenysin_C01_starter | starter | AAU39359 | BGC0000381 | 9864322 | *Bacillus licheniformis* | DSM 13 | LchAA |
| microcystin | PKS-NRPS | microcystin_C05_DCL | DCL | AAF00961 | BGC0001017 | 10788786 | *Microcystis aeruginosa* | PCC7806 | McyB |
| microcystin | PKS-NRPS | microcystin_C04_epimerization | epimerization | AAF00960 | BGC0001017 | 10788786 | *Microcystis aeruginosa* | PCC7806 | McyA |
| microcystin | PKS-NRPS | microcystin_C01_hybridC | hybrid C | AAF00958 | BGC0001017 | 10788786 | *Microcystis aeruginosa* | PCC7806 | McyE |
| microcystin | PKS-NRPS | microcystin_C02_LCL | LCL | AAF00958 | BGC0001017 | 10788786 | *Microcystis aeruginosa* | PCC7806 | McyE |
| microcystin | PKS-NRPS | microcystin_C06_LCL | LCL | AAF00961 | BGC0001017 | 10788786 | *Microcystis aeruginosa* | PCC7806 | McyB |
| microcystin | PKS-NRPS | microcystin_C07_LCL | LCL | AAF00962 | BGC0001017 | 10788786 | *Microcystis aeruginosa* | PCC7806 | McyC |
| microcystin | PKS-NRPS | microcystin_C03_modifiedAA | modified amino acid | AAF00960 | BGC0001017 | 10788786 | *Microcystis aeruginosa* | PCC7806 | McyA |
| mycosubtilin | PKS-NRPS | mycosubtilin_C05_DCL | DCL | AAF08796 | BGC0001103 | 10557314 | *Bacillus subtilis* | ATCC 6633 | MycB |
| mycosubtilin | PKS-NRPS | mycosubtilin_C07_DCL | DCL | AAF08796 | BGC0001103 | 10557314 | *Bacillus subtilis* | ATCC 6633 | MycB |
| mycosubtilin | PKS-NRPS | mycosubtilin_C11_DCL | DCL | AAF08797 | BGC0001103 | 10557314 | *Bacillus subtilis* | ATCC 6633 | MycC |
| mycosubtilin | PKS-NRPS | mycosubtilin_C04_epimerization | epimerization | AAF08796 | BGC0001103 | 10557314 | *Bacillus subtilis* | ATCC 6633 | MycB |

| | | | | | | | | | |
|---|---|---|---|---|---|---|---|---|---|
| mycosubtilin | PKS-NRPS | mycosubtilin_C06_epimerization | epimerization | AAF08796 | BGC0001103 | 10557314 | *Bacillus subtilis* | ATCC 6633 | MycB |
| mycosubtilin | PKS-NRPS | mycosubtilin_C10_epimerization | epimerization | AAF08797 | BGC0001103 | 10557314 | *Bacillus subtilis* | ATCC 6633 | MycC |
| mycosubtilin | PKS-NRPS | mycosubtilin_C01_hybridC | hybrid C | AAF08795 | BGC0001103 | 10557314 | *Bacillus subtilis* | ATCC 6633 | MycA |
| mycosubtilin | PKS-NRPS | mycosubtilin_C02_LCL | LCL | AAF08795 | BGC0001103 | 10557314 | *Bacillus subtilis* | ATCC 6633 | MycA |
| mycosubtilin | PKS-NRPS | mycosubtilin_C03_LCL | LCL | AAF08795 | BGC0001103 | 10557314 | *Bacillus subtilis* | ATCC 6633 | MycA |
| mycosubtilin | PKS-NRPS | mycosubtilin_C08_LCL | LCL | AAF08796 | BGC0001103 | 10557314 | *Bacillus subtilis* | ATCC 6633 | MycB |
| mycosubtilin | PKS-NRPS | mycosubtilin_C09_LCL | LCL | AAF08796 | BGC0001103 | 10557314 | *Bacillus subtilis* | ATCC 6633 | MycB |
| mycosubtilin | PKS-NRPS | mycosubtilin_C05_DCL | DCL | AAF08796 | BGC0001103 | 10557314 | *Bacillus subtilis* | ATCC 6633 | MycB |
| nodularin | PKS-NRPS | nodularin_C01_hybridC | hybrid C | AAO64407 | na | 15528492 | *Nodularia spumigena* | NSOR10 | NdaF |
| nodularin | PKS-NRPS | nodularin_C02_LCL | LCL | AAO64407 | na | 15528492 | *Nodularia spumigena* | NSOR10 | NdaF |
| nodularin | PKS-NRPS | nodularin_C04_LCL | LCL | AAO64402 | na | 15528492 | *Nodularia spumigena* | NSOR10 | NdaB |
| nodularin | PKS-NRPS | nodularin_C03_modifiedAA | modified amino acid | AAO64403 | na | 15528492 | *Nodularia spumigena* | NSOR10 | NdaA |
| nostopeptolide | PKS-NRPS | nostopeptolide_C01_LCL | LCL | AAF15891 | BGC0001028 | 12853152 | *Nostoc sp.* | GSV224 | NosA |
| nostopeptolide | PKS-NRPS | nostopeptolide_C02_LCL | LCL | AAF15891 | BGC0001028 | 12853152 | *Nostoc sp.* | GSV224 | NosA |
| nostopeptolide | PKS-NRPS | nostopeptolide_C03_LCL | LCL | AAF15891 | BGC0001028 | 12853152 | *Nostoc sp.* | GSV224 | NosA |
| nostopeptolide | PKS-NRPS | nostopeptolide_C04_LCL | LCL | AAF15891 | BGC0001028 | 12853152 | *Nostoc sp.* | GSV224 | NosA |
| nostopeptolide | PKS-NRPS | nostopeptolide_C05_LCL | LCL | AAF17280 | BGC0001028 | 12853152 | *Nostoc sp.* | GSV224 | NosC |
| nostopeptolide | PKS-NRPS | nostopeptolide_C06_LCL | LCL | AAF17280 | BGC0001028 | 12853152 | *Nostoc sp.* | GSV224 | NosC |
| nostopeptolide | PKS-NRPS | nostopeptolide_C07_LCL | LCL | AAF17280 | BGC0001028 | 12853152 | *Nostoc sp.* | GSV224 | NosC |
| nostopeptolide | PKS-NRPS | nostopeptolide_C08_LCL | LCL | AAF17281 | BGC0001028 | 12853152 | *Nostoc sp.* | GSV224 | NosD |
| nostopeptolide | PKS-NRPS | nostopeptolide_C09_LCL | LCL | AAF17281 | BGC0001028 | 12853152 | *Nostoc sp.* | GSV224 | NosD |
| penicillin | NRPS | penicillin_C01_DCL | DCL | ABA70582 | BGC0000404 | 16713314 | *Penicillium chrysogenum* | AS-P-78 | PcbAB |
| penicillin | NRPS | penicillin_C02_LCL | LCL | ABA70582 | BGC0000404 | 16713314 | *Penicillium chrysogenum* | AS-P-78 | PcbAB |
| pristinamycin | NRPS | pristinamycin_C04_DCL | DCL | CBW45647 | na | 10449311 | *Streptomyces pristinaespiralis* | Pr11 | SnbDE |
| pristinamycin | NRPS | pristinamycin_C03_epimerization | epimerization | CBW45637 | na | 10449311 | *Streptomyces pristinaespiralis* | Pr11 | SnbC |
| pristinamycin | NRPS | pristinamycin_C02_LCL | LCL | CBW45637 | na | 10449311 | *Streptomyces pristinaespiralis* | Pr11 | SnbC |
| pristinamycin | NRPS | pristinamycin_C05_LCL | LCL | CBW45647 | na | 10449311 | *Streptomyces pristinaespiralis* | Pr11 | SnbDE |
| pristinamycin | NRPS | pristinamycin_C06_LCL | LCL | CBW45647 | na | 10449311 | *Streptomyces pristinaespiralis* | Pr11 | SnbDE |
| pristinamycin | NRPS | pristinamycin_C07_LCL | LCL | CBW45647 | na | 10449311 | *Streptomyces pristinaespiralis* | Pr11 | SnbDE |
| pristinamycin | NRPS | pristinamycin_C01_starter | starter | CBW45637 | na | 10449311 | *Streptomyces pristinaespiralis* | Pr11 | SnbC |
| pyochelin | NRPS | pyochelin_C01_cyclization | cyclization | AAC83656 | BGC0000412 | 10555976 | *Pseudomonas aeruginosa* | PAO1 | PchE |
| pyochelin | NRPS | pyochelin_C02_cyclization | cyclization | AAC83657 | BGC0000412 | 10555976 | *Pseudomonas aeruginosa* | PAO1 | PchF |
| pyoverdine | NRPS | pyoverdine_C02_DCL | DCL | AAX16297 | BGC0000413 | 15743962 | *Pseudomonas aeruginosa* | str 10-15 | PvdI |
| pyoverdine | NRPS | pyoverdine_C04_DCL | DCL | AAX16297 | BGC0000413 | 15743962 | *Pseudomonas aeruginosa* | str 10-15 | PvdI |
| pyoverdine | NRPS | pyoverdine_C01_LCL | LCL | AAX16297 | BGC0000413 | 15743962 | *Pseudomonas aeruginosa* | str 10-15 | PvdI |
| pyoverdine | NRPS | pyoverdine_C03_LCL | LCL | AAX16297 | BGC0000413 | 15743962 | *Pseudomonas aeruginosa* | str 10-15 | PvdI |
| pyoverdine | NRPS | pyoverdine_C05_LCL | LCL | AAX16296 | BGC0000413 | 15743962 | *Pseudomonas aeruginosa* | str 10-15 | PvdJ |
| pyoverdine | NRPS | pyoverdine_C06_LCL | LCL | AAX16296 | BGC0000413 | 15743962 | *Pseudomonas aeruginosa* | str 10-15 | PvdJ |
| pyoverdine | NRPS | pyoverdine_C07_LCL | LCL | AAX16295 | BGC0000413 | 15743962 | *Pseudomonas aeruginosa* | str 10-15 | PvdD |
| pyoverdine | NRPS | pyoverdine_C08_LCL | LCL | AAX16295 | BGC0000413 | 15743962 | *Pseudomonas aeruginosa* | str 10-15 | PvdD |
| pyoverdine | NRPS | pyoverdine_C02_DCL | DCL | AAX16297 | BGC0000413 | 15743962 | *Pseudomonas aeruginosa* | str 10-15 | PvdI |
| pyridomycin | PKS-NRPS | pyridomycin_C02_LCL | LCL | AEF33078 | BGC0001039 | 21454714 | *Streptomyces pyridomyceticus* | NRRL B-2517 | PyrE |
| pyridomycin | PKS-NRPS | pyridomycin_C03_LCL | LCL | AEF33080 | BGC0001039 | 21454714 | *Streptomyces pyridomyceticus* | NRRL B-2517 | PyrG |
| pyridomycin | PKS-NRPS | pyridomycin_C01_starter | starter | AEF33078 | BGC0001039 | 21454714 | *Streptomyces pyridomyceticus* | NRRL B-2517 | PyrE |
| salinosporamide | PKS-NRPS | salinosporamide_C01_LCL | LCL | ABP53498 | BGC0001041 | 19590008 | *Salinispora tropica* | CNB-440 | SalA |

| | | | | | | | | | |
|---|---|---|---|---|---|---|---|---|---|
| sporolide | PKS-NRPS | sporolide_C01_LCL | LCL | ABP55165 | BGC0000150 | 18232689 | *Salinispora tropica* | CNB-440 | SpoT10 |
| streptolydigin | PKS-NRPS | streptolydigin_C01_LCL | LCL | CBA11557 | BGC0001046 | 19875077 | *Streptomyces lydicus* | NRRL 2433 | SlgN2 |
| surfactin | NRPS | surfactin_C05_DCL | DCL | CAA49817 | BGC0000433 | 8355609 | *Bacillus subtilis* | W168 | SrfAB |
| surfactin | NRPS | surfactin_C09_DCL | DCL | CAA49818 | BGC0000433 | 8355609 | *Bacillus subtilis* | W168 | SrfAC |
| surfactin | NRPS | surfactin_C04_epimerization | epimerization | CAA49816 | BGC0000433 | 8355609 | *Bacillus subtilis* | W168 | SrfA1 |
| surfactin | NRPS | surfactin_C08_epimerization | epimerization | CAA49817 | BGC0000433 | 8355609 | *Bacillus subtilis* | W168 | SrfAB |
| surfactin | NRPS | surfactin_C02_LCL | LCL | CAA49816 | BGC0000433 | 8355609 | *Bacillus subtilis* | W168 | SrfA1 |
| surfactin | NRPS | surfactin_C03_LCL | LCL | CAA49816 | BGC0000433 | 8355609 | *Bacillus subtilis* | W168 | SrfA1 |
| surfactin | NRPS | surfactin_C06_LCL | LCL | CAA49817 | BGC0000433 | 8355609 | *Bacillus subtilis* | W168 | SrfAB |
| surfactin | NRPS | surfactin_C07_LCL | LCL | CAA49817 | BGC0000433 | 8355609 | *Bacillus subtilis* | W168 | SrfAB |
| surfactin | NRPS | surfactin_C01_starter | starter | CAA49816 | BGC0000433 | 8355609 | *Bacillus subtilis* | W168 | SrfA1 |
| syringomycin | NRPS | syringomycin_C02_dual | dual | AAC80285 | BGC0000437 | 9830033 | *Pseudomonas syringae pv. syringae* | | SyrE |
| syringomycin | NRPS | syringomycin_C03_dual | dual | AAC80285 | BGC0000437 | 9830033 | *Pseudomonas syringae pv. syringae* | | SyrE |
| syringomycin | NRPS | syringomycin_C04_dual | dual | AAC80285 | BGC0000437 | 9830033 | *Pseudomonas syringae pv. syringae* | | SyrE |
| syringomycin | NRPS | syringomycin_C08_dual | dual | AAC80285 | BGC0000437 | 9830033 | *Pseudomonas syringae pv. syringae* | | SyrE |
| syringomycin | NRPS | syringomycin_C05_LCL | LCL | AAC80285 | BGC0000437 | 9830033 | *Pseudomonas syringae pv. syringae* | | SyrE |
| syringomycin | NRPS | syringomycin_C06_LCL | LCL | AAC80285 | BGC0000437 | 9830033 | *Pseudomonas syringae pv. syringae* | | SyrE |
| syringomycin | NRPS | syringomycin_C07_LCL | LCL | AAC80285 | BGC0000437 | 9830033 | *Pseudomonas syringae pv. syringae* | | SyrE |
| syringomycin | NRPS | syringomycin_C09_LCL | LCL | AAC80285 | BGC0000437 | 9830033 | *Pseudomonas syringae pv. syringae* | | SyrE |
| syringomycin | NRPS | syringomycin_C01_starter | starter | AAC80285 | BGC0000437 | 9830033 | *Pseudomonas syringae pv. syringae* | | SyrE |
| teicoplanin | NRPS | teicoplanin_C01_DCL | DCL | CAG15009 | BGC0000441 | 15113000 | *Actinoplanes teichomyceticus* | | TeiA |
| teicoplanin | NRPS | teicoplanin_C02_DCL | DCL | CAG15010 | BGC0000441 | 15113000 | *Actinoplanes teichomyceticus* | | TeiB |
| teicoplanin | NRPS | teicoplanin_C03_DCL | DCL | CAG15011 | BGC0000441 | 15113000 | *Actinoplanes teichomyceticus* | | TeiC |
| teicoplanin | NRPS | teicoplanin_C04_DCL | DCL | CAG15011 | BGC0000441 | 15113000 | *Actinoplanes teichomyceticus* | | TeiC |
| teicoplanin | NRPS | teicoplanin_C05_DCL | DCL | CAG15011 | BGC0000441 | 15113000 | *Actinoplanes teichomyceticus* | | TeiC |
| teicoplanin | NRPS | teicoplanin_C06_DCL | DCL | CAG15012 | BGC0000441 | 15113000 | *Actinoplanes teichomyceticus* | | TeiD |
| thaxtomin | NRPS | thaxtomin_C01_LCL | LCL | AAG27087 | BGC0000444 | 11115114 | *Streptomyces acidiscabies* | | TxtA |
| thaxtomin | NRPS | thaxtomin_C02_LCL | LCL | AAG27088 | BGC0000444 | 11115114 | *Streptomyces acidiscabies* | | TxtB |
| thiocoraline | NRPS | thiocoraline_C03_DCL | DCL | CAJ34374 | BGC0000445 | 16408310 | *Micromonospora sp.* | ML-1 | TioR |
| thiocoraline | NRPS | thiocoraline_C02_epimerization | epimerization | CAJ34374 | BGC0000445 | 16408310 | *Micromonospora sp.* | ML-1 | TioR |
| thiocoraline | NRPS | thiocoraline_C04_LCL | LCL | CAJ34375 | BGC0000445 | 16408310 | *Micromonospora sp.* | ML-1 | TioS |
| thiocoraline | NRPS | thiocoraline_C05_LCL | LCL | CAJ34375 | BGC0000445 | 16408310 | *Micromonospora sp.* | ML-1 | TioS |
| thiocoraline | NRPS | thiocoraline_C01_starter | starter | CAJ34374 | BGC0000445 | 16408310 | *Micromonospora sp.* | ML-1 | TioR |
| tubulysin | PKS-NRPS | tubulysin_C04_cyclization | cyclization | CAF05649 | BGC0001053 | 15324808 | *Angiococcus disciformis* | And18 | TubD |
| tubulysin | PKS-NRPS | tubulysin_C01_LCL | LCL | CAF05647 | BGC0001053 | 15324808 | *Angiococcus disciformis* | And18 | TubB |
| tubulysin | PKS-NRPS | tubulysin_C02_LCL | LCL | CAF05648 | BGC0001053 | 15324808 | *Angiococcus disciformis* | And18 | TubC |
| tubulysin | PKS-NRPS | tubulysin_C03_LCL | LCL | CAF05648 | BGC0001053 | 15324808 | *Angiococcus disciformis* | And18 | TubC |
| tubulysin | PKS-NRPS | tubulysin_C05_LCL | LCL | CAF05650 | BGC0001053 | 15324808 | *Angiococcus disciformis* | And18 | TubE |
| tyrocidine | NRPS | tyrocidine_C02_DCL | DCL | AAC45929 | BGC0000452 | 9352938 | *Brevibacillus brevis* | ATCC 8185 | TycB |

| | | | | | | | | | |
|---|---|---|---|---|---|---|---|---|---|
| tyrocidine | NRPS | tyrocidine_C06_DCL | DCL | AAC45930 | BGC0000452 | 9352938 | *Brevibacillus brevis* | ATCC 8185 | TycC |
| tyrocidine | NRPS | tyrocidine_C01_epimerization | epimerization | AAC45928 | BGC0000452 | 9352938 | *Brevibacillus brevis* | ATCC 8185 | TycA |
| tyrocidine | NRPS | tyrocidine_C05_epimerization | epimerization | AAC45929 | BGC0000452 | 9352938 | *Brevibacillus brevis* | ATCC 8185 | TycB |
| tyrocidine | NRPS | tyrocidine_C03_LCL | LCL | AAC45929 | BGC0000452 | 9352938 | *Brevibacillus brevis* | ATCC 8185 | TycB |
| tyrocidine | NRPS | tyrocidine_C04_LCL | LCL | AAC45929 | BGC0000452 | 9352938 | *Brevibacillus brevis* | ATCC 8185 | TycB |
| tyrocidine | NRPS | tyrocidine_C07_LCL | LCL | AAC45930 | BGC0000452 | 9352938 | *Brevibacillus brevis* | ATCC 8185 | TycC |
| tyrocidine | NRPS | tyrocidine_C08_LCL | LCL | AAC45930 | BGC0000452 | 9352938 | *Brevibacillus brevis* | ATCC 8185 | TycC |
| tyrocidine | NRPS | tyrocidine_C09_LCL | LCL | AAC45930 | BGC0000452 | 9352938 | *Brevibacillus brevis* | ATCC 8185 | TycC |
| tyrocidine | NRPS | tyrocidine_C10_LCL | LCL | AAC45930 | BGC0000452 | 9352938 | *Brevibacillus brevis* | ATCC 8185 | TycC |
| tyrocidine | NRPS | tyrocidine_C11_LCL | LCL | AAC45930 | BGC0000452 | 9352938 | *Brevibacillus brevis* | ATCC 8185 | TycC |
| vibriobactin | NRPS | vibriobactin_C01_starter | starter | AAF93940 | na | 12040125 | *Vibrio cholerae* | N16961 | VC0775 |
| viomycin | NRPS | viomycin_C01_DCL | DCL | AAP92496 | BGC0000458 | 12936980 | *Streptomyces vinaceus* | ATCC 11861 | VioF |
| viomycin | NRPS | viomycin_C02_LCL | LCL | AAP92491 | BGC0000458 | 12936980 | *Streptomyces vinaceus* | ATCC 11861 | VioA |
| viomycin | NRPS | viomycin_C03_LCL | LCL | AAP92491 | BGC0000458 | 12936980 | *Streptomyces vinaceus* | ATCC 11861 | VioA |
| viomycin | NRPS | viomycin_C04_LCL | LCL | AAP92499 | BGC0000458 | 12936980 | *Streptomyces vinaceus* | ATCC 11861 | VioI |
| viomycin | NRPS | viomycin_C05_LCL | LCL | AAP92503 | BGC0000458 | 12936980 | *Streptomyces vinaceus* | ATCC 11861 | VioM |
| yersiniabactin | PKS-NRPS | yersiniabactin_C01_cyclization | cyclization | AAC69587 | BGC0000467 | 9818149 | *Yersinia pestis* | KIM6 | Irp1 |
| yersiniabactin | PKS-NRPS | yersiniabactin_C02_cyclization | cyclization | AAC69587 | BGC0000467 | 9818149 | *Yersinia pestis* | KIM6 | Irp1 |
| yersiniabactin | PKS-NRPS | yersiniabactin_C03_cyclization | cyclization | AAC69588 | BGC0000467 | 9818149 | *Yersinia pestis* | KIM6 | Irp2 |

**Table S9.** Minimal inhibitory concentration (MIC) values for chitinimines I/III and II against a range of Gram-positive bacteria.

| Strain | Minimal Inhibitory Concentration (μg/mL) | |
|---|---|---|
| | **Chitinimine I/III** | **Chitinimine II** |
| *Enterococcus faecium* DSM25390 | 64 | 256 |
| *Bacillus cereus* DSM31 | 32 | 64 |
| *Bacillus subtilis* ATCC9799 | 64 | 128 |
| *Staphylococcus aureus* DSM21979 | 128 | 256 |
| *Staphylococcus aureus* RN4220 | >256 | 128 |
| *Staphylococcus aureus* ATCC6538 | 256 | 256 |
| *Staphylococcus aureus* StaAu068 | 256 | >256 |
| *Staphylococcus aureus* Sa9 | 256 | 128 |
| *Staphylococcus capitis* StaCa010 | 128 | 256 |
| *Staphylococcus haemolyticus* StaHa024 | 128 | 256 |
| *Staphylococcus hominis* StaHo017 | 256 | 256 |
| *Staphylococcus lugdunensis* StaLu018 | 128 | 256 |
| *Mycobacterium smegmatis* MC2-155 | 128 | >256 |

**Table S10.** KS proteins encoded in *pfaA* homologs used for phylogenetic analysis of the genome mining results.

| Species | Strain | Protein ID (NCBI) |
|---|---|---|
| *Shewanella pealeana* | ATCC 700345 | WP_012156130.1 |
| *Microcystis aeruginosa* | NIES-843 | WP_012265837.1 |
| *Microcystis aeruginosa* | str. Chao 1910 | WP_190357376.1 |
| *Nostoc* sp. | PCC 7524 / ATCC 29411 | WP_015137690.1 |
| *Trichormus variabilis* | NIES-23 | WP_096637125.1 |
| *Nostoc commune* | NIES-4072 | WP_109009590.1 |
| *Nostoc flagelliforme* | CCNUN1 | WP_100903912.1 |
| *Nostoc sphaeroides* | CCNUC1 | WP_152591243.1 |
| *Anabaena* sp. | YBS01 | WP_011321408.1 |
| *Scytonema hofmannii* | PCC 7110 | WP_051077101.1 |
| *Collimonas arenae* | Cal35 | WP_038484514.1 |
| *Chromobacterium sp.* | ATCC 53434 | WP_158300868.1 |
| *Paraburkholderia megapolitana* | LMG 23650 | WP_091015040.1 |
| *Chitinimonas koreensis* | DSM 17726 | WP_084300472.1 |
| *Ottowia thiooxydans* | DSM 14619 | WP_051237177.1 |
| *Tistlia consotensis* | USBA 355 | WP_085121617.1 |
| *Tahibacter aquaticus* | DSM 21667 | WP_133819585.1 |
| *Aquimarina* sp. | TRL1 | WP_176027148.1 |
| *Flavobacterium* sp. | N502540 | WP_264530534.1 |
| *Flavobacterium* sp. | F-323 | WP_230002366.1 |
| *Chitinivorax tropics* | DSM 27165 | WP_184033482.1 |
| *Paludibacterium paludis* | BCRC 80514 | WP_189533434.1 |
| *Bowmanella denitrificans* | JL63 | WP_102796139.1 |
| *Streptomyces griseus* | JCM 4516 | WP_193463672.1 |
| *Streptomyces* sp. | 43Y-GA-1 | WP_249627416.1 |
| *Streptomyces baarnensis* | NRRL B-2842 | WP_030081506.1 |
| *Streptomyces* sp. | WY228 | WP_218784432.1 |
| *Mycolicibacterium* sp. | CBMA 234 | WP_155924955.1 |
| *Mycobacterium fortunisiensis* | TNTM28 | WP_246584842.1 |
| *Mycobacterium simulans* | FB-527 | WP_260860972.1 |
| *Serratia* sp. | AS12 | WP_013814526.1 |
| *Serratia plymuthica* | S13 | WP_020439798.1 |
| | RVH1 | WP_006328030.1 |
| | Isolate 68f6912a-a76c-11e8-a962-3c4a9275d6c8 | WP_166728837.1 |
| | V4 | WP_208904410.1 |
| | 3Rp8 | WP_064800005.1 |
| | 3Re4-18 | WP_006328030.1 |
| | IV-11-34 | WP_166728837.1 |
| | MBSA-MJ1 | WP_202291812.1 |
| | A294 | WP_166728837.1 |
| | C-1 | WP_252978576.1 |
| | FDAARGOS_907 | WP_232246046.1 |
| | FDAARGOS_889 | WP_197912311.1 |
| | FDAARGOS_896 | WP_232246922.1 |
| | B37/06 | WP_241922251.1 |
| *Dickeya solani* | GBBC 2040 | WP_223849466.1 |
| | RNS 05.1.2A | WP_057083446.1 |
| *Dickeya* sp. | NCPPB 3274 | WP_238556095.1 |
| | Secpp 1600 | WP_255412271.1 |
| *Dickeya zeae* | EC1 | WP_237712637.1 |

| | | |
|---|---|---|
| *Dickeya fangzhongdai* | ND14b | WP_240476029.1 |
| | PA1 | WP_236883942.1 |
| | DSM 101947 | WP_225623144.1 |
| | ZXC1 | WP_276196108.1 |
| | 908C | WP_245167343.1 |
| | S1 | WP_242449541.1 |
| | B16 | WP_231348849.1 |
| | 643b | WP_239788825.1 |
| | AP6 | WP_241043386.1 |
| *Dickeya dadantii* | A622-S1-A17 | WP_226055486.1 |
| | S3-1 | WP_245000879.1 |
| | FZ06 | WP_263065003.1 |
| *Dickeya oryzae* | ZYY5 | WP_268906959.1 |
| *Xenorhabdus hominickii* | ANU1 | WP_084022942.1 |
| *Xenorhabdus budapestensis* | C-7-2 | WP_209028459.1 |
| *Xenorhabdus innexi* | HGB1681 | WP_086953270.1 |
| *Xenorhabdus szentirmaii* | DSM 16338 | WP_084616193.1 |

**Table S11.** KS-CLF heterodimers encoded in *pfaC* homologs used for phylogenetic analyses of the genome mining results.

| Species | Strain | Protein ID (NCBI) |
|---|---|---|
| *Shewanella pealeana* | ATCC 700345 | WP_012156132.1 |
| *Nostoc* sp. | PCC 7524 / ATCC 29411 | WP_015137694.1 |
| *Collimonas arenae* | Cal35 | WP_038484511.1 |
| *Chromobacterium sp.* | ATCC 53434 | WP_158300869.1 |
| *Paraburkholderia megapolitana* | LMG 23650 | WP_091015038.1 |
| *Chitinimonas koreensis* | DSM 17726 | WP_028447101.1 |
| *Tistlia consotensis* | USBA 355 | WP_085121616.1 |
| *Tahibacter aquaticus* | DSM 21667 | WP_133819586.1 |
| *Chitinivorax tropics* | DSM 27165 | WP_184033479.1 |
| *Paludibacterium paludis* | BCRC 80514 | WP_189533432.1 |
| *Bowmanella denitrificans* | JL63 | WP_155924956.1 |
| *Streptomyces griseus* | JCM 4516 | WP_193463673.1 |
| *Streptomyces* sp. | 43Y-GA-1 | WP_249627417.1 |
| *Streptomyces baarnensis* | NRRL B-2842 | WP_030081504.1 |
| *Streptomyces* sp. | WY228 | WP_218784434.1 |
| *Mycolicibacterium* sp. | CBMA 234 | WP_155924956.1 |
| *Mycobacterium fortunisiensis* | TNTM28 | WP_246584843.1 |
| *Mycobacterium simulans* | FB-527 | WP_186241947.1 |
| *Serratia* sp. | AS12 | WP_013814525.1 |
| *Serratia plymuthica* | S13 | WP_020439797.1 |
| | RVH1 | WP_006328028.1 |
| | Isolate 68f6912a-a76c-11e8-a962-3c4a9275d6c8 | WP_006328028.1 |
| | V4 | WP_208904409.1 |
| | 3Rp8 | WP_006328028.1 |
| | 3Re4-18 | WP_064799185.1 |
| | IV-11-34 | WP_006328028.1 |
| | MBSA-MJ1 | WP_197912312.1 |
| | A294 | WP_006328028.1 |
| | C-1 | WP_252978577.1 |
| | FDAARGOS_907 | WP_197913209.1 |
| | FDAARGOS_889 | WP_197912312.1 |
| | FDAARGOS_896 | WP_197929578.1 |
| | B37/06 | WP_241921682.1 |
| *Dickeya solani* | GBBC 2040 | WP_022632850.1 |
| | RNS 05.1.2A | WP_057083445.1 |
| *Dickeya* sp. | NCPPB 3274 | WP_042859297.1 |
| | Secpp 1600 | WP_107758962.1 |
| *Dickeya zeae* | EC1 | WP_016943533.1 |
| *Dickeya fangzhongdai* | ND14b | WP_038660417.1 |
| | PA1 | WP_121479904.1 |
| | DSM 101947 | WP_100849226.1 |
| | ZXC1 | WP_276196109.1 |
| | 908C | WP_209126573.1 |
| | S1 | WP_049854996.1 |
| | B16 | WP_038918429.1 |
| | 643b | WP_239788826.1 |
| | AP6 | WP_161131253.1 |
| *Dickeya dadantii* | A622-S1-A17 | WP_226055485.1 |

| | | |
|---|---|---|
| | S3-1 | WP_216282268.1 |
| | FZ06 | WP_263065004.1 |
| *Dickeya oryzae* | ZYY5 | WP_016943533.1 |
| *Xenorhabdus hominickii* | ANU1 | WP_069315170.1 |
| *Xenorhabdus budapestensis* | C-7-2 | WP_209028460.1 |
| *Xenorhabdus innexi* | HGB1681 | WP_086953269.1 |
| *Xenorhabdus szentirmaii* | DSM 16338 | WP_038236910.1 |

**Table S12. Bacterial and fungal strains tested for susceptibility to the chitinimines and their culture conditions.** LB = lysogeny broth, BHI = brain heart infusion broth, R2A = Reasoner's 2A medium, NA = nutrient agar, YPD = Yeast extract Peptone Dextrose.

| Species | Strain | Biosafety level | Growth medium | Temperature |
|---|---|---|---|---|
| **Gram-positive bacteria** | | | | |
| *Enterococcus faecium* | DSM 25390 | 2 | LB | 37°C |
| *Staphylococcus aureus* | DSM 21979 | 2 | LB | 37°C |
| *Staphylococcus aureus* | RN4220 | 2 | LB | 37°C |
| *Staphylococcus aureus* | ATCC 6538 | 2 | LB | 37°C |
| *Staphylococcus aureus* | StaAu068 | 2 | LB | 37°C |
| *Staphylococcus aureus* | Sa9 | 2 | LB | 37°C |
| *Staphylococcus capitis* | StaCa010 | 2 | LB | 37°C |
| *Staphylococcus epidermidis* | StaEp012 | 2 | LB | 37°C |
| *Staphylococcus haemolyticus* | StaHa024 | 2 | LB | 37°C |
| *Staphylococcus hominis* | StaHo017 | 2 | LB | 37°C |
| *Staphylococcus lugdunensis* | StaLu018 | 2 | LB | 37°C |
| *Bacillus cereus* | DSM 31/ATCC 14579 | 2 | LB | 30°C |
| *Bacillus subtilis* | ATCC 9799 | 1 | LB | 30°C |
| *Listeria monocytogenes* | LMH7738 | 2 | BHI | 30°C |
| *Gordonia bronchialis* | DSM 43247 | 2 | R2A | 28°C |
| *Mycobacterium smegmatis* | MC2-155 | 1 | LB | 37°C |
| **Gram-negative bacteria** | | | | |
| *Acinetobacter baumannii* | DSM 25645 | 2 | LB | 37°C |
| *Enterobacter roggenkampii* | DSM 16690 | 2 | LB | 37°C |
| *Klebsiella pneumoniae* | DSM 103517 | 2 | LB | 28°C |
| *Burkholderia multivorans* | 11/0583 | 2 | LB | 30°C |
| *Burkholderia singularis* | LMG 28154 | 2 | LB | 28°C |
| *Salmonella newport* | C487 | 2 | LB | 37°C |
| *Salmonella enterica* | 14029 (+GFP) | 2 | LB | 37°C |
| *Salmonella typhimurium* | LT2 | 2 | LB | 37°C |
| *Salmonella enteritidis* | ATCC 13046 | 2 | LB | 37°C |
| *Salmonella heidelberg* | #10 | 2 | LB | 37°C |
| *Caballeronia udeis* | LMG 27134 | 1 | R2A | 28°C |
| *Massilia sp. Root* 335 | DSM 102448 | 1 | R2A | 28°C |
| *Massilia flava* | DSM 26639 | 1 | R2A | 28°C |
| *Paraburkholderia megapolitana* | LMG 23650 | 1 | R2A | 28°C |
| *Trinickia dinghuensis* | LMG30259 | 1 | R2A | 28°C |
| *Robbsia andropogonis* | DSM 9511 | 1 | NA | 28°C |
| **Fungi** | | | | |
| *Candida albicans* | SC5314 | 2 | YPD | 30°C |
| *Candida albicans* | DPL1007 | 2 | YPD | 30°C |
| *Candida albicans* | DSY296 | 2 | YPD | 30°C |
| *Candida auris Clade I* | MDR OS299 | 2 | YPD | 37°C |
| *Candida auris Clade I* | *reference strain* B8441 | 2 | YPD | 37°C |
| *Candida auris Clade II* | *reference strain* B11220 | 2 | YPD | 37°C |
| *Candida. glabrata* | ATCC 2001 | 2 | YPD | 37°C |

[7] M. M. Zdouc, K. Blin, N. L. L. Louwen, J. Navarro, C. Loureiro, C. D. Bader, C. B. Bailey, L. Barra, T. J. Booth, K. A. J. Bozhüyük, J. D. D. Cediel-Becerra, Z. Charlop-Powers, M. G. Chevrette, Y. H. Chooi, P. M. D’Agostino, T. de Rond, E. Del Pup, K. R. Duncan, W. Gu, N. Hanif, E. J. N. Helfrich, M. Jenner, Y. Katsuyama, A. Korenskaia, D. Krug, V. Libis, G. A. Lund, S. Mantri, K. D. Morgan, C. Owen, C.-S. Phan, B. Philmus, Z. L. Reitz, S. L. Robinson, K. S. Singh, R. Teufel, Y. Tong, F. Tugizimana, D. Ulanova, J. M. Winter, C. Aguilar, D. Y. Akiyama, S. A. A. Al-Salihi, M. Alanjary, F. Alberti, G. Aleti, S. A. Alharthi, M. Y. A. Rojo, A. A. Arishi, H. E. Augustijn, N. E. Avalon, J. A. Avelar-Rivas, K. K. Axt, H. B. Barbieri, J. C. J. Barbosa, L. G. Barboza Segato, S. E. Barrett, M. Baunach, C. Beemelmanns, D. Beqaj, T. Berger, J. Bernaldo-Agüero, S. M. Bettenbühl, V. A. Bielinski, F. Biermann, R. M. Borges, R. Borriss, M. Breitenbach, K. M. Bretscher, M. W. Brigham, L. Buedenbender, B. W. Bulcock, C. Cano-Prieto, J. Capela, V. J. Carrion, R. S. Carter, R. Castelo-Branco, G. Castro-Falcón, F. O. Chagas, E. Charria-Girón, A. A. Chaudhri, V. Chaudhry, H. Choi, Y. Choi, R. Choupannejad, J. Chromy, M. S. C. Donahey, J. Collemare, J. A. Connolly, K. E. Creamer, M. Crüsemann, A. A. Cruz, A. Cumsille, J.-F. Dallery, L. C. Damas-Ramos, T. Damiani, M. de Kruijff, B. D. Martín, G. D. Sala, J. Dillen, D. T. Doering, S. R. Dommaraju, S. Durusu, S. Egbert, M. Ellerhorst, B. Faussurier, A. Fetter, M. Feuermann, D. P. Fewer, J. Foldi, A. Frediansyah, E. A. Garza, A. Gavriilidou, A. Gentile, J. Gerke, H. Gerstmans, J. P. Gomez-Escribano, L. A. González-Salazar, N. E. Grayson, C. Greco, J. E. G. Gomez, S. Guerra, S. G. Flores, A. Gurevich, K. Gutiérrez-García, L. Hart, K. Haslinger, B. He, T. Hebra, J. L. Hemmann, H. Hindra, L. Höing, D. C. Holland, J. E. Holme, T. Horch, P. Hrab, J. Hu, T.-H. Huynh, J.-Y. Hwang, R. Iacovelli, D. Iftime, M. Iorio, S. Jayachandran, E. Jeong, J. Jing, J. J. Jung, Y. Kakumu, E. Kalkreuter, K. B. Kang, S. Kang, W. Kim, G. J. Kim, H. Kim, H. U. Kim, M. Klapper, R. A. Koetsier, C. Kollten, Á. T. Kovács, Y. Kriukova, N. Kubach, A. M. Kunjapur, A. K. Kushnareva, A. Kust, J. Lamber, M. Larralde, N. J. Larsen, A. P. Launay, N.-T.-H. Le, S. Lebeer, B. T. Lee, K. Lee, K. L. Lev, S.-M. Li, Y.-X. Li, C. Licona-Cassani, A. Lien, J. Liu, J. A. V. Lopez, N. V. Machushynets, M. I. Macias, T. Mahmud, M. Maleckis, A. M. Martinez-Martinez, Y. Mast, M. F. Maximo, C. M. McBride, R. M. McLellan, K. M. Bhatt, C. Melkonian, A. Merrild, M. Metsä-Ketelä, D. A. Mitchell, A. V. Müller, G.-S. Nguyen, H. T. Nguyen, T. H. J. Niedermeyer, J. H. O’Hare, A. Ossowicki, B. O. Ostash, H. Otani, L. Padva, S. Paliyal, X. Pan, M. Panghal, D. S. Parade, J. Park, J. Parra, M. P. Rubio, H. T. Pham, S. J. Pidot, J. Piel, B. Pourmohsenin, M. Rakhmanov, S. Ramesh, M. H. Rasmussen, A. Rego, R. Reher, A. J. Rice, A. Rigolet, A. Romero-Otero, L. R. Rosas-Becerra, P. Y. Rosiles, A. Rutz, B. Ryu, L.-A. Sahadeo, M. Saldanha, L. Salvi, E. Sánchez-Carvajal, C. Santos-Medellin, N. Sbaraini, S. M. Schoellhorn, C. Schumm, L. Sehnal, N. Selem, A. D. Shah, T. K. Shishido, S. Sieber, V. Silviani, G. Singh, H. Singh, N. Sokolova, E. C. Sonnenschein, M. Sosio, S. T. Sowa, K. Steffen, E. Stegmann, A. B. Streiff, A. Strüder, F. Surup, T. Svenningsen, D. Sweeney, J. Szenei, A. Tagirdzhanov, B. Tan, M. J. Tarnowski, B. R. Terlouw, T. Rey, N. U. Thome, L. R. Torres Ortega, T. Tørring, M. Trindade, A. W. Truman, M. Tvilum, D. W. Udwary, C. Ulbricht, L. Vader, G. P. van Wezel, M. Walmsley, R. Warnasinghe, H. G. Weddeling, A. N. M. Weir, K. Williams, S. E. Williams, T. E. Witte, S. M. W. Rocca, K. Yamada, D. Yang, D. Yang, J. Yu, Z. Zhou, N. Ziemert, L. Zimmer, A. Zimmermann, C. Zimmermann, J. J. J. van der Hooft, R. G. Linington, T. Weber, M. H. Medema, “MIBiG 4.0: advancing biosynthetic gene cluster curation through global collaboration” *Nucleic Acids Research* **2025**, *53*, D678–D690.

### Acknowledgements

This work was supported by the Research Foundation-Flanders (FWO) through a PhD fellowship to FEAB (11PP324N). MMZ was supported by the Dutch Research Council (NWO) Grant KICH1.LWV04.21.013.

### Author Contributions

JM and ELCdlS initially identified the chitinimine BGC via a preliminary bioinformatic search. MHM, MA, MMZ and FEAB carried out the genome mining. Bioinformatic analysis of clusters resulting from the genome mining search was carried out by FEAB and JM. In-depth bioinformatic and phylogenetic analyses of the chitinimines BGC was done by FEAB and JM. HG generated the plasmid construct that would allow insertional mutagenesis of the chitinimine BGC. FEAB generated the *ΔchtnA* mutant via conjugation and performed the comparative metabolic profiling and UHPL-ESI-Q-TOF-MS(/MS) analyses. FEAB, GV and LLH performed metabolite extraction and preparative HPLC purification of the chitinimines. EL elucidated the structures of the chitinimines by NMR spectroscopic analysis. DDR and FEAB determined the absolute stereochemical configuration of amino acid constituents of the chitinimines using Marfey's method. DDR and FEAB set up basic and acid hydrolysis experiments of the chitinimines. Bioactivity assays against bacteria were carried out by FEAB and LLH. Antifungal and cytotoxicity tests were executed by GV, OVG, LLH and FEAB. PVD conceived the antifungal and cytotoxicity experiments. AW conceived experiments and reviewed the manuscript. JM and FEAB conceived the experiments and wrote the manuscript.
